# Micro𝕊plit: Semantic Unmixing of Fluorescent Microscopy Data

**DOI:** 10.1101/2025.02.10.637323

**Authors:** Ashesh Ashesh, Federico Carrara, Igor Zubarev, Vera Galinova, Melisande Croft, Melissa Pezzotti, Daozheng Gong, Francesca Casagrande, Elisa Colombo, Stefania Giussani, Elena Restelli, Eugenia Cammarota, Juan Manuel Battagliotti, Nikolai Klena, Moises Di Sante, Raghabendra Adhikari, Daniel Feliciano, Gaia Pigino, Elena Taverna, Oliver Harschnitz, Nicola Maghelli, Norbert Scherer, Damian Edward Dalle Nogare, Joran Deschamps, Francesco Pasqualini, Florian Jug

**Affiliations:** Fondazione Human Technopole, V.le Rita Levi-Montalcini, 20157, Milan, Italy; University of Pavia, Corso Strada Nuova, 65, 27100, Pavia, Italy; University of Chicago, 5801 S Ellis Ave, 60637, Chicago, USA; HHMI/Janelia Research Campus, 19700 Helix Drive, Ashburn, 20147, VA, USA

**Keywords:** deep learning, fluorescence microscopy, computational multiplexing, semantic channel unmixing, variational inference

## Abstract

Fluorescence microscopy, a key driver for progress in the life sciences, faces limitations due to the microscope’s optics, fluorophore chemistry, and photon exposure limits, necessitating trade-offs in imaging speed, resolution, and depth. Here, we introduce Micro𝕊plit, a computational multiplexing technique based on deep learning that allows multiple cellular structures to be imaged in a single fluorescent channel and then unmix them by computational means, allowing faster imaging and reduced photon exposure. We show that Micro𝕊plit efficiently separates up to four superimposed noisy structures into distinct denoised fluorescent image channels. Furthermore, using Variational Splitting Encoder-Decoder (VSE) networks, our approach can sample diverse predictions from a trained posterior of solutions. The diversity of these samples scales with the uncertainty in a given input, allowing us to estimate the true prediction errors by computing the variability between posterior samples. We demonstrate the robustness of Micro𝕊plit networks, which are trained for each splitting task at hand, across various datasets and noise levels and show its utility to image more, to image faster, and to improve downstream analysis. We provide Micro𝕊plit along with all associated training and evaluation datasets as open resources, enabling life scientists to immediately benefit from the potential of computational multiplexing and thus help accelerate the rate of their scientific discovery process.

## 1 Introduction

Fluorescence microscopy is an essential tool for exploring structures and dynamics within cells, tissues, and organisms. Multiplexed acquisition techniques involving multiple fluorophores, each tagged to a different cellular structure, are used to acquire multichannel image data. However, overlap in fluorophore excitation spectra imposes practical limits on the number of fluorophores that can be used in a biological sample (see Figure 1a) without problems such as cross-talk or bleed-through [1]. Even if a suitable set of fluorescent labels have been chosen, the sequential nature of multiplexed acquisitions requires multiple exposures of the sample, which reduces the photon budget available for other purposes [2, 3] and also limits the maximum sampling frequency in live-cell imaging scenarios. A limited photon budget can be addressed by reducing the photon exposure per acquisition, which in turn will lead to more noisy images that will be harder to analyze [3–5].

**Fig. 1.**
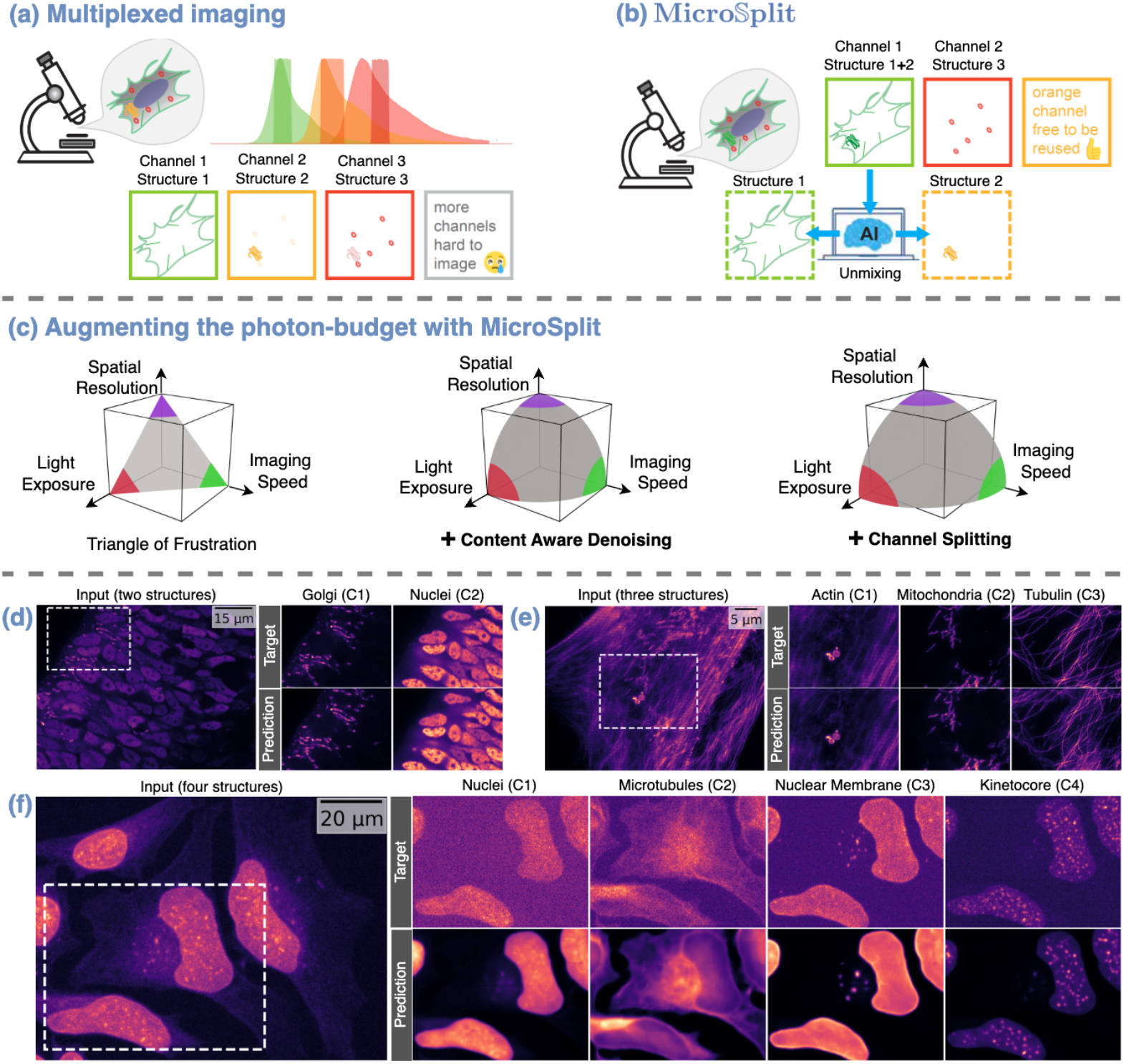
Semantic unmixing of fluorescence microscopy data with Micro𝕊plit. **(a)** Multiplexed fluorescent imaging is conducted by labeling cellular components with different fluorescent markers and imaging one after another into separate image channels. **(b)** We propose a method that allows multiple cellular structures to be imaged simultaneously in a single image channel. The superimposed structures are then split into separate channels using an adequately trained neural network. **(c)** Imaging multiple structures in one go saves precious photon budget. It is well known that the total photon budget limits the capabilities of light microscopes (left). Using content-aware denoising methods, *e*.*g*. [3, 4], can help repurpose some photon budget by acquiring more noisy raw images (middle). Imaging multiple cellular components in a single fluorescent channel saves additional photon budget that can then, for example, be invested into imaging additional structures, image more gentle, or image faster (right). **(d-f)** Exemplary qualitative results of two-channel splitting (d), three channel splitting (e), and four-channel splitting (f). Each panel shows the image channel containing multiple structures to be split (input), insets of the noisy target channels (C*i, i* ∈ {1, 2, 3, 4}) as they are used during training (target), and insets of the split channels as predictions by Micro𝕊plit (prediction). Note that the predictions are noticeable less noisy than the training data itself, which is a key feature of Micro𝕊plit’s unsupervised denoising component (see also the main text and Figure 2).

In this work, we describe a computational multiplexing approach, Micro𝕊plit, which addresses the limitations outlined above. We propose labeling and imaging multiple biological structures in a single fluorescent channel and then employing our proposed method to split superimposed structures from within this multistructure channel computationally into separate unmixed image channels (see Figure 1b). We recently presented the underlying methodological components that can enable such computational multiplexing in theory [6, 7] and have now created a practical method that (*i*) combines the benefits of both approaches into a single framework, (*ii*) enables the processing of volumetric image data by creating a highly optimized network architecture for it (see Figure 1 and Table 1), and (*iii*) we propose a procedure to assess the calibration of a trained network and the estimation of prediction errors (see Figure 2 and Section 2.3). Micro𝕊plit is designed for easy use by microscopists and life scientists, even those who are not machine learning experts.

**Table 1.**
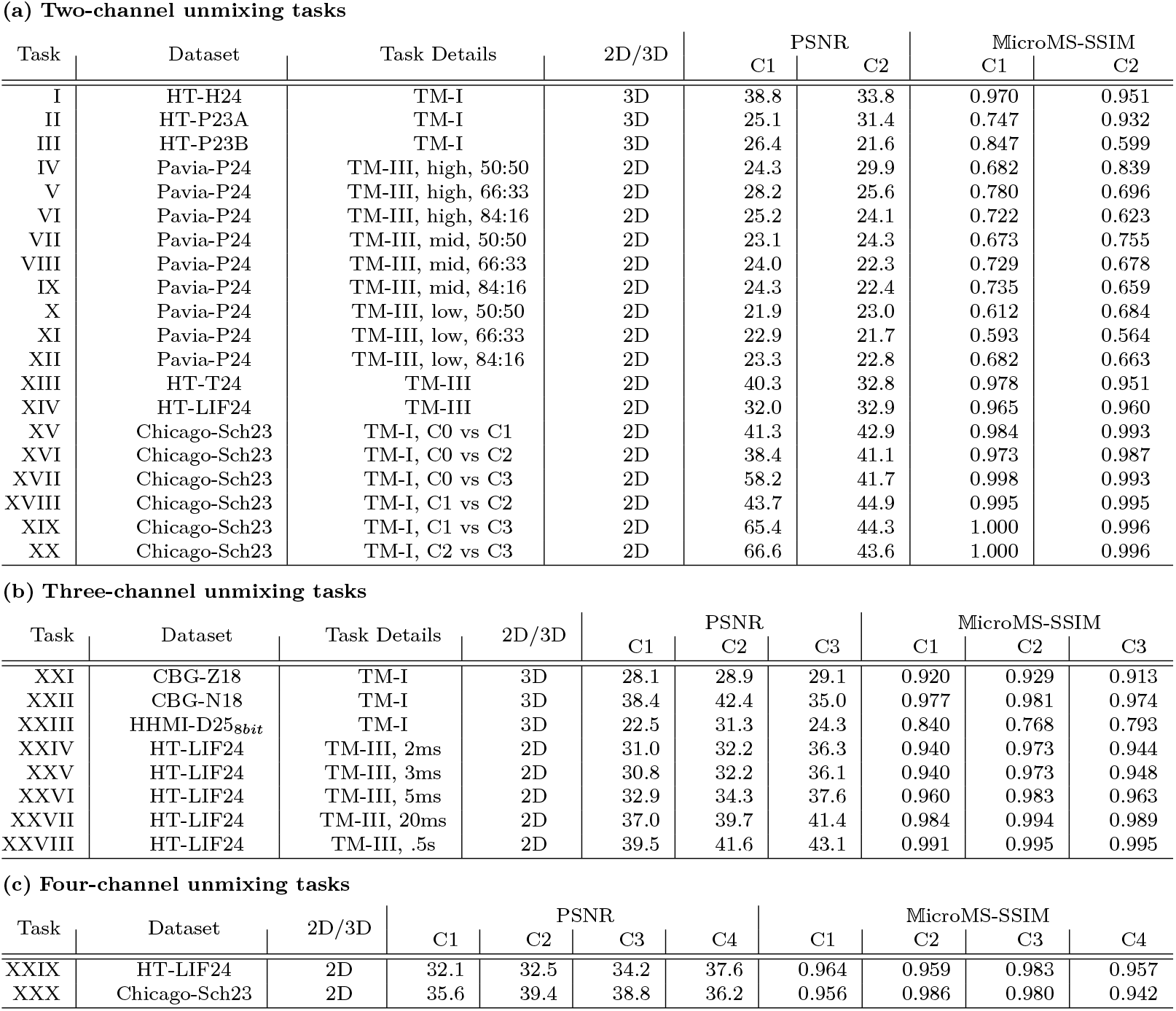
Quantitative performance on two-, three-, and four-channel splitting tasks. Throughout this work, we refer to specific splitting tasks by their *Task-Id* (first column). Some of the microscopy datasets we use (column two), give rise to multiple splitting tasks, *e*.*g*. by selecting a subset of the fluorescent channels or using different noise levels or channel mixing weights (see Section 4.1). Such *Task Details* are in abbreviated form given in the third column along with the training mode used for the task. For tasks with Task-Id XXIX and XXX, the training modes used were TM-IIIand TM-I, respectively. For volumetric tasks (labeled 3D in column *2D/3D*), we fed a 3D image stack to Micro𝕊plit, which in turn also predicts 3D outputs (posterior samples). We evaluate all the tasks on held-out test sets and report 𝕄icroMS-SSIM [8] and CARE-PSNR (PSNR) metrics for each unmixed channel prediction. In Table G.3 we list all standard errors for the results in the above table. Note that Task XXIII is the most difficult semantic unmixing task of visually different cellular structures we encountered so far, particularly w.r.t. the quality of predictions for the third channel. In Section 2.5 and Supplementary Section B.1.1, we elaborate on how low SNR and several other properties of the raw data are contributing to the simplicity or difficulty of semantic unmixing, and how potentially occurring problems can be mitigated.

| Task | Dataset | Task Details | 2D/3D | PSNR |  | MicroMS-SSIM |  |
| --- | --- | --- | --- | --- | --- | --- | --- |
|  |  |  |  | C1 | C2 | C1 | C2 |
| I | HT-H24 | TM-I | 3D | 38.8 | 33.8 | 0.970 | 0.951 |
| II | HT-P23A | TM-I | 3D | 25.1 | 31.4 | 0.747 | 0.932 |
| III | HT-P23B | TM-I | 3D | 26.4 | 21.6 | 0.847 | 0.599 |
| IV | Pavia-P24 | TM-III, high, 50:50 | 2D | 24.3 | 29.9 | 0.682 | 0.839 |
| V | Pavia-P24 | TM-III, high, 66:33 | 2D | 28.2 | 25.6 | 0.780 | 0.696 |
| VI | Pavia-P24 | TM-III, high, 84:16 | 2D | 25.2 | 24.1 | 0.722 | 0.623 |
| VII | Pavia-P24 | TM-III, mid, 50:50 | 2D | 23.1 | 24.3 | 0.673 | 0.755 |
| VIII | Pavia-P24 | TM-III, mid, 66:33 | 2D | 24.0 | 22.3 | 0.729 | 0.678 |
| IX | Pavia-P24 | TM-III, mid, 84:16 | 2D | 24.3 | 22.4 | 0.735 | 0.659 |
| X | Pavia-P24 | TM-III, low, 50:50 | 2D | 21.9 | 23.0 | 0.612 | 0.684 |
| XI | Pavia-P24 | TM-III, low, 66:33 | 2D | 22.9 | 21.7 | 0.593 | 0.564 |
| XII | Pavia-P24 | TM-III, low, 84:16 | 2D | 23.3 | 22.8 | 0.682 | 0.663 |
| XIII | HT-T24 | TM-III | 2D | 40.3 | 32.8 | 0.978 | 0.951 |
| XIV | HT-LIF24 | TM-III | 2D | 32.0 | 32.9 | 0.965 | 0.960 |
| XV | Chicago-Sch23 | TM-I, C0 vs C1 | 2D | 41.3 | 42.9 | 0.984 | 0.993 |
| XVI | Chicago-Sch23 | TM-I, C0 vs C2 | 2D | 38.4 | 41.1 | 0.973 | 0.987 |
| XVII | Chicago-Sch23 | TM-I, C0 vs C3 | 2D | 58.2 | 41.7 | 0.998 | 0.993 |
| XVIII | Chicago-Sch23 | TM-I, C1 vs C2 | 2D | 43.7 | 44.9 | 0.995 | 0.995 |
| XIX | Chicago-Sch23 | TM-I, C1 vs C3 | 2D | 65.4 | 44.3 | 1.000 | 0.996 |
| XX | Chicago-Sch23 | TM-I, C2 vs C3 | 2D | 66.6 | 43.6 | 1.000 | 0.996 |

(b) Three-channel unmixing tasks
| Task | Dataset | Task Details | 2D/3D | PSNR |  |  | MicroMS-SSIM |  |  |
| --- | --- | --- | --- | --- | --- | --- | --- | --- | --- |
|  |  |  |  | C1 | C2 | C3 | C1 | C2 | C3 |
| XXI | CBG-Z18 | TM-I | 3D | 28.1 | 28.9 | 29.1 | 0.920 | 0.929 | 0.913 |
| XXII | CBG-N18 | TM-I | 3D | 38.4 | 42.4 | 35.0 | 0.977 | 0.981 | 0.974 |
| XXIII | HHMI-D25<sub>8bit</sub> | TM-I | 3D | 22.5 | 31.3 | 24.3 | 0.840 | 0.768 | 0.793 |
| XXIV | HT-LIF24 | TM-III, 2ms | 2D | 31.0 | 32.2 | 36.3 | 0.940 | 0.973 | 0.944 |
| XXV | HT-LIF24 | TM-III, 3ms | 2D | 30.8 | 32.2 | 36.1 | 0.940 | 0.973 | 0.948 |
| XXVI | HT-LIF24 | TM-III, 5ms | 2D | 32.9 | 34.3 | 37.6 | 0.960 | 0.983 | 0.963 |
| XXVII | HT-LIF24 | TM-III, 20ms | 2D | 37.0 | 39.7 | 41.4 | 0.984 | 0.994 | 0.989 |
| XXVIII | HT-LIF24 | TM-III, .5s | 2D | 39.5 | 41.6 | 43.1 | 0.991 | 0.995 | 0.995 |

**Table 1 Quantitative performance on two-, three-, and four-channel splitting tasks.**
| Task | Dataset | 2D/3D | PSNR |  |  |  | MicroMS-SSIM |  |  |  |
| --- | --- | --- | --- | --- | --- | --- | --- | --- | --- | --- |
|  |  |  | C1 | C2 | C3 | C4 | C1 | C2 | C3 | C4 |
| XXIX | HT-LIF24 | 2D | 32.1 | 32.5 | 34.2 | 37.6 | 0.964 | 0.959 | 0.983 | 0.957 |
| XXX | Chicago-Sch23 | 2D | 35.6 | 39.4 | 38.8 | 36.2 | 0.956 | 0.986 | 0.980 | 0.942 |

**Fig. 2.**
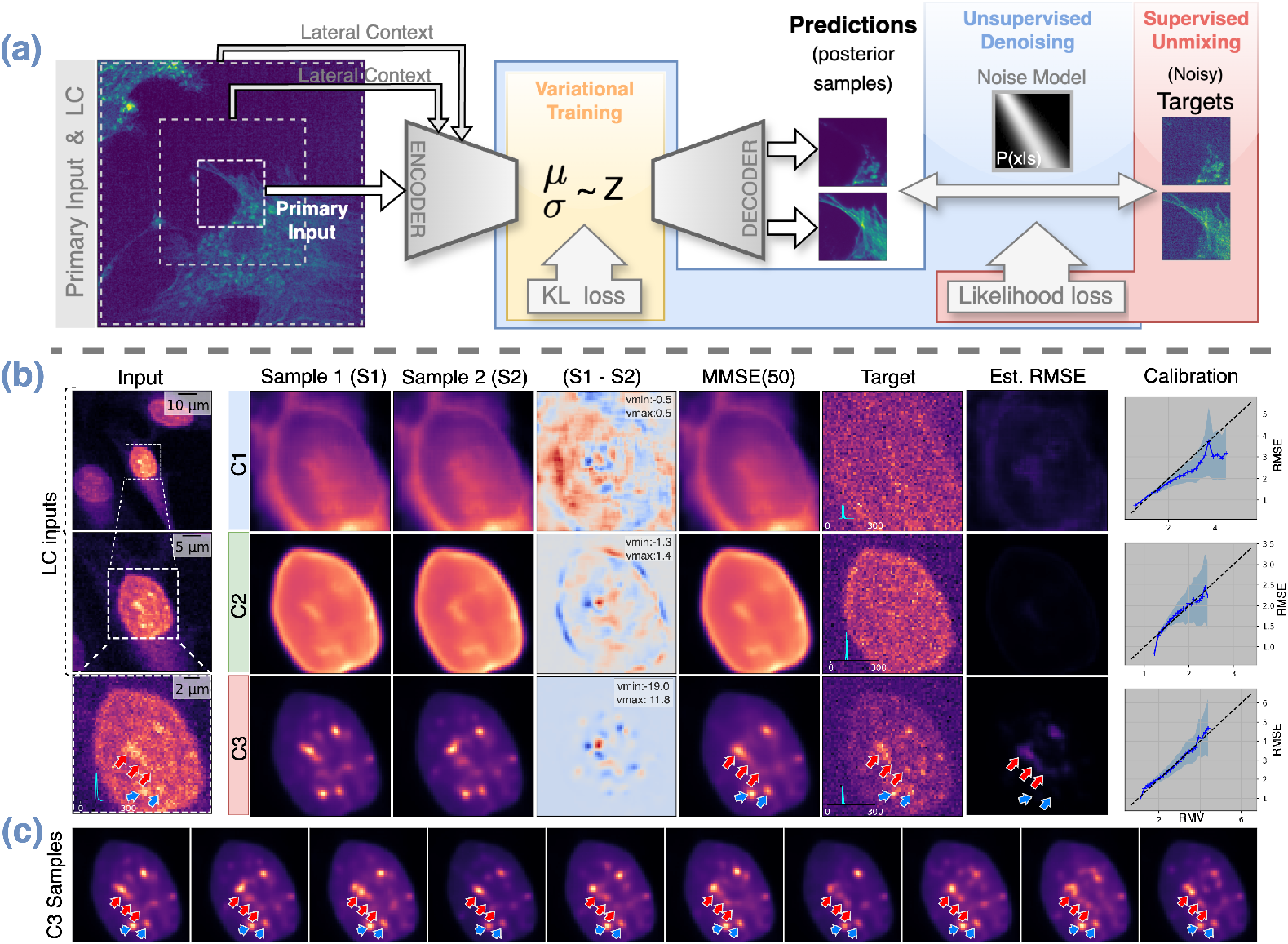
Network Architecture, Posterior Sampling, and Calibration. **(a)** Micro𝕊plit jointly learns to denoise input images (unsupervised, blue shaded area) and split superimposed structures (supervised, red shaded area). For this, we use a hierarchical network architecture, variational training (yellow shaded area), and use lateral context (LC, [6]) for better performance. Due to unsupervised denoising, using an adequate noise model [7, 9], the supervised target images can be subject to noise, and the final prediction will still be noise-free. **(b)** Leftmost column shows an input patch containing three labeled structures in a single channel at native resolution (bottom) and two LC input patches that add additional spatial context surrounding the primary input patch (towards the top). Since the input is noisy, not all structural details in the data are visible. To account for this noise-induced data uncertainty, the variational architecture we use is capable of sampling plausible “interpretations” from a learned posterior of possible solutions. For each of the three superimposed image structures (C1-C3) we show two such posterior samples in columns two and three, and their difference as a heat map in column four. The fifth column shows an approximation of the posterior minimum mean-squared-error (MMSE), computed by pixel-wise averaging 50 posterior samples. The last three columns show the noisy target images (as used during training), an estimate of the pixel-wise root mean square error (RMSE) and the calibration plots for the three channels, respectively. The calibration plots show that the uncertainty estimated solely from predictions (RMV) correlate with the true prediction error (RMSE), which means that this easy-to-calculate quantity (RMV) can be used as a surrogate for an otherwise difficult-to-assess prediction error of Micro𝕊plit. **(c)** Ten posterior samples for C3, the channel showing the highest estimated RMSE values in (b). Note that puncta at low estimated RMSE value locations (cyan arrows) remain relatively unchanged and closely resemble the structure present in the noisy target at that location, while puncta at high estimated RMSE locations (red arrows) show significant variations between samples in (c).

Using Micro𝕊plit allows users to (*i*) simultaneously image more structures than previously possible by combining up to four structures in a single channel, or to (*ii*) image the same number of structures but in fewer channels and, hence, at reduced photon exposure (see Figure 1c). The saved photon budget can then be allocated towards other objectives such as faster temporal sampling, better signal-to-noise ratio in raw acquisitions, *e*.*g*. to image more gentle and avoid phototoxicity, or any combination of these possibilities. Note that each semantic unmixing task requires training a dedicated Micro𝕊plit model. While training a universal foundation model is conceptually possible, we deliberately employ narrow, task-specific models to avoid out-of-distribution issues and to ensure robust performance on the data at hand.

Our experiments show that Micro𝕊plit is capable of separating two, three, or even four jointly imaged structures into denoised unmixed channel predictions, even if the training data are itself noisy (see Figure 1d-f). A Micro𝕊plit model learns to unmix superimposed structures by learning from examples (*i*.*e*. in a supervised way). At the same time, it co-learns to denoise the data without supervision (see Figure 2a). This means that it is sufficient to use a body of noisy training data and Micro𝕊plit will still learn to predict denoised images for each unmixed structure (images labeled “Target”^1^ in Figures 1 and 2 show such noisy training data). These properties make our method straightforward to apply in practical applications, as we show in detail in Section 2.

An additional and distinctive feature of the Variational Splitting Encoder-Decoder (VSE) networks we are using is their ability to generate multiple plausible solutions for a given input image [7]. When given inputs are unambiguous and thus can be split with high certainty, the generated solutions will closely resemble each other. In contrast, as the uncertainty in inputs increases, the diversity among the sampled solutions will reflect these inherent ambiguities by becoming more different from one another. We show that the networks we trained are *calibrated*, allowing us to estimate the otherwise hard to assess true error of predictions, even in the absence of ground truth images that are notoriously hard or even impossible to come by. Technically, this is enabled by evaluating the easy-to-compute inter-sample variability (see Section 2.3). This feature is crucial because it enables users to identify the parts of their data where Micro𝕊plit predictions may be unreliable due to ambiguities in the fed inputs. Such regions can either be disregarded or subjected to expert review, effectively addressing a persistent challenge in AI-driven bioimage analysis–the difficulty in evaluating the accuracy of predictions [1].

We show the performance of Micro𝕊plit on 24 2D and 5 3D semantic unmixing tasks, showing that we achieve accurate results in a wide range of noise levels and superimposed structures, consistently leading to high-quality denoised single-structure predictions.

In addition to the above-mentioned use cases, we propose a way to use Micro𝕊plit to remove unwanted structured image artifacts by unmixing them from the structures in which we are truly interested. We showcase this capability by removing spurious fluorescent puncta from a fluorescent microscopy dataset (see Figure 4 and Section 2.4.2).

## 2 Results

As introduced above, Micro𝕊plit enables microscopists to image multiple structures that would typically be imaged in separate fluorescence channels and instead: (*i*) capture these structures in one channel, and then unmix the superimposed structures contained in that channel by computational means. The training procedure we propose requires supervision signals (target images) for each structure to be unmixed (see Figure 1b). In the following section, we outline three practical ways to obtain suitable training data and will use them in various experimental settings to demonstrate their effectiveness.

### 2.1 Training Modes and Required Training Data

Traditionally, data are acquired using conventional multiplexing (see Figure 1a). Hence, we propose ***Training Mode I***, where previously recorded multiplexed image channels are used as targets for supervised training to achieve semantic unmixing of cellular structures (see Figure 2a). We generate mixed input images by pixel-wise summation of multiplexed image channels. These input images closely resemble what can later be acquired in a single acquisition (see Figure 1b). Although this approach yields high-quality training data, some differences in noise properties and intensity variations between structures may arise, as discussed in Section C.2. Experiments using *Training Mode I* show consistently high semantic unmixing performance, as seen in Figures 1 and 2, Table 1, and in Sections B.2 and C.2.

In cases where multiplexed data for *Training Mode I* are not available, we propose ***Training Mode II*** . Here, we assume that images of each single structure exist, but we no longer assume that all structures have been imaged in each sample. As before, here we also generate superimposed inputs by summing images showing different fluorescent structures. However, unlike before, summed-up input images no longer originate from the same sample, and any spatial correlations that might exist between these cellular structures are lost. Since a network trained with *Training Mode II* cannot leverage these correlations, we reasoned that *Training Mode I* should be at least as performant as *Training Mode II* . In Section C.2 we test this hypothesis and show that *Training Mode II* indeed comes with a slight performance drop in cases where the channels to be unmixed offer actionable spatial correlations that Micro𝕊plit can leverage.

A variation of this approach, ***Training Mode II-b***, was used to obtain the results shown in Section 2.4.2. Instead of summing uncorrelated sets of images, we used image data of superimposed structures. If those structures mix in some areas, but are visible in isolation in others, we cropped regions of interest (ROIs) showing individual structures and summed them randomly into superimposed training inputs. This training mode is best understood in the context of the results we present in Section 2.4.2.

Finally, in ***Training Mode III***, we do not create input images by summing images of individual structures but rather acquire them also at the microscope. In this mode, as in *Training Mode I*, all structures of interest must be individually labeled to allow multiplexed imaging of the required training targets. Additionally, we acquire an additional image channel by exciting all used labels at once and collecting the entirety of the emitted light, hence, directly imaging also the superimposed input directly at the microscope. Naturally, *Training Mode III* then uses this channel instead of the summation of the target channels as input to Micro𝕊plit. The advantage of this is that the input image is also subject to realistic image noise and that the relative intensity of the different structures is realistic as well (see results for tasks with IDs IV-XIV and XXIII-XXIX).

These training modes provide flexibility in the preparation of training data for Micro𝕊plit, ensuring robust performance under a variety of experimental conditions and noise levels. We provide an overview of training modes in Supplementary Table G.3.

### 2.2 Micro𝕊plit Yields High-Quality Unmixed Structures

To explore the performance of Micro𝕊plit in various biological samples, imaging modalities, and training modes (see Section 2.1), we collected a total of 10 data sets and defined a total of 30 + 6 semantic unmixing tasks, as shown in Table 1 and Table C.3. A brief overview of the datasets can be found in the Online Methods (Section 4) and more details are given in Supplementary Section F.

In Figure 1d-f we show qualitative results for three of these tasks. A quantitative assessment to the available ground truth, called training *targets* throughout this manuscript, can be found in Table 1. In the same table, we list all tasks and the achieved quality of unmixed channels in terms of CARE-PSNR (PSNR) [1] and 𝕄icroMS-SSIM [8], a variation on SSIM optimized for quantitative evaluation of microscopy image data.

Throughout all tasks, we observed average PSNR and 𝕄icroMS-SSIM values of 32.53 and 0.886, respectively, which for more common tasks, such as image denoising, would typically be considered high enough to warrant downstream processing and analysis. The lowest score of all semantic unmixing experiments still shows PSNR/𝕄icroMS-SSIM values of 21.6/0.564 (Task III-channel 2 and Task VI-channel 2, respectively), which, depending on the desired analysis to be performed, is arguably still reliable enough for downstream analysis.

However, neither PSNR nor 𝕄icroMS-SSIM are sufficient to know how trustworthy the predictions of Micro𝕊plit are, since these metrics can only be calculated when ground truth targets are available. For this reason, Micro𝕊plit employs a variational training paradigm of its Splitting Encoder-Decoder Neural Network, as shown in Figure 2. The network architecture we use is similar to a hierarchical variational autoencoder (HVAE) [10], sometimes also referred to as a Ladder-VAE [11]. The most prominent difference in our setup is that Micro𝕊plit is not an autoencoder, since the predictions are not meant to be the same as the given input. Details about the precise network architectures and the training procedure of Micro𝕊plit can be found in Section 4.3, as well as in Section A.1 and in [6, 7]. In the next section, we show how the variational nature of Micro𝕊plit can be exploited to estimate prediction errors caused by ambiguous inputs being fed.

### 2.3 Error Estimation, Data Uncertainty, and Calibration

Micro𝕊plit exploits the variational nature of its underlying architecture to enable uncertainty quantification. Since variational networks are not simple point-predictors, but instead are capable of learning an approximate posterior of possible solutions [1], Micro𝕊plit can efficiently sample such solutions. As for VAEs [12], more likely solutions will be sampled more frequently, suggesting that the analysis of the inter-sample variability can be a good surrogate for the uncertainty in the input data and, therefore, also for the trustworthiness of semantic unmixing results.

We tested this hypothesis and show our findings in Figure 2b,c. More specifically, we show an input patch and the lateral context that was fed to Micro𝕊plit, two posterior samples, the difference between them as a heatmap, and the (approximate) minimum mean squared error (MMSE) prediction, obtained by pixelwise averaging of 50 posterior samples. We also show the target patches used to train the shown 3-channel splitting task (Task XXIII). Note also here how much lower the signal-to-noise ratio is in the training data than in the predictions of a trained Micro𝕊plit network. Finally, we show the estimated pixel-wise root mean squared error (RMSE), computed using the 50 posterior samples that we also used to generate the previously shown MMSE solution.

The overall idea might be best conveyed by looking at a concrete example. Cyan and red arrows in Figure 2b,c point at image locations that might show puncta-like structures. Note that in the 10 posterior samples shown in Figure 2c, the locations pointed at with cyan arrows remain relatively unchanged and the predictions match the structure present in the target channel. However, those locations pointed at with red arrows change their appearance considerably from one posterior sample to the next. This is, in fact, also reflected in the estimated RMSE patches shown in Figure 2b, where higher estimated RMSE values appear precisely at locations where the posterior samples show elevated variability and is hence indicating that the input data is ambiguous in these regions. However, whether variability in posterior samples actually manifests itself in locations where predictions of Micro𝕊plit are prone to error and, conversely, also only in these places, is not obvious. Posterior samples could consistently predict structures in places where they are not, or consistently predict the absence of structures where, in fact, such structures should be. To show that a trained Micro𝕊plit network does not consistently hallucinate the presence or absence of structures, we propose to check how well calibrated the network is [7, 13–15].

A calibrated neural network is one whose predicted probabilities or uncertainties accurately reflect the true likelihood of outcomes. In the context of a regression task like the one executed by Micro𝕊plit, calibration means that the predicted error estimates of the model align with the actual distribution of errors in the data. We therefore estimated the true error by computing the pixel error between the MMSE prediction of Micro𝕊plit and the ground truth target images we derived from the available training data, and plotted this “true error” against the RMSE errors we described above. As the calibration plots in Figure 2b show, the true error and RMSE scale roughly linearly with respect to each other, meaning that the estimated RMSE, which can be computed from posterior samples only, is a good estimator of the true error. Hence, calibrated Micro𝕊plit networks offer a reliable and efficient way to estimate the uncertainty of the data and, therefore, also the magnitude of the error of its predictions. This property is a considerable advantage over non-variational approaches and will help future users to (*i*) filter images or image regions that lead to uncertain predictions, and (*ii*) provide evidence to themselves and others that unmixed images do not suffer from hallucinations.

### 2.4 Downstream Processing of Unmixed Data

Sample preparation and microscopy data acquisition are typically only the first steps in the pursuit of scientific insight. Once images have been acquired, their content must be analyzed and quantified. In this work, we cannot cover the full breadth of image data analyses that life scientists routinely perform on microscopy data [16]. However, in many analysis pipelines, image segmentation is a key step since it determines the identity, location, shape, and relative location of biological entities of interest. In the following section, we have quantified the segmentation performance on Micro𝕊plit predictions compared to the same analysis carried out on traditionally multiplexed image data.

#### 2.4.1 Segmentation of Unmixed Image Data is of High Quality

We show that a typical segmentation task, as it is conducted in biological research laboratories on a daily basis, leads to comparable quality results when conducted on regular multiplexed microscopy data or on unmixed images. To this end, we compared the segmentation results of three bioimage analysts who were instructed to interactively train a pixel-classifier until they reach best-possible results. We show the results in Figure 3 and Figure S7, and explain the details of this experiment below.

**Fig. 3.**
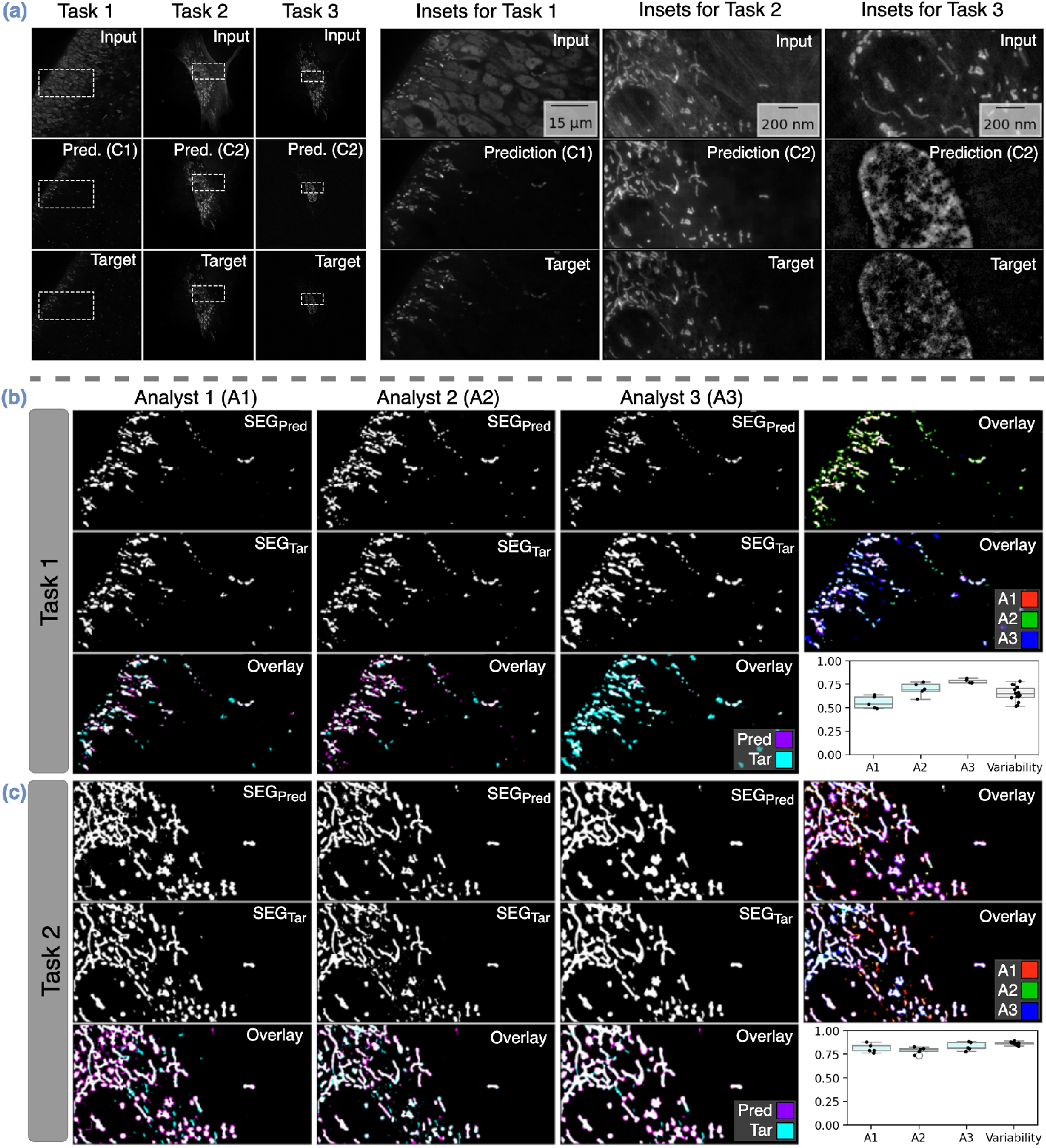
Segmentations on Unmixed Channels are in line with Inter-Observer Variability. We define three segmentation tasks, each on one unmixed channel predicted by Micro𝕊plit, and compare the segmentations created by three Bioimage Analysts with each other. **(a)** For the three tasks at hand, we show overview images (left) and insets (right) of three two-channel splitting tasks, with the superimposed raw input image in the top row, the predicted channel to be segmented in the middle row, and a single-channel control acquisitions via regular multiplexing (target, see Figure 1a) in the bottom row. **(b, c)** For Task 1 and Task 2, we show segmentation results obtained by three analysts (A1-A3) obtained on the Micro𝕊plit predictions (top row), the single-channel control acquisitions (target, middle row). The rightmost column shows overlays of the results obtained by all three analysts, with consistently segmented pixels being shown in white. The bottom row first shows the overlay of the segmentation results of a single analyst on predicted (top) vs. single channel control inputs (middle), and in the last column a box plot of all pairwise DICE scores between predicted vs. target channel inputs of the entire test set (5 images of size 1600 1600 for Task 1 and size 4096 4096 for Task 2). Note that the intra-observer variability between predicted and target segmentations (first three box-plots) are in about the same range as the inter-observer variability shown in the rightmost box-plot.

**Fig. 4.**
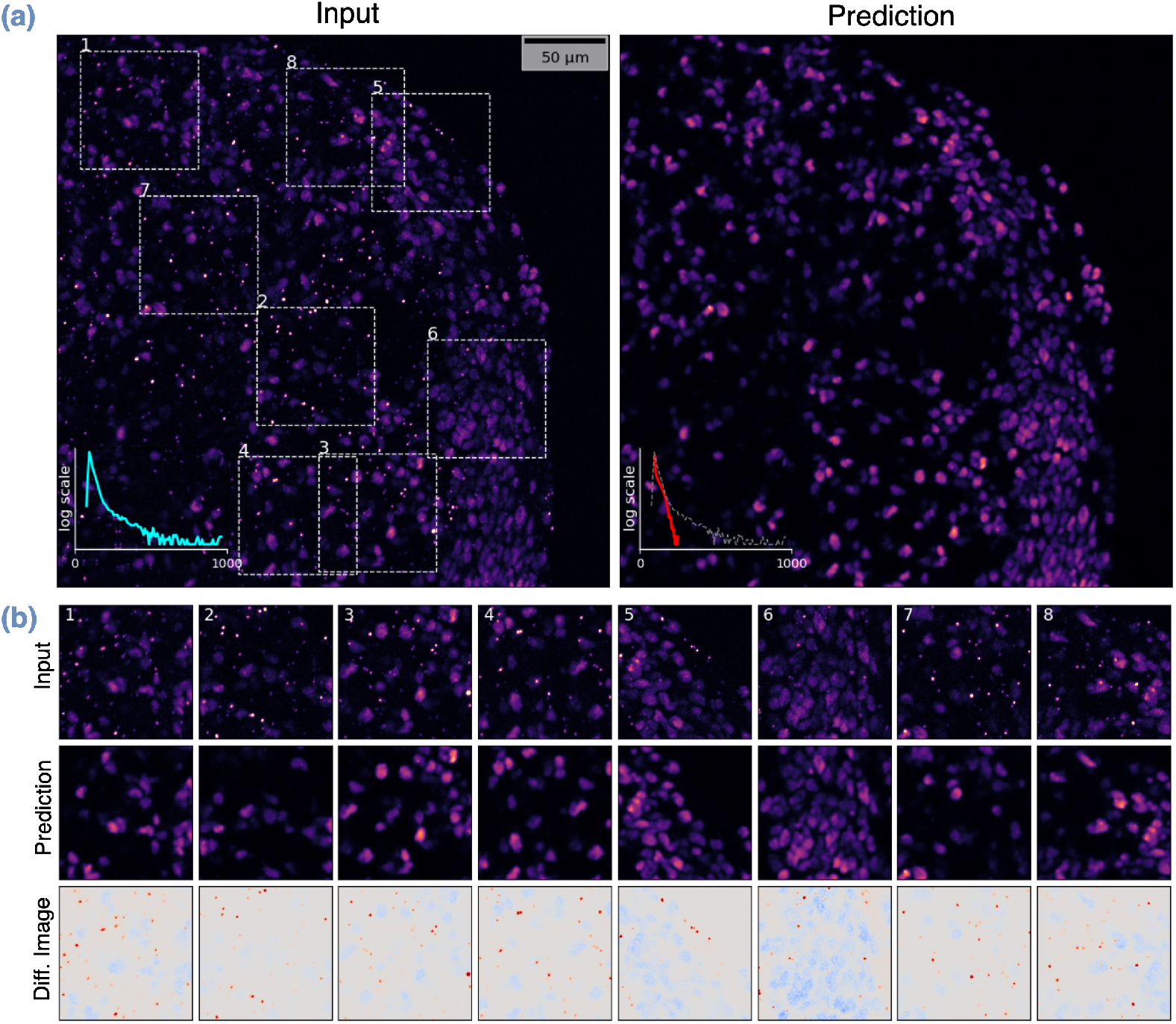
Removing Image Artifacts with Micro𝕊plit. We remove unwanted puncta present in raw microscopy data using our image unmixing approach by (*i*) selecting a set of cropped regions of interest (ROI) in the raw data that do not contain the unwanted structures (puncta, in this case), and other set of ROIs that do only contain said puncta without the fluorescent signal of interest, followed by (*ii*) creating superimposed input images by mixing (summing) randomly sampled ROIs of the two collected sets. These mixed images, together with the original pairs of ROIs, are now the input and target input used to train a Micro𝕊plit network. **(a)** An example of a raw input image (left), showing labeled nuclei and unwanted puncta, and the puncta-free prediction by Micro𝕊plit. Both images show intensity histograms in cyan and red, respectively. The dashed gray plot in the predicted image corresponds to the cyan plot of the input. The y-axes are shown on a logarithmic scale to better emphasize the puncta intensities Micro𝕊plit successfully removed. **(b)** For better visual comparison, we show input and prediction insets from (a) in the top two rows, respectively, and the difference images (input - prediction) in the bottom row (red, intensities removed; gray intensities unchanged; blue, intensities increased).

First, each analyst segmented the single-channel microscopy data acquired by regular multiplexed fluorescent microscopy (Target) and the two-channel semantic unmixing predictions of Micro𝕊plit (Prediction) without being informed about the nature of the data. We then compared the consistency between the segmentations of each analyst on the two sets of data and the consistency of the segmentations between the analysts. We observed that the variability between segmentations on target and unmixed images lies within the range of the inter-observer variability between the three analysts. We qualitatively show the segmentation results on two experiments (Tasks) in Figure 3bc and plot the measured intra-observer variabilities (A1,A2,A3) and the inter-observer variability.

Hence, in all experiments we conducted, the quality of segmentations remained unchanged when using unmixed image data compared to multiplexed images. But more interestingly, imaging two structures in a single channel can free up photon budget, which then becomes available for imaging at a higher frame rate, imaging at a higher signal-to-noise ratio, or imaging more labeled structures in additional image channels.

#### 2.4.2 Removing Unwanted Imaging Artifacts

Naturally, Micro𝕊plit predictions are just images and any subsequent analysis can be performed with them. As a second downstream processing example, we present an innovative way to remove imaging artifacts. More specifically, we imaged the post-mitotic neuronal marker CTIP2 in sliced hPSC-derived forebrain organoids using a secondary antibody conjugated to a 555 dye. The resulting image data not only had CTIP2+ nuclear staining (wanted signal), but also puncta that accumulated because of non-specific signal (*i*.*e*. undesired artifacts).

Ideally, we could use Micro𝕊plit to unmix the labeled nuclei (desired image content) and the undesired puncta (*i*.*e*. imaging artifacts). However, for this we would need training targets that contain only nuclei and others that contain only puncta, a requirement that the experimental setup of our colleagues does not permit. In Section 2.1, we introduce *Training Mode II-b*, the manual selection of image regions that show only the artifacts (puncta) or only the desired structures (the signal, in this example, nuclei). The selected image regions are then combined, just as in any *Training Mode II* experiment, and used to train a Micro𝕊plit network. Since crops without puncta or crops that show only puncta are of limited size with respect to full microscopy images (the smallest crops we collected were 100 100 pixels), lateral contextualization (LC; see Figure 2a) could not be used and was therefore disabled during training. In total, we have manually cropped 82 content regions and 48 puncta regions from the data. This led to a total of 53.7*M* pixels for training and took an analyst about eight hours of focused work.

After training had converged, we evaluated the model on previously held out full-size microscopy images. A representative result is shown in Figure 4a,b. The insets in Figure 4b allow a more detailed comparison of the raw input data containing the undesired puncta, and the predictions of the trained network. In the bottom row of the figure, we plot difference heat maps between input and prediction, which show how puncta are consistently absent in the predicted image while nuclei intensities remain mostly unaltered.

### 2.5 Limitations of Micro𝕊plit

In this study, we conduct an in-depth analysis of various factors that impact the performance of semantic unmixing. Since semantic unmixing is a new method in the toolbox of microscopists, its full positive impact and its most useful application areas will only reach community consensus in the next few years. Here, we share our experience as accurately and unfiltered as possible and recommend practical and actionable steps to improve the achievable performance.

After working with semantic unmixing for a few years, we found the following data and network properties to be the key factors affecting the performance of Micro𝕊plit: (*i*) multi-fold brightness differences (intensity skew) of the structures to be unmixed, (*ii*) the signal-to-noise ratio (SNR) of the raw data, the degree of structural (dis-)similarity between the structures to be unmixed, (*iv*) helpful and reliable spatial correlations between the structures to be unmixed, and (*v*) the size of structures to be unmixed in relation to the input patch and receptive-field size of the used network architecture. In Supplementary Section B we further discuss limitations and mitigation strategies in greater detail.

To better understand the limitations of Micro𝕊plit, we have varied the quality of image channels in two directions: signal-to-noise ratio (SNR) and relative brightness of two imaged channels (skew). More specifically, in the Pavia-P24 dataset, we have manually chosen three laser power settings per channel corresponding to a relatively high SNR, a medium SNR, and a low SNR setting. We then acquired 9 sets of training data (for all combinations of illumination laser power settings). We quantify our findings in Section B.4, where we have trained 9 Micro𝕊plit networks, one for each set of training data mentioned above, and observed that a higher degree of skew (unequal channel intensities) makes it more difficult for Micro𝕊plit to pick up the weaker of the two structures (see Table ST2).

We also analyzed the effect of skewness with the HT-P23A (Task II) and Chicago-Sch23 (Task XX) datasets. In all cases, predictions seem fit for downstream processing, although we observe blurriness in predictions for the weaker channel in a skewed HT-P23A derived task. More details can be found in Supplementary Section B.4, and in Figures S8 and S9.

To look closer at the effect of the overall signal-to-noise (SNR) of the used training data, we acquired the HT-LIF24 dataset, which also partitions into multiple subsets that are imaged at increasingly shorter exposure times (and hence at lower SNR). Again, we have trained a Micro𝕊plit network for each data subset. We ensured that the underlying content across these training sets is identical, which allowed us to do a quantitative evaluation of the performance drop as the SNR of the training data decreases. Quantitative results are shown in Table 1b, in Supplementary Section B.1, and qualitative examples are given in Figure S60 and and Figure S5. Although the results clearly show the expected trend (semantic unmixing quality decreases when using lower SNR training data), even the shortest exposure time of 2ms still leads to unmixed predictions that are fit for downstream processing and further analysis (in all cases, we measured a PSNR*>* 30.8 and 𝕄icroMS-SSIM *>* 0.94, as shown in Table 1b).

We additionally conducted experiments with synthetic Gaussian and Poisson noise on two sub-datasets from the HHMI-D25 dataset and observed that increased noise reduces the precision of fine structural details. Qualitative and quantitative results are provided in Supplementary Section B.1.1. These findings are consistent with previous observations made for denoi𝕊plit [7]. Relative to high-SNR ground truth, Micro𝕊plit tends to smooth fine structural details. Although other architectures may produce outputs that appear visually sharper, Micro𝕊plit behaves in line with the well-understood behavior of models receiving ambiguous inputs: when the input does not contain enough information to uniquely determine the underlying structure, a model can either generate smooth, “averaged” predictions that minimize expected error, or produce artificially sharp predictions that commit to one of several plausible interpretations. This trade-off is known as the perception–distortion tradeoff [17, 18], which states that beyond a certain point, improving perceptual sharpness necessarily increases reconstruction error, and vice versa.

When working with noisy input images, predictions on full-sized micrographs, which are not fitting into GPU memory in one go, are prone to show some tiling artifacts [6, 7]. Since Micro𝕊plit uses a variational network architecture and the data uncertainty in noisy inputs is higher, posterior samples (and even MMSE predictions) of neighboring tiles can have small intensity mismatches along their edges, causing said tiling artifacts. Additionally, the used network architectures are deep, having receptive field sizes well beyond the primary input patch being fed to the network. It is therefore advisable to use *inner tiling* instead of the more commonly used *outer tiling*, as previously described in [6].

The potential downstream applications of Micro𝕊plit include segmentation, tracking, detecting the presence or absence of certain structures, counting structures, and estimating dimensional properties such as radius or length of structures of interest. However, we would not recommend using Micro𝕊plit’s prediction in downstream tasks that rely on precise pixel intensities.

Finally, since Micro𝕊plit relies on distinguishable structural appearances (structural priors) for effective semantic unmixing, we performed a series of experiments to determine the sensitivity to structure similarity using the microtubule channel from the HT-LIF24 dataset. We created a series of increasingly more challenging 2-channel splitting tasks by superimposing patches from the same microtubule channel, effectively mixing structurally identical data. To provide a single parameter that can render the semantic unmixing task increasingly simpler, we decided to scale one of the two superimposed copies, resulting in structurally similar yet controllably different appearances (unscaled and scaled by factors of 1.032, 1.063, 1.125, 1.25, 1.5, and 2). The results of these experiments, see Figures S2 and S3, show that Micro𝕊plit can successfully unmix the structures starting at the scaling factor of 1.063 (see also Supplementary Section B.3 for more details). However, network training performance improves more slowly (*i*.*e*. longer training is needed) as structural similarity increases, as evidenced by the delayed inflection point and convergence behavior in Figure S3. These results underscore the importance of structural dissimilarity for effective semantic unmixing, but they also highlight the capabilities of our method handling even very similar (only slightly scaled) distributions of structures.

## 3 Discussion

In this work, we introduced Micro𝕊plit, a novel method that enables the semantic separation of multiple cellular structures superimposed in a single fluorescent image. Using Variational Splitting Encoder-Decoder (VSE) networks, we show that Micro𝕊plit achieves high-quality semantic unmixing results for up to four superimposed cellular structures, while reliably denoising the given data and additionally providing a mechanism for estimating the error in its own predictions by evaluating the localized data uncertainty in the given input. This is particularly valuable in fluorescence microscopy, where the data is typically meant to be used for scientific downstream analyses, and the soundness of unmixed channels is essential.

Arguably, the greatest utility of Micro𝕊plit lies in its ability to dramatically reduce photon exposure while still producing unmixed image channels of quality comparable to conventional multiplexed acquisitions. In Supplementary Section C.4, we quantitatively demonstrate on a representative example that Micro𝕊plit enables about a ten-fold overall reduction in emitted photos while still leading to comparable image quality. Such gains stem from two factors, *i*.*e*. (*i*) by filtering fewer photons during image acquisition, and (*ii*) via Micro𝕊plit ‘s built-in denoising capability. In this way, Micro𝕊plit effectively reduces the required photon budget, such that the additionally available photons can then be used to achieve faster imaging, higher signal-to-noise images, or the imaging of additional structures that might previously not be imaged at the same time in the same sample due to photon budget limitations. This is particularly beneficial in live-cell imaging, where minimizing phototoxicity and maximizing temporal resolution are critical for many real-world applications.

Moreover, Micro𝕊plit’s self-supervised denoising capability allows it to produce denoised predictions even when trained exclusively on noisy data. This feature is crucial for practical applications, as it reduces the need for high-quality, low-noise training data, which can be difficult or even impossible to acquire. The ability to estimate prediction errors through inter-sample variability further enhances the utility of Micro𝕊plit, providing users with a measure of uncertainty that can guide the user to exclude unreliable images or image regions from further downstream analysis.

Micro𝕊plit distinguishes itself from existing computational multiplexing techniques, such as spectral unmixing [19] or PICASSO [20], by requiring only a single superimposed image as input, rather than multiple images with different spectral overlaps. This not only simplifies the imaging process, but also allows for faster data acquisition, making Micro𝕊plit particularly useful for live-cell imaging. Furthermore, our results demonstrate that in noisy conditions, Micro𝕊plit performs better than PICASSO, highlighting Micro𝕊plit’s robustness and applicability to real-world microscopy data (see Section C.1 and Figure S4 for an in-depth comparison to PICASSO).

Lastly, Micro𝕊plit can also be used to remove imaging artifacts, and we demonstrate this in Section 2.4.2. This is always possible when artifacts and structures can, at least occasionally, seen in isolation (*i*.*e*. non-overlapping in localized regions of the superimposed image). This is a novel and powerful way to remove structured noises (*i*.*e*. image artifacts) and is a truly unique feature that other content-aware denoising methods are not capable of [1].

Although Micro𝕊plit offers all the above-mentioned advantages, it naturally also has limitations users must be informed about. As elaborated on, in detail, in Section 2.5, the performance of the model can degrade when the signal-to-noise ratio (SNR) of the input data becomes too low, when the relative intensities of the structures to be unmixed are highly skewed, or when the structures to be unmixed are too similar to each other.

While the experiments we presented give clear examples for how microscopists can benefit from Micro𝕊plit, they are only scratching the surface of what we believe microscopists and life scientists will be using Micro𝕊plit and its future derivations for. Not only will currently infeasible experimental setups become possible^2^, existing imaging protocols can be rendered more light-efficient, faster, and cost-effective. We predict that also beyond fluorescence microscopy, image unmixing will find applications, for example, in various biomedical imaging modalities. Future work could additionally explore the integration of Micro𝕊plit with adaptive imaging strategies, where the imaging parameters (*e*.*g*. exposure time or laser power) are dynamically adjusted based on the uncertainty estimates provided by the model.

To make experimentation with Micro𝕊plit easy and to reduce the entry barrier as much as possible, we have developed a comprehensive open source library^3^ and user-friendly example notebooks4 for all experiments we conducted. To this end, we have made all training and evaluation data publicly available5. Hence, our work is fully transparent, enables others to reproduce our results, and reuse our methods and implementations to elevate the rate of their scientific discovery process [21, 22].

In summary, Micro𝕊plit represents a significant advance in semantic unmixing for fluorescence microscopy, enabling the simultaneous imaging of multiple structures in a single channel and addressing key limitations in imaging speed, resolution, and photon budget. Its ability to provide high-quality denoised predictions of unmixed cellular structures, along with reliable uncertainty estimates, makes it a powerful and practical tool. We believe that Micro𝕊plit will find widespread use in a variety of biological imaging applications, enabling applications that are today simply not possible.

## Acknowledgements

The authors thank Talley Lambert, Lecturer on Systems Biology and Associate Director of Imaging Technology in the Core for Imaging Technology and Education (CITE) for useful discussions and invaluable feedback. This work was supported by the European Union through the Horizon Europe program (IMAGINE project, grant agreement 101094250-IMAGINE and AI4LIFE project, grant agreement 101057970-AI4LIFE) as well as core funding of Fondazione Human Technopole. We want to thank the IT and HPC team at Human Technopole and the BMBF-funded de.NBI Cloud within the German Network for Bioinformatics Infrastructure (de.NBI) (031A532B, 031A533A, 031A533B, 031A534A, 031A535A, 031A537A, 031A537B, 031A537C, 031A537D, 031A538A) for access to their compute infrastructure. We are grateful to Nereo Kalebic (Human Technopole) for providing the ferret sample and to Jussi Helppi and his team (Biomedical Services of the MPI-CBG in Dresden). The authors would like to thank Dott. Sara Barozzi from the Imaging Technological Development Unit of IFOM (Milan, Italy) for the preparation of the samples used for the generation of the HT-LIF24 data set. Finally, we would like to thank Janelia and the Howard Hughes Medical Institute for their support.

## 4 Online Methods

### 4.1 Used Microscopy Datasets

Here, we briefly describe the ten microscopy datasets we have used in this work, eight of which have been generated de novo by the authors; two others were taken from a previous publication [3]. All datasets will be publicly available, enabling full reproducibility of all experiments and results we have presented, and allowing others to directly compare their own methodological improvements to the approach we have presented.

The datasets cover a broad spectrum of noise levels (SNR), number of imaged fluorescent channels, overall intensity and relative intensity of these channels, imaging modality (*e*.*g*. confocal, spinning disk confocal, structured illumination, live, fixed, and expanded samples, 2D and 3D, and so forth).

Using these datasets, we have trained Micro𝕊plit models for a total of 36 tasks in total, covering 2, 3, and 4 channel semantic unmixing experiments (as shown throughout all figures and summarized in Tables 1 and C.3). While we give an overview below, please refer to the supplement for additional details about each dataset.

All datasets are openly available (https://github.com/CAREamics/Micro𝕊plit-reproducibility#links-to-all-dataseets-used-in-the-manuscript). We provide a short description below with a more detailed description in Section F.

#### HT-H24

The HT-H24 dataset was imaged by the Harschnitz group at Human Technopole. HT-H24 dataset contains immunofluorescent staining of SOX2 (555) and MAP2 (488) in DIV25 dorsal forebrain organoids generated from WTC-11 (UCSFi001-A) induced pluripotent stem cells (iPSCs) (UCSFi001-A). This is a 3D dataset and therefore enabled the creation of 3D semantic unmixing Task I.

#### HT-P23A and HT-P23B

The HT-P23A and HT-P23B datasets were imaged by the Pigino group at Human Technopole. These are 2-channel 3D datasets containing labeled microtubules and mitochondria. The samples have been prepared using cryo expansion microscopy [23]. In HT-P23A, the fluorescence intensity of labeled microtubules is strongly overpowered by the mitochondrial signal. In HT-P23B, we reduced the concentration of primary and secondary antibodies pertaining to microtubules to ensure a more equal signal.

#### Pavia-P24

The Pavia-P24 dataset was imaged at the Synthetic Physiology Laboratory in the University of Pavia. Here, we varied two important factors that affect the semantic unmixing performance, namely the SNR (high, mid and low) and relative strength of individual structures in the superimposed input (50/50, 66/33 and 84/16). We refer to these two factors in the Task-Detail column of Table 1. For example, the Task-Detail column value ‘mid, 66/33’ means that the acquisition was done with a medium SNR and the first channel used twice the laser power compared to the second channel (66*/*33 = 2).

#### HT-T24

The HT-T24 dataset was imaged by the Taverna group at Human Technopole. This dataset is a 3 channel confocal microscopy imaging of fixed E37 ferret brain sections. It is a 3-channel 2D dataset. The first two channels contain the SOX2 (transcription factor used to label stem cell nuclei) and Golgi marker Grasp65, and the last channel contains the superimposed image containing the above-mentioned two structures.

#### HT-LIF24

This dataset was imaged at the National Facility for Light Imaging at Human Technopole. Here, we imaged with five different exposure times which we refer to in the *Task Details* column of Table 1. For instance, the *Task Details* column value ‘5ms’ signifies that the acquisition used to train the task had an exposure duration of 5 milliseconds. As a result, the different acquisitions vary in their signal-to-noise ratio (SNR). We made sure that these acquisitions show exactly the same part of the sample, that is, for a given location in the specimen, we have 5 different acquired images, each imaged with a different exposure duration and, therefore, at a different SNR.

#### Chicago-Sch23

This is a four-channel, super-resolution (structured illumination) microscopy dataset of live human BJ fibroblast cells acquired at the Scherer Lab at the University of Chicago [24]. The four channels show: Actin (CellMask Orange), Mitochondria (MitoTracker Green), Microtubules (Tubulin Tracker Deep Red), and Nuclei (Hoechst). We create one four-channel splitting task (Task XXX) and multiple two-channel splitting tasks (Tasks XV to XX) using this dataset.

#### CBG-Z18 and CBG-N18

Besides the datasets we imaged for this work, we also used the publicly available CBG-Z18 (Liver) dataset and CBG-N18 (Zebrafish) dataset, originally introduced in [3].

#### HT-H23

The HT-H23 dataset was imaged by the Harschnitz group at Human Technopole. It imaged the post-mitotic neuronal marker CTIP2 in sliced hPSC-derived forebrain organoids using a secondary antibody conjugated to a 555 dye. The resulting image data not only shows CTIP2+ nuclear staining, but also puncta that accumulate because of non-specific signal. In general, unspecific puncta may arise due to suboptimal sample preparation, fixation issues, and/or non-specific antibody staining.

#### HHMI-D25

This is a six-color confocal 3D dataset of mouse liver tissue, imaged from a total of three mice, and acquired in the Feliciano Lab on the Janelia Research Campus, Howard Hughes Medical Institute (HHMI), Ashburn, Virginia, USA. Each image includes six fluorescence channels labeling distinct subcellular structures: mitochondria, cortical actin, peroxisomes, lysosomes, lipid droplets, and nuclei. Heterozygous PhAMexcised female mice carrying the Mito-Dendra2 transgene were used to provide genetic labeling of mitochondria, while the remaining structures were visualized via immunostaining. There are three sub-datasets within HHMI-D25, namely: HHMI-D25_8*bit*_, HHMI-D25_16*bit*_, and HHMI-D25_16*bit*,0.25_. In HHMI-D25_8*bit*_, all images are stored as 8-bit unsigned integers, while images of the HHMI-D25_16*bit*_ and HHMI-D25_16*bit*,0.25_ subsets, they are stored as 16-bit unsigned integers. The HHMI-D25_16*bit*,0.25_ subset is additionally downscaled and therefore four times smaller in terms of pixels, now only counting 512 × 512 pixels per image (while the other images are of size 2048 × 2048).

### 4.2 Evaluation Metrics

In this work, we have predominantly used the metrics CARE-PSNR [3] and 𝕄icroMS-SSIM [8], which are slight alterations of PSNR (Peak Signal-to-Noise Ratio) and SSIM (Structural Similarity Index Measure), respectively [1]. We did so, since CARE-PSNR and 𝕄icroMS-SSIM are designed to better assess the quality of fluorescence microscopy images than their classical alternatives [3, 8]. Please refer to Supplementary Section E for more details.

### 4.3 Training Details

As shown in Figure 1a, a superimposed image patch is fed to Micro𝕊plit as input. For a *k* channel semantic unmixing task (in this work, *k ∈*[2, 3, 4]), Micro𝕊plit outputs *k* predicted images, each containing one of the structures that are superimposed in the given input patch.

Micro𝕊plit combines the strengths of *µ*Split [6] and denoi𝕊plit [7] to enable efficient semantic unmixing in fluorescence microscopy. From *µ*Split, we inherit the ability to incorporate lateral image context (LC) through additional inputs, which capture additional spatial information around the primary input patch at increasingly lower resolutions (see Figure 1a and Figure S1). This allows the model to leverage the surrounding image context, typically leading to improved predictions. From denoi𝕊plit, Micro𝕊plit adopts unsupervised denoising capabilities and the ability to generate diverse predictions, which in turn enable robust uncertainty quantification in calibrated networks (see Figure 2).

When semantic unmixing 2D image data, we employ the Hierarchical VAE (HVAE) architecture very similar to the one described in [6, 7], while for volumetric 3D data, we implemented a suitable 3D version as described in Supplementary Section A.

The loss function for Micro𝕊plit is a weighted combination of the losses from *µ*Split and denoi𝕊plit.

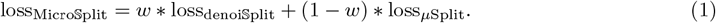

Unless explicitly specified, *w* = 0.9 is used in all experiments. This simple design also gives us the ability to switch to pure *µ*Split or denoi𝕊plit mode by simply setting *w* to 0 or 1, respectively.

For 3D data, the extra spatial dimension (Z) is averaged out before computing the KL loss, maintaining performance across volumetric inputs.

Please refer to the Supplementary Section A for more details on the used model architectures and further training details, or find our open implementation on GitHub (https://github.com/CAREamics/Micro𝕊plit-reproducibility).

## Footnotes

1 In many supervised training settings, the supervision data is referred to as “ground truth”. In this manuscript we instead refer to the supervision signals as “Target” since Micro𝕊plit can be trained exclusively on noisy data that is therefore not *ground truth*.

2 Imaging 8 or even more cellular structures on a regular microscope set up to image up to 4 fluorescent channels is one such example.

3 https://github.com/CAREamics/careamics

4 https://careamics.github.io

5 For the time being, please find all data at https://github.com/CAREamics/Micro𝕊plit-reproducibility#links-to-all-dataseets-used-in-the-manuscript.

## Notes

https://github.com/CAREamics/MicroSplit-reproducibility

### Competing Interest Statement

The authors have declared no competing interest.

### Summary of Updates

Making the EU happy by not thanking the European Commission. Small improvements on main text. Useful new supplementary material sections, most notably back-on-the-envelope computations on how much photon-budget is saved on one concrete example.

