## Supplementary Material for "Micro𝕊plit: Semantic Unmixing of Fluorescent Microscopy Data"

### A Model Architecture and Training

In Micro𝕊plit, we have combined the benefits of $\mu$Split [6] and denoi𝕊plit [7], cast those ideas in a common learning framework, enabled direct training and prediction on volumetric data, and have extensively tested and evaluated its performance on a wide range of datasets and semantic unmixing tasks. We have also made openly available all training and prediction code and all data we used. We do this to foster rapid adoption of Micro𝕊plit by the scientific community and to allow others to improve our approach and compare our results with relative ease. In this section, we will discuss in detail the aspects of the above-mentioned works that were integrated into Micro𝕊plit. Subsequently, we also describe our loss function and important training hyperparameters.

From $\mu$Split, we inherit the ability to efficiently incorporate spatial context using additional inputs, called *Laterally Contextualization* (LC) inputs [6]. Here we feed, next to the primary input, for which the prediction will be made, additional LC inputs that help the network to better understand the image context from which the primary input is taken (see Figure 2a and S1). These successive LC inputs are larger and larger patches centered on the primary input patch, but downscaled to the same pixel dimensions as the primary input itself. Hence, LC inputs capture the spatial context around the primary input patch but do so at lower resolutions to ensure efficient learning and predictions in a reasonably sized overall network.

The network architectures we proposed come in flavors that trade computational complexity and GPU consumption with the best possible prediction quality. Its most GPU-efficient variant, *Lean-LC*, can train on a single GPU using less than 5GB of GPU memory. If more resources are available, it is advisable to opt for setups such as *Deep-LC*, which show better predictive performance at increased computational cost.

In Figure S1, we have briefly described the three $\mu$Split variants. The white regions correspond to features originating from the primary input patch. As in U-Net architectures, spatial resolution halves at each successive hierarchy level through pooling operations, causing these white areas to progressively diminish in size. In $\mu$Split (and hence also in Micro𝕊plit), the pooled embeddings undergo zero-padding before being concatenated with feature maps from lateral contextualization (LC) inputs. These LC features are processed through dedicated 'Input branch' sub-networks consisting of convolutional layers with non-linear activations, dropout, and normalization components. 'Input branch' does not have pooling operations, and so their feature maps, at each hierarchy level, maintain spatial dimensions identical to the primary-input (represented by gray regions). This preservation enables merged embeddings from both pathways to retain the original spatial dimensions throughout the network hierarchy (gray squares). This is the core idea of LC Micro𝕊plit has inherited from $\mu$Split. In the caption of Figure S1 we provide more details regarding the differences in the three variants of $\mu$Split. Please refer to [6] for more details.

From denoi𝕊plit, we inherit the ability to jointly perform unsupervised denoising, using suitable *Noise Models*. This also enables Micro𝕊plit to sample diverse predictions from a learned approximate posterior that captures a notion of the data uncertainty, as demonstrated by our trained networks being calibrated (see Section 2.3). While also $\mu$Split is a variational approach that is in theory capable of generating multiple predictions from its posterior, we found that denoi𝕊plit, arguably due to its different KL-loss formulation, produces a higher diversity that is better in line with the uncertainty in the data. In Figure S1, we present the architecture of Micro𝕊plit, which has LC inputs and Noise models, all integrated into a single setup.

As mentioned above, we also enabled Micro𝕊plit to operate directly on volumetric image data, a possibility that was absent in both $\mu$Split and denoi𝕊plit.

#### A.1 Loss Function used to train Micro𝕊plit

The loss function of Micro𝕊plit is the weighted average between the $\mu$Split loss and denoi𝕊plit loss, that is,

$$\text{loss}_{\text{Micro}\mathbb{S}\text{plit}} = w * \text{loss}_{\text{denoi}\mathbb{S}\text{plit}} + (1 - w) * \text{loss}_{\mu\text{Split}} \tag{2}$$

Unless explicitly specified, $w = 0.9$ is used in all experiments we conducted. This simple design also gives us the ability to switch to pure $\mu$Split or denoi𝕊plit mode by simply setting $w$ to 0 or 1, respectively.

To incorporate LC inputs into the denoi𝕊plit setup, we observed the need to modify the KL loss formulation used in $\text{loss}_{\text{denoi}\mathbb{S}\text{plit}}$. In denoi𝕊plit, pixel-wise KL divergence is computed at every hierarchy level. Let $KL_i$ denote the pixel-wise KL divergence tensor at the $i^{th}$ hierarchy level. KL-loss component for this hierarchy level, $kl_i$ is defined as

$$\text{kl}_i = \alpha \cdot \sum_{j,h,w} \text{KL}_i[j, h, w]. \tag{3}$$

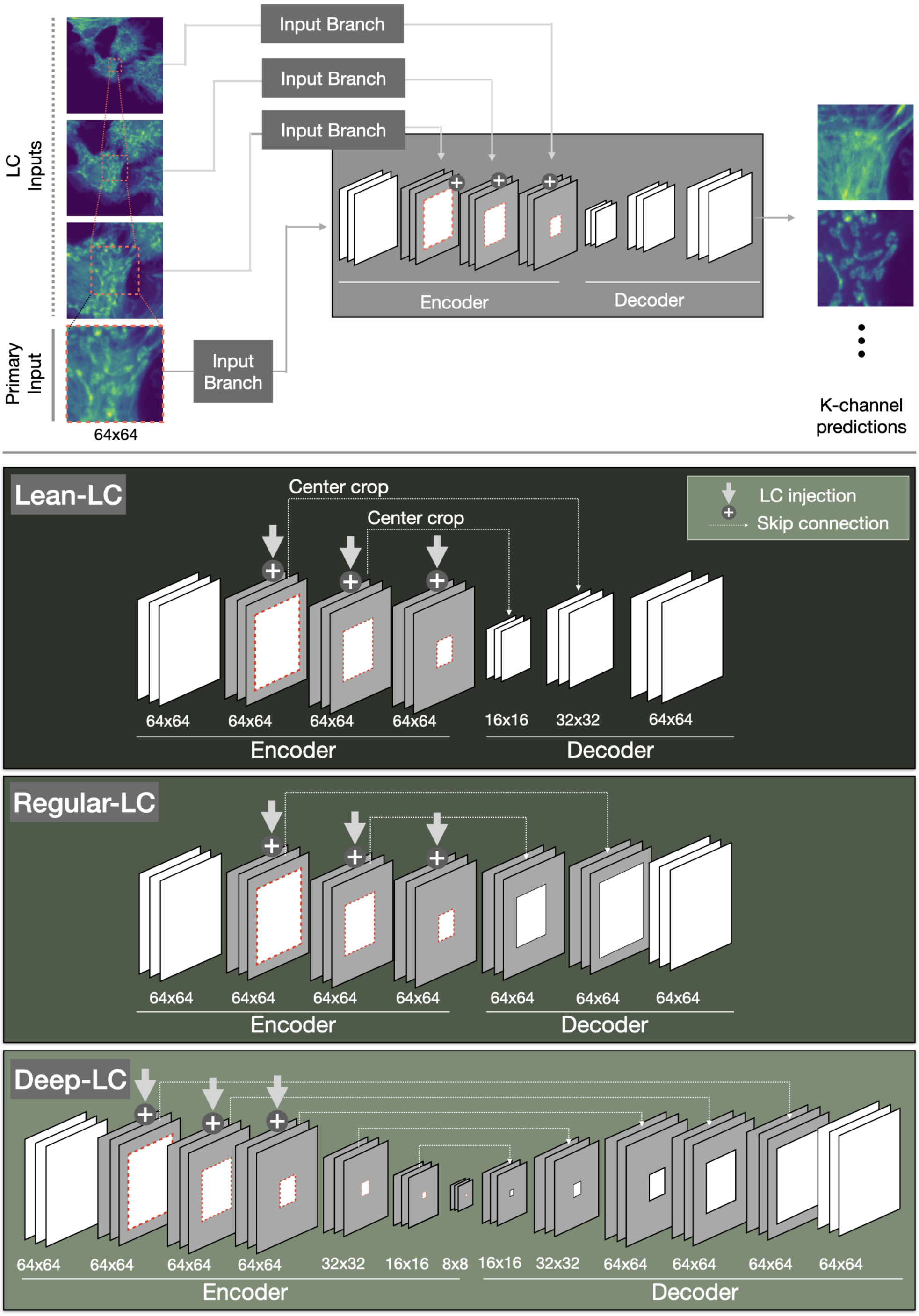


**Fig. S1 Network architectures employed by Micro𝕊plit.** This figure provides a detailed breakdown of the components shown in Figure 2a. Specifically, it illustrates the encoder-decoder structure of our architecture, which is adapted from our prior work, $\mu$Split [6]. Since this diagram focuses on the transformation of input data into predictions, it does not include the loss terms (such as the KL-divergence loss and the noise-model-based likelihood loss). As in [6], we improve the predictive capacity of Micro𝕊plit by feeding not only an image patch (primary input), but also additional image context. To this end, we introduced *lateral contextualization* (LC). These LC inputs capture a larger portion of the input data but at increasingly lower pixel-resolution. This lead to three meta-architectures, Lean-LC, Regular-LC, and Deep-LC, each with increasingly higher computational and GPU memory demands but also leading to increasingly higher unmixing performance. The gray region shown in the embeddings in all three meta-architectures represents the extra spatial size in the embeddings coming solely due to LC inputs. In Lean-LC, embeddings of different hierarchy levels in the encoder benefit from LC inputs, but the decoder does not. In Regular-LC, a more GPU-consuming variation, both the encoder's and the decoder's embeddings use LC (*i.e.* show gray regions in the figure). Finally, since in Regular-LC the spatial dimensions of the embedding layers do not decrease, it enabled us to stack some additional hierarchy levels on top of the previously used ones. We labeled the resulting architecture as Deep-LC, the most performant but also most resource heavy variation. In order to get best results, we have used Deep-LC whenever possible, but it is important to note that with just two simple hyper-parameter switches, one can instantly make use of Regular-LC or Lean-LC and train on cheaper and older consumer GPUs.

With LC inputs, the spatial dimensions of the latent space tensors and therefore $KL_i$ do not decrease and the summing operation in this formulation leads to a higher value, owing to the larger number of summands (which are all non-negative). This gives unnecessarily high weight to the KL loss with respect to the likelihood loss component. We observed that this degrades the performance. To handle this, we center-cropped $KL_i$ to the shape they would assume if there were no LC inputs. Let $KL_i^{cropped}$ denote the appropriately center-cropped version of $KL_i$. Our modified KL loss for denoiSplit becomes

$$\mathrm{kl}_i = \alpha \cdot \sum_{j,h,w} \mathrm{KL}_i^{cropped}[j, h, w]. \tag{4}$$

We encountered a very similar issue when working with volumetric data. In that case, pixel-wise KL-divergence is a 4-dimensional tensor $C \times Z \times H_i \times W_i$. As we work with larger and larger $Z$, the summation in Equation 3 would increase since the summation would be on all 4 dimensions. This again leads to giving more weight to the KL loss component against the likelihood loss, thus rendering the performance inferior. Note that this affect will become more severe when we increase the number of $Z$ frames in the input. In other words, adding more information in the input was not beneficial. To handle this, we separately took care of the extra $Z$ dimension by taking the average along this dimension. The resultant 3D tensor is then passed to Equation 4 to compute the KL loss component. Note that there is an additional dimension of *batch size* which we have not mentioned in the above explanation. That is because KL-loss is computed separately for every element in the batch.

##### A.2 Hyper-parameters used during Training

We use *PyTorch* package for creating our training and evaluation pipelines. We use a `batch_size` of 32, `max_epoch` of 400 and `learning_rate` of 0.001. We use *Adamax* optimizer and *ReduceLROnPlateau* as the learning rate scheduler with `lr_scheduler_patience` set to 150. During training, we use 16-bit precision. We use 2 LC inputs in our *Deep-LC* configuration (`multiscale_lowres_count = 3`). Please refer to our code (https://github.com/CAREamics/MicroSplit-reproducibility) for more details. We want to state that all experiments were done using code hosted at https://github.com/juglab/MicroSplit. However, we have developed https://github.com/CAREamics/MicroSplit-reproducibility with the objective of providing user-friendly code with easier adaptability to custom datasets.

#### B Analyzing Factors that Affect Predictive Performance

##### B.1 Pixel-noise (pixel-wise independent noise)

From our different experiments, we have found that there are two prominent factors that affect the performance on semantic unmixing tasks. The first factor is the amount of pixel-independent noise that is present in the superimposed input images and target images. More noise means inferior performance. To quantify the effect of noise, we imaged our HT-LIF24 dataset with exposure durations of $2ms$, $3ms$, $5ms$, $20ms$, and $500ms$. We imaged it such that the underlying content in these sub-datasets is identical. That is, for every frame in the $2ms$ acquisition, we have the corresponding higher SNR frames in $3ms$, $5ms$, $20ms$ and $500ms$ acquisitions. We trained a three-channel semantic unmixing task separately for each exposure duration sub-dataset. We made predictions on the held-out test set input frames from their respective exposure duration sub-datasets. We evaluate the prediction against the target channels present in the $500ms$ sub-dataset.

Although the results (Tasks XXIV-XXIX) in Table 1 show the expected trend (semantic unmixing quality decreases when using lower SNR training data), even the shortest exposure time of $2ms$ still lead to unmixed predictions that are fit for downstream processing and analysis (in all cases we measured a PSNR $> 30.8$ and MicroMS-SSIM $> 0.94$). To explain the performance drop, supplementary figure S60 shows that reduced PSNR primarily results from the loss of high-frequency details.

###### B.1.1 Lessons learned from the difficult Task XXIII (on HHMI-D25 data)

Out of all tasks mentioned in the Table 1, the task XXIII which uses HHMI-D25$_{8bit}$ dataset has considerable performance issues, especially in the third channel (see Supplementary Figure S57). In the next few paragraphs we will describe our approach to investigating this issue and present a working solution. We do this to provide an example for how users of MicroSplit can improve solutions that might initially not lead up to the required semantic unmixing quality.

Signal-to-noise (SNR) is one of the factors which plays a role in almost all deep-learning based methods and denoising approaches, and semantic unmixing is no exception. To investigate the role of SNR for task XXIII, we first denoised the raw data of HHMI-D25$_{8bit}$ using Noise2Void [4], and then trained Micro$\mathbb{S}$plit using those denoised images, calling this training task 'Task XXXI'. Comparing the results of tasks XXIII and XXXI, see also Table C.3, it becomes apparent that prior denoising has a rather strong positive effect on the quality of the achieved semantic unmixing results ($>$ 7db PSNR improvement), suggesting that the low SNR in the original data might indeed have caused the bad performance.

Since HHMI-D25$_{8bit}$ is stored in unsigned int8, meaning that pixel values are in $[0, 255]$, there are actually only a few distinct integer values presenting most of the data. Denoising this data, besides increasing the SNR, also increases the number of unique pixel values. We hypothesized that in addition to SNR, an overly discreet nature of pixel values might also be detrimental to semantic unmixing. To test this hypothesis, we imaged the HHMI-D25$_{16bit}$ data subset, where the unsigned int16 format was used to store the data, increasing the pixel intensity range to $[0, 65535]$. Doing so, we ensured that the SNR (ratio of average foreground value to average background value) is as similar as possible for both, the 8 and 16 bit versions of the HHMI-D25 data. Using the 16bit data did indeed improve the quality of predictions, also for the problematic third channel, as can be seen in Supplementary Figures S58 and S62. The quantitative metrics in Table C.3, however, do not capture the improvement we can perceive by comparing those figures. To fully validate our hypothesis regarding SNR and overly discrete pixel intensities, we imaged another data subset of HHMI-D25, namely HHMI-D25$_{16bit,0.25}$, which not only uses unsigned uint16 format but also bins four pixels into one (thereby increasing SNR on the cost of spatial resolution). On this data subset we defined Task XXXVI, and indeed observe a much improved semantic unmixing performance (see Table C.3 and Figure S59).

We also experimented with synthetic Gaussian and Poisson noise with HHMI-D25 dataset versions. The motivation was to start from a working setup, make the training and evaluation dataset noisy and inspect the performance degradation. For this purpose we picked HHMI-D25$_{16bit}$ and HHMI-D25$_{8bit,denoised}$. We added Gaussian noise ($\sigma$) and Poisson noise ($\lambda$). Given an image $x$, its noisy version can be expressed $Poi(x/\lambda) \cdot \lambda + \epsilon$, where $\epsilon \sim \mathbb{N}(0, \sigma)$ and $Poi()$ represents the Poisson distribution with parameter $\lambda$. As it was to be expected, the performance degrades with noise (see Tasks XXXII, XXXIV, and XXXV in Supplementary Figures S62, and S61, and a quantitative comparison in Supplementary Table C.3.

##### B.1.2 Out-of-distribution SNR

. We also use the set of models trained on different HT-LIF24 sub-datasets to understand how the performance degrades with out-of-distribution inputs. For this, we evaluate the performance of Micro$\mathbb{S}$plit trained on one exposure duration on the superimposed input images coming from a different exposure duration. We present the results in Figure S5. The different curves represent individual Micro$\mathbb{S}$plit models trained on one specific exposure duration sub-dataset as specified in the legend. On the x-axis, we have different evaluation sub-datasets, referred to by their exposure duration. From CARE-PSNR and $\mathbb{M}$icroMS-SSIM plots, one can observe that performance improves as one increases the exposure duration. Additionally, upon observing performance on $2ms$ and $500ms$ sub-datasets, one can see that in most cases, the larger the difference between the exposure duration of the training sub-dataset and evaluation sub-dataset, the higher the performance drop. For instance, if we look at CARE-PSNR plot, the two worst-performing Micro$\mathbb{S}$plit models on $2ms$ acquisition were trained on $20ms$ and $500ms$. And the two worst-performing Micro$\mathbb{S}$plit models on $500ms$ acquisition were trained on $2ms$ and $3ms$. On a different note, one can observe much less variation in SSIM and MS-SSIM plots. We discuss this aspect in Section E.

#### B.2 Spatial Correlation

The structures present in the cell are spatially correlated. For example, nuclei are typically in the central regions of cells, while the cell boundary is, by definition, on the boundary of a cell. The knowledge about the cell surface, therefore, can tell something about where the nucleus should or should not be present. We wanted to understand how important this spatial correlation is for our semantic unmixing task.

For this, we worked with the Pavia-P23 dataset where we modified the input patch process during training. The default method is to pick a random location in a frame, extract patches for both structures (target channels) from that location, and sum them to create superimposed input (*i.e. Training Mode I*). Using *Training Mode I*, the spatial correlation between the imaged structures is naturally maintained. To disrupt this, we conducted experiments with the following alterations.

In the first alteration, we kept *Training Mode I* for 50% of all training patches, but created the other 50% of training patches by picking two different random locations and adding them together to create an input patch (*i.e. Training Mode II*). Hence, we maintain sound spatial correlations between the structures to be unmixed in half the training data. In the second alteration, we create all training data patches according to *Training Mode II*, thus eliminating all spatial correlations between the structures that should be unmixed and forcing the trained network to rely fully on the structural appearance of the structures only.

We report the metrics CARE-PSNR and 𝕄icroMS-SSIM in Table ST8 which shows that the absence of spatial correlation indeed results in a drop in performance by $0.3 - 0.5$ dB PSNR.

#### B.3 Similarity of Structures to be Unmixed

Since our method relies heavily on the spatial appearance of the structures to be unmixed, we wondered how dissimilar two structures have to be to lead to good semantic unmixing results. Hence, it is best to image structures in single channels that are as dissimilar as possible.

In $\mu$Split [6], we proposed a synthetic semantic unmixing dataset based on combinations of sinusoidal curves that required longer-range structure integration. Although the structures were very simple, the network without using LC was unable to split the input and ended up only predicting the input for both target channels equally.

However, to better evaluate to what extent a network can separate structures that have a similar appearance, we designed the following experiments. Using the microtubule channel of the HT-LIF24 dataset, we created a 2-channel splitting task by mixing patches from the microtubule channel with other patches from the exact same channel. We reasoned that the network will not be able to split structures that are literally taken from the same set of data (see bottom row in Figure S2). To give the network a chance, we then started to alter one of the two superimposed copies by scaling the data (uppermost four rows in Figure S2). This leads to the superposition of data that is structurally still very similar, but one copy will have a slightly larger appearance.

In Figures S2 and S3 we show the results of these experiments, conducted with a range of different relative scaling between the two copies of the microtubule data. The results clearly show that the network is capable of semantic unmixing the structures in all cases, even if the scaling factor was as low as 1.125. Note also that it takes the network increasingly longer before the splitting performance starts lifting off and reaching its peak, as can be inferred from the inflection point and convergence behavior seen in the PSNR *vs.* training steps plot in Figure S3.

#### B.4 Unequal Channel Intensities

Another factor that plays a key role in the performance of Micro𝕊plit is the skewness in intensity between different target channels in the superimposed input image. This effect can be caused by diverging fluorophore densities for different structures, by fluorophore bleaching, or by using inadequate laser lines and/or laser power settings for some fluorophores. Any of this can lead to acquisitions where one or more of the structures are only weakly present in the superimposed input. We hypothesized that predictions for the brighter (dominant) structure should still be of good quality but that the quality for the dim structure might be worse.

To test this hypothesis, we worked with the 2-channel semantic unmixing tasks from the Pavia-P24 dataset, where we varied the laser power for the two channels we imaged from a balanced setting (50:50) to increasingly more skewed settings (*i.e.* 66:33 and 84:16). Note that the total laser power was kept the same. We acquired images at these three levels of skewness at three different total laser powers and resulting signal-to-noise ratios, giving us a total of 9 sets of image acquisitions. We show the results of all semantic unmixing experiments in Table ST2, where 'Skew' denotes the asymmetry in the power distribution ('Balanced', 50:50; 'Mid', 66:33; 'High', 84:16).

We found that, across three total laser power levels, the performance of the bright first channel generally increases as we go from Balanced to High skew. The second dim channel shows the inverse behavior. Note that since the samples we imaged at different imaging settings were not the same, each experiment is based on a different body of training and evaluation data. This causes additional fluctuations in the performance metrics we report and makes the presented results to be non-monotonic. If all 9 imaging sessions had captured the same samples, we would expect the results to show a monotonic trend.

***Unequal Channel Intensities - Two Extreme Examples***

For a first example, we worked with the nucleus and tubulin channels of the Chicago-Sch23 dataset. As always, we create the superimposed input by summing the two channels. Here, the nucleus is very weak (the channels being very skewed in their relative intensity), so much so that nuclei are de facto invisible to the naked eye. However, Micro𝕊plit was still able to unmix these channels at a reasonable quality. One can observe the superimposed input, the two targets, and the predictions of Micro𝕊plit in Figure S8.

For a second example, we inspect Task II which uses the HT-P23A dataset. Here, the two structures are microtubules and nuclei, with the latter being the weaker channel. Similarly to the first example, the weaker nucleus channel is effectively invisible in superimposed images, as can be seen in Figure S9. However, unlike in the previous example, the structure of nuclei in this dataset is less regular and more variable. Due to this and in light of the high uncertainty caused by the highly skewed channel intensities, Micro𝕊plit MMSE predictions for the nucleus channel become rather blurry, as can be seen in Figure S9. Still, in many practical applications, *i.e.* every time the detailed texture of the nuclei is not important, such predictions can still be sufficient for the desired data analysis to be carried out.

##### B.5 Sufficient Lateral Image Context

Our method combines the benefits of $\mu$Split and denoi𝕊plit. One of the benefits of $\mu$Split architecture we demonstrated in [6] is its ability to utilize additional surrounding spatial context for a given superimposed input patch through LC inputs and its ability to employ much deeper architectures. In [6], we showed that having sufficient spatial context helps with relatively large structures spanning hundreds of pixels.

#### C Additional Experiments

##### C.1 Micro𝕊plit vs. PICASSO

In this section, we compare the performance of Micro𝕊plit with PICASSO [20] (see also Figure S4). For semantic unmixing $k$ channels Micro𝕊plit needs as input a single superimposed image from the microscope whereas PICASSO needs $k$ images from the microscope which correspond to $k$ spectrally overlapping fluorophores. Due to this mismatch in the data requirement, a direct comparison is not feasible. So, to compare Micro𝕊plit with PICASSO, we generate synthetic inputs using our HT-LIF24 dataset. However, we argue that our way of generation does not degrade the performance of PICASSO but instead, it should be easier for PICASSO to predict on this data as compared to the real data.

Since fluorophores can be ordered according to the wavelength of the maximum intensity in their emission spectra, we first define such an order of our structure types. Next, for generating every channel of the input for PICASSO, we define three weights and take the weighted average of the three structures using these weights. The weights are set according to the order of the structures set above. For example, for the first channel, the weight given to the second structure will be higher than the weight given to the third structure. We generate two sets of weights, one being harder than the other. In the hard case, the dominant structure type is given 1.5 times more weight than the next dominant type and 3 times more weight than the least dominant type. In the easy version, the dominant structure type is given 2.5 times more weight than the next dominant type and 5 times more than the least dominant type. Once the input channels are generated, we add Gaussian noise of $\sigma = 500$ to each channel independently. We also train Micro𝕊plit on this data with the same Gaussian noise applied on top of the data. In Figure S4, we show the results on one random frame. In Table ST6 and Table ST7, we show the quantitative results.

##### C.2 *Training Mode I* vs *Training Mode II*- How Important are Spatial Correlations?

In this experiment, we inspect the performance drop between data acquisition types I and II. In cells, the location of different structures is often quite co-related with one another. For example, actin is mostly concentrated on the cell periphery whereas the nucleus is typically found at the center of the cell. In acquisition modes I and II, input is created by summing the crops from individual channels. In acquisition type I, since the channels are independently acquired, summing the crops of these different channels will generate an input patch where the naturally occurring co-location property cannot be preserved. In acquisition type II, since both channels are concurrently acquired, the inputs are created from summing the crops of different

| Task | Dataset | Synthetic Noise | PSNR | | | MicroMS-SSIM | | |
|---|---|---|---|---|---|---|---|---|
| XXIII | HHMI-D25$_{8bit}$ | - | 22.5 | 31.3 | 24.3 | 0.840 | 0.768 | 0.793 |
| XXXI | HHMI-D25$_{8bit,denoised}$ | - | 37.9 | 35.2 | 38.6 | 0.990 | 0.903 | 0.974 |
| XXXII | HHMI-D25$_{8bit,denoised}$ | $\sigma = 20, \lambda = 30$ | 31.8 | 27.3 | 32.4 | 0.939 | 0.664 | 0.859 |
| XXXIII | HHMI-D25$_{16bit}$ | - | 23.2 | 27.9 | 24.8 | 0.772 | 0.849 | 0.779 |
| XXXIV | HHMI-D25$_{16bit}$ | $\sigma = 2K, \lambda = 5K$ | 23.2 | 27.9 | 24.8 | 0.827 | 0.861 | 0.778 |
| XXXV | HHMI-D25$_{16bit}$ | $\sigma = 4K, \lambda = 10K$ | 23.0 | 27.4 | 24.6 | 0.746 | 0.838 | 0.700 |
| XXXVI | HHMI-D25$_{16bit,0.25}$ | - | 32.4 | 32.2 | 35.0 | 0.990 | 0.929 | 0.991 |

**Table ST1 Performance of MicroSplit on sub-datasets of HHMI-25.** This table presents quantitative results of MicroSplit across various tasks defined on parts of the HHMI-25 dataset. While the original Task XXIII on HHMI-D25$_{8bit}$ does not lead to satisfying predictions, mainly for channel 3 (see Supplementary Figure S57), tasks on similar data with higher SNR (Task XXXI, see Supplementary Figure S61, row 3), increased pixel diversity (Task XXXIII, see Supplementary Figure S62, row 2), or both (Task XXXVI, see Supplementary Figure S59) demonstrate notably improved semantic unmixing performance. Tasks XXXII, XXXIV, and XXXV are identical to Tasks XXXI and XXXIII, respectively, but with added synthetic noise, simply to demonstrate how the lower SNR drops the unmixing performance achievable with MicroSplit.

| Skew | Channel 1 Laser Power | | | Channel 2 Laser Power | | |
|---|---|---|---|---|---|---|
| | High | Mid | Low | High | Mid | Low |
| Balanced | 24.3 | 23.1 | 21.9 | 29.9 | 24.3 | 23.0 |
| Mid | 28.2 | 24.0 | 22.9 | 25.6 | 22.3 | 21.7 |
| High | 25.2 | 24.3 | 23.3 | 24.1 | 22.4 | 22.8 |

**Table ST2** Varying the laser power and skew with Pavia-P24 dataset. Skew column denotes the relative importance given to channel 1. High skew means larger laser power allocated to Channel 1 compared to Channel 2

channels with each crop taken from the same location in the micrographs and therefore these inputs preserve the naturally occurring co-location property. In this experiment we quantify the benefit of using this co-location information.

We work with the HT-T24 dataset which falls under *Training Mode III*. We train three models. In the first model, we create the input using *Training Mode I*, that is, we create the input by picking target patches from the same location and therefore maintain the spatial co-relation. In the second model, we use *Training Mode II*, meaning that we pick crops from random locations from the different channels and use them to create the input. This model naturally does not have access to naturally occurring spatial co-relation information in its training data. The third model is trained using *Training Mode III*. We evaluate all three models on the held-out test set where the inputs have spatial co-relation preserved and are not synthetic, that is, they are imaged from the microscope. In Table ST3, we find that the first model outperforms the second by **1.3** CARE-PSNR and **0.009** MicroMS-SSIM. Naturally, the model trained with *Training Mode III* is best and outperforms *Training Mode I* by **0.6** CARE-PSNR and **0.004** MicroMS-SSIM.

#### C.3 *Training Mode I vs. Training Mode III*- Summed *vs.* Acquired Inputs

Here, we quantify how much the performance degrades if, during training, the input is created by simply summing the two channels as compared to input coming directly from the microscope. We find that while there is indeed a performance drop as can be seen in Table ST3, the drop is not detrimental. This experiment shows the utility of our approach in the case when synthetic input is used for its training but for evaluation, inputs coming directly from the microscope are used.

#### C.4 MicroSplit enables a more effective use of the available photon budget

**By filtering fewer photons:** Traditional multiplexed imaging relies on emission filters that selectively pass photons from one fluorophore at a time in order to minimize spectral overlap. As a consequence, a substantial fraction of emitted photons is discarded, and relaxing the filters leads to bleedthrough artifacts. MicroSplit changes this trade-off because multiple structures can be imaged in the same acquisition. This allows microscopists to use substantially broader emission filters, collecting photons from several fluorophores simultaneously, without introducing the ambiguities that would otherwise arise in a multi-channel setting.

To roughly quantify this advantage, we analyzed three fluorophores from the HT-LIF24 dataset (DAPI, FITC, TRITC), using their emission spectra (downloaded from fpbase.org) normalized as probability mass

| Model | PSNR | SSIM |
|---|---|---|
| Training Mode I: input = $C_1 + C_2$ | 35.9 | 0.956 |
| Training Mode II: input = $C_1 + C_2$ (shuffled) | 34.6 | 0.947 |
| Training Mode III: input comes from microscope | 36.5 | 0.960 |

**Table ST3** Performance comparison for different Acquisition Modes. We use the Sox2 vs Golgi splitting task of the HT-T24 dataset for this purpose. For Acquisition Mode I and II, real input image is not used during training. Instead, input is created by synthetically summing the two target channels. In all cases, evaluation is done on the held-out test set using the real input channel, that is, on the input which is not synthetic and comes directly from the microscope.

| | C1 | | | | C2 | | | |
|---|---|---|---|---|---|---|---|---|
| | (2D) Z=1 | Z=5 | Z=9 | Z=15 | (2D) Z=1 | Z=5 | Z=9 | Z=15 |
| CARE-PSNR | 35.8 | 38.7 | 39.5 | 39.7 | 30.9 | 33.7 | 34.5 | 34.7 |
| MicroSSIM | 0.865 | 0.878 | 0.885 | 0.886 | 0.729 | 0.757 | 0.772 | 0.767 |
| MicroS3IM | 0.950 | 0.970 | 0.973 | 0.974 | 0.929 | 0.950 | 0.956 | 0.956 |

**Table ST4** Performance improvement with 3D models on HT-H24 dataset. As we increase the number of z-slices fed to the model, we see the performance improve in all our metrices.

| | Set I | | | Set II | | |
|---|---|---|---|---|---|---|
| | $C_1$ | $C_2$ | $C_3$ | $C_1$ | $C_2$ | $C_3$ |
| Input $C_1$ | 0.50 | 0.33 | 0.17 | 0.625 | 0.25 | 0.125 |
| Input $C_2$ | 0.33 | 0.50 | 0.33 | 0.25 | 0.625 | 0.25 |
| Input $C_3$ | 0.17 | 0.33 | 0.50 | 0.125 | 0.25 | 0.625 |

**Table ST5** We use the weights to mix the three channels $C_1$, $C_2$ and $C_3$.

functions. We compared the fraction of photons transmitted by conventional multi-color filter configurations to the photons captured when using a broad, highly permissive filter suitable for Micro𝕊plit. Across three representative scenarios, with emission filter thresholds chosen to (*i*) maximize photon collection, (*ii*) reduce bleedthrough by 25%, and (*iii*) reduce bleedthrough by 50%, with conventional multiplexed imaging being respectively 22%, 34%, and 55% less photons efficient than the results obtained using Micro𝕊plit (see top half of Supplementary Figure S6).

In practical terms, this means that even in bleedthrough-optimized multiplexed imaging, each channel discards a large fraction of emitted photons, whereas Micro𝕊plit can reclaim much of this loss by aggregating photons from multiple fluorophores in a single measurement.

**By enabling gentler imaging due to denoising:** Our method, next to performing semantic unmixing, also performs unsupervised denoising. Since denoising improves the signal-to-noise (SNR) of images it is applied to, microscopists can acquire the raw data more gentle, accepting a lower initial SNR [3]. To illustrate this on a concrete example, we assessed the similarity of a biological structure imaged at various exposure times between 2ms and 20ms with very high SNR data of the same regions of interest acquired at 500ms exposure time. More concretely, we have conducted these experiments on the 3 channel data (Nucleus, Microtubules, Kinetocore) of the HT-LIF24 dataset. As shown in Supplementary Figure S6 (bottom), even the denoised 2ms exposure micrographs lead to a considerably higher quality w.r.t. the 500ms images than even the 5ms raw data, for Channel 1 even compared to the 20ms raw acquisitions, suggesting at least a 3 to 10-fold reduction of the required photon budget and therefore enabling users of Micro𝕊plit to image considerably more gentle to reach the same quality required for downstream-processing.

### D Details on Uncertainty Quantification and Calibration

In this section, we detail our uncertainty estimation and calibration module. Our approach builds on the methodology of denoi𝕊plit [7], originally inspired by [15], with a modification to the calibration process as described below. Our variational approach is inherently capable of sampling, meaning it can generate slightly different outputs for the same input each time a prediction is made. We leverage this property to produce multiple predictions for a single input, resulting in several predicted values per pixel. Using these, we calculate the standard deviation for each pixel. To ensure these pixel-wise standard deviations correspond closely to the actual prediction errors, we apply a straightforward linear scaling. Opting for

| Model | Channel 1 | | | Channel 2 | | | Channel 3 | | |
|---|---|---|---|---|---|---|---|---|---|
| | A | B | C | A | B | C | A | B | C |
| PICASSO Set I | 24.1 | 0.633 | 0.751 | 24.1 | 0.898 | 0.863 | 31.6 | 0.858 | 0.914 |
| PICASSO Set II | 24.2 | 0.638 | 0.757 | 24.1 | 0.904 | 0.864 | 32.2 | 0.841 | 0.911 |
| Micro𝕊plit | **33.9** | **0.730** | **0.902** | **34.9** | **0.909** | **0.884** | **38.21** | **0.910** | **0.958** |

**Table ST6** Quantiative comparison with PICASSO. A: RI-PSNR, B: MicroSSIM, C: MicroS3IM

| Model | Channel 1 | | | Channel 2 | | | Channel 3 | | |
|---|---|---|---|---|---|---|---|---|---|
| | A | B | C | A | B | C | A | B | C |
| PICASSO Set I | .779 | .021 | .020 | .587 | .007 | .012 | .454 | .026 | .005 |
| PICASSO Set II | .779 | .032 | .019 | .587 | .007 | .012 | .478 | .038 | .006 |
| Micro𝕊plit | .839 | .025 | .008 | .475 | .005 | .010 | .613 | .012 | .003 |

**Table ST7** Standard error table for Table ST6. A: CARE-PSNR, B: MicroSSIM, C: 𝕄icroMS-SSIM

| Corr. | Balanced | | | | Mid-Skew | | | | High-Skew | | | |
|---|---|---|---|---|---|---|---|---|---|---|---|---|
| | PSNR | | 𝕄icroMS-SSIM | | PSNR | | 𝕄icroMS-SSIM | | PSNR | | 𝕄icroMS-SSIM | |
| 0% | 23.9 | 29.5 | 0.594 | 0.731 | 27.7 | 25.2 | 0.755 | 0.656 | 24.9 | 23.8 | 0.695 | 0.552 |
| 50% | 24.1 | 29.6 | 0.614 | 0.761 | 28.1 | 25.5 | 0.761 | 0.657 | 25.1 | 23.9 | 0.781 | 0.699 |
| 100% | 24.3 | 29.9 | 0.682 | 0.839 | 28.2 | 25.6 | 0.780 | 0.696 | 25.2 | 24.1 | 0.722 | 0.623 |

**Table ST8** Here, we inspect the importance of spatial correlation in the structures. In row 1, input is created by summing patches taken from different locations. In row 2, for 50% of the time, input is created by summing patches taken from different locations. In the remaining 50% of times, input is created by summing patches taken from the same location. In row 3, all training input is created by always summing together co-located patches. Evaluation is done on co-located patches.

a linear transformation, rather than a more complex method, helps minimize the risk of overfitting. This simple linear transformation based approach allows us to easily estimate pixel-wise errors for a test input: we simply run multiple predictions, compute the standard deviation for each pixel, and then apply the learned linear transformation. Importantly, this process only requires the test input, the trained model, and the parameters of the linear transformation.

Next, we provide a technically accurate description of the methodology. For each input image $x$ (of dimensions $H \times W$), we generate $k = 50$ predictions. At every pixel location $p$ in channel $i$, this gives us $k$ predicted values, which we use to compute the pixel-wise standard deviation $\sigma_i[p]$. Next, for calibration, for each channel $i$, we sort the computed pixel-wise standard deviations and bin them over $l = 30$ equally sized bins. Specifically, we implement the above-mentioned logic by storing pixel location $p$ of the relevant pixels in these bins $\{B_i^1, B_i^2 \dots B_i^l\}$. We then compute the root mean variance (RMV) and root mean squared error (RMSE) for each channel $i$ and bin $j$ as

$$\mathrm{RMV}_i(j) = \sqrt{\frac{1}{|B_i^j|} \sum_{p \in B_i^j} \sigma_i[p]^2} \quad \text{, and} \tag{5}$$

$$\mathrm{RMSE}_i(j) = \sqrt{\frac{1}{|B_i^j|} \sum_{p \in B_i^j} (c_i[p] - \hat{c}_i[p])^2}, \tag{6}$$

where $c_i$ and $\hat{c}_i$ represent the ground truth image and the estimated image for the $i^{\mathrm{th}}$ channel respectively. Subsequently, we fit a linear relationship between RMSE and RMV for each channel $i$, learning the coefficients $\alpha_i$ and $\beta_i$ that minimize the objective

$$\arg\min_{\alpha_i, \beta_i} ||\mathrm{RMSE}_i - (\alpha_i \cdot \mathrm{RMV}_i + \beta_i)||_2^2. \tag{7}$$

Note, that denoi𝕊plit employed pixel-level regression instead of bin-based regression. However, that method has a key limitation: since images often contain many background pixels, the regression tends to be biased toward these regions, resulting in poor calibration for foreground pixels, which is problematic.

During evaluation, we sample multiple predictions for a given input. For a channel $i$, we compute the pixel-wise standard deviation $\sigma_i$ over the multiple samples. The estimate of pixel-wise RMSE is given by

| Dataset | 2D/3D | Pixel size [nm] | Exposure [ms] | Pixel/Voxel (million) |
|---|---|---|---|---|
| Pavia-P24 | 2D | 68 | 200 | 27.9 |
| HT-T24 | 2D | 110 | 400 | 2831.3 |
| HT-LIF24 | 2D | 285 | 2, 3, 5, 20, 500 | 4126.8 |
| Chicago-Sch23 | 2D | 270 | 100 | 7516.2 |
| HT-H24 | 3D | 285 | 300 | 1985.8 |
| HHMI-D25$_{8bit}$ | 3D | 45 | - | 3850.4 |
| CBG-Z18 | 3D | 300 | - | 1362.1 |
| CBG-N18 | 3D | 196 | - | 510.7 |
| HT-P23A | 3D | 37 | - | 637.5 |
| HT-P23B | 3D | 42 | - | 416.5 |

**Table ST9 Dataset Overview:** This table summarizes key quantitative characteristics of the datasets used in the presented work. The column 'Pixel/Voxel' shows the total number of pixels/voxels in the entire dataset. Note that individual training tasks only use a (at times) small subset of the entire dataset they are derived of.

$\text{RMSE}_i = \sigma_i * \alpha_i + \beta_i$. It is worth noting that the estimation of $\alpha_i$ and $\beta_i$ is done using the validation data and evaluation for the calibration is done on the test data.

With Micro𝕊plit, we have presented a simple, yet effective way to estimate pixel-wise uncertainty estimates and evaluate the calibration of the predicted uncertainty. One of the advantages of the presented calibration procedure is that it does not alter the original predictions but instead learns a mapping that best predicts the measured error.

### E Evaluation Metrics

SSIM (Structural Similarity Index Measure) [25, 26] and PSNR (Peak Signal-to-Noise Ratio) are one of the most popular metrics used in regression tasks. While MSE-based metrics like PSNR estimate pixel-wise distance between the target and the prediction, SSIM captures structural similarity. For large images that may have sub-regions having different structural characteristics, people found it useful to work with MS-SSIM (Multi-Scale SSIM) [27] instead of SSIM. Since microscopy images are typically large in pixel dimensions, MS-SSIM therefore seemed a better alternative over SSIM. However, our earlier work showed that when working with fluorescence microscopy data, especially with differing exposure durations, PSNR, SSIM, and MS-SSIM are all ill-suited [8]. Among several issues presented in the above-mentioned work, here we illustrate the issue most relevant to Micro𝕊plit. The issue is that the input to Micro𝕊plit is typically a noisy image which means that it was imaged using a small exposure duration. This means that the image covers a smaller portion of the full dynamic range. In other words, the pixel intensities found in the input image are relatively low in magnitude. Since the network was trained with noisy target images, which had a similar dynamic range, the prediction by the network also covers a similar amount of dynamic range. However, for the HT-LIF24 dataset, we have the corresponding high SNR target, which covers a much larger portion of the dynamic range and so has larger pixel values. So, the problem is: how to compare a prediction having a low-dynamic range with a high-dynamic range image? The core idea in the solution used in CARE-PSNR [3] and 𝕄icroMS-SSIM [8] is to estimate an optimal linear transformation for the prediction. This linear transformation makes the pixel intensity values of the scaled prediction as close to the corresponding target channel as possible. The relevant metric (PSNR, SSIM, or MS-SSIM) is then computed between the target and the scaled prediction. We have used CARE-PSNR and 𝕄icroMS-SSIM in this work.

### F Dataset Details

For this work, we acquired several datasets, with some designed for specific purposes. We are making all these datasets public. Next, we briefly describe the datasets. In Table F, we provide a brief overview of the different datasets used in this work. For more details, please refer to the metadata stored in the dataset files.

#### F.1 The HT-P23A and HT-P23B datasets

This dataset was acquired by the Pigino group at Human Technopole, Milan, Italy. Data generation procedure was based on [23].

***Cryo-ExM of MDCK-II cells***

MDCK-II cells were seeded on 12 mm coverslips contained within 6-well plates at a density of 300,000 cells / well, grown at 37 degree Celsius and 5% $CO_2$. When cells reached approximately 50% confluence, a subset of coverslips was incubated with 150 nM MitoTracker Deep Red FM (ThermoFisher Scientific, Catalog Number M22426) for 30 minutes and checked for dye incorporation using an EVOS M5000 Imaging System (ThermoFisher Scientific). Cells were then immediately processed for cryo-ExM1. In brief, cells on coverslips were rapidly plunged frozen in -180 degree Celsius liquid ethane using a manual plunger. Coverslips were transferred to an Eppendorf tube containing frozen, desiccated acetone supplemented with .1% paraformaldehyde (PFA) and .02% glutaraldehyde (GA) for overnight freeze substitution in dry ice. The next morning, the coverslips underwent successive ethanol baths in progressively decreasing percentages, as follows: ethanol 100% (5 min), ethanol 100% (5 min), ethanol 95% (5 min), ethanol 95% (5 min), ethanol 70% (3 min), ethanol 50% (3 min) and PBS. Substituted cells then underwent anchoring in 2% acrylamide and 1.4% formaldehyde in PBS for 3 hrs. Coverslips were then exposed to activated monomer solution containing sodium acrylate (19%), acrylamide (10%), N,N'-methylenbisacrylamide (0.1%) and ddH2O, activated by APS and TEMED (0.05%). Expansion gels polymerized for 1 hr at 37 degree Celsius, and were then denatured in denaturation solution containing SDS (200 mM), NaCl (200 mM) and Tris (50 mM) in ultrapure water at 95 degree Celsius for 1.5 hrs. Expanded gels were then washed in PBS 3X, 10 minutes for each wash. Final concentrations of solutions are reported.

***Expansion immunolabeling***

The following day, gels were stained with anti-alpha tubulin (ABCD Antibodies, ABCD_AA345) and anti-beta tubulin antibodies (ABCD antibodies, AA_344), diluted `1:300` in PBS 2% BSA. Half of the MitoTracker labeled gels were given Mouse IgG2a raised antibody, and the rest of the gels were stained with Guinea Pig IgG raised antibody. Tubulin-labeled gels were incubated at 37 degrees Celsius for 3 hrs while gently shaking. After primary labeling, gels were washed 3X in PBS 0.1% Tween, 10 minutes for each wash. Mouse IgG2a tubulin-labeled gels were then stained with Goat Anti-Mouse IgG Secondary Antibody AlexaFluor 488 (ThermoFisher Scientific, Catalog Number A-11001), while Guinea Pig IgG tubulin-labeled antibodies were stained with Goat Anti-Guinea Pig IgG Secondary Antibody AlexaFluor 647 (ThermoFisher Scientific, Catalog Number A-21450). Secondary antibody labeled gels were then incubated at 37 degrees Celsius for 3 hrs while gently shaking. The gels were washed 3X in PBS 0.1% Tween, 10 minutes for each wash. Gels were further washed 2X, 30 minutes in ddH2O, and then placed in ddH2O overnight for expansion.

In HT-P23A, the fluorescence intensity of labeled microtubules strongly overpowered the mitochondrial signal in the condition in which both microtubules and mitochondria were co-fluorescent in the 647 channel. To ensure a more equal signal, we imaged HT-P23B, where we reduced the concentration of tubulin Guinea Pig IgG labeling to 1:450, and the concentration of Anti-Guinea Pig IgG Secondary Antibody to 1:450 to ensure a more equal signal.

***Imaging expansion gels***

Expanded gels were cut into approximately 1.5 $cm^2$ pieces and placed on a 24 mm coverslip coated in poly-D-lysine (Sigma, Catalog Number P7886) and placed within a 35 mm imaging chamber. Imaging was performed on a Zeiss LSM980-NLO, in confocal mode. Z-stacks with a step size of 0.15 $\mu m$ were collected with a frame size of $2024 \times 2024$ pixels using a 63X oil immersion objective (1.4NA).

#### F.2 The Pavia-P23 dataset

This dataset was acquired at the Synthetic Physiology Lab at the University of Pavia, Pavia, Italy.

***HaCaT cell line culture and media***

The HaCaT cell line was kindly gifted by Professor Hugo de Jonge (Department of Molecular Medicine, University of Pavia). Cells were maintained in DMEM/F12 no phenol red (catalog `#21041-025`, Gibco) with 10% Fetal Bovine Serum (FBS) (catalog `#10270-106`, Gibco) and 1% Penicillin/Streptomycin (P/S) (catalog `#A001`, HiMedia). Cells were never allowed to go beyond 75–80 % confluency during routine splitting.

| | Laser Power % (50/50) | | Laser Power % (33/66) | | Laser Power % (16/84) | |
|---|---|---|---|---|---|---|
| | 446nm | 477nm | 446nm | 477nm | 446nm | 477nm |
| SNR: high | 40 | 40 | 27 | 53 | 13 | 67 |
| SNR: mid | 20 | 20 | 13 | 27 | 7 | 33 |
| SNR: low | 10 | 10 | 7 | 13 | 3 | 17 |

**Table ST10** Laser power distribution in the several sub-datasets of Pavia-P24 dataset

###### *Generation of the HaCaT FUCCIplex-prototype clonal cell line*

The HaCaT cell line expressing the structural actin and tubulin fluorophores and the FUCCIplex sensor was generated as described previously in [28]. Briefly, cells were plated at the density of 6 × 104 cells per well in a 12-well plate and infected with the FUCCIplex lentivirus (10$\mu l$ of concentrated FUCCIplex lentivirus particles (> 108 TU/ml, VectorBuilder). Positive cells were then selected with 20$\mu l/ml$ of Hygromycin B (50$mg/ml$ in PBS) (catalog `#10687010`, Invitrogen) and expanded. Positive cells expressing the FUCCIplex sensor were then infected with a genetically encoded Lifeact-ACTB-RFP probe (rLV Ubi-LifeAct-TagRFP, 1 × 107$TU/ml$, 10$\mu l$) that specifically tags the (F-)actin filaments with a red fluorescent reporter. Lastly, the alpha-tubulin locus (TUBA1B, NM006082.3) was genome-edited via CRISPR/Cas9 using the ThermoFisher Scientific True Tag system (catalog `#A42992`) to tag the tubulin N-terminus of the protein with an EGFP fluorophore. Positive cells were clonally selected and expanded. The TrueTag homology arm primer forward sequence is: `TCCTGTCGCCTTCGCCTCCTAATCCCTAGCCACTATGGGAGGTAAGCCCTTGCATTCG`; The reverse is: `CCTGAAAGCAGCCGGGAGCCGCACGGCTTACTCACACCGCTTCCACTACCTGAACC`). The TrueGuide Synthetic gRNA (sgRNA) sequence is `GCACGGCTTACTCACCATAG`. For imaging experiments, HaCaT cells were plated in culture medium into porcine skin-coated (0.2%, w/v) ibidi $\mu$-slide 8 well-chambered coverslips (catalog `#80807`, ibidi) at a density of 50.000 cells/well.

###### *Micro𝕊plit: Imaging Methods for Pavia-P24 dataset*

Imaging was performed using a Nikon Ti2 Eclipse inverted microscope integrated with a Crest V3 X-Light spinning disk confocal unit and a Teledyne Photometrics Kinetix Scientific sCMOS camera. The dataset was acquired using a CFI SR HP Plan Apo Lambda S 100XC objective lens (silicon oil immersion, N.A. 1.35, WD 0.31–0.28$mm$, MRD73950).

A Lumencore Celesta Light Engine (TSX5030FV, Lumencore) provided illumination for fluorescence imaging, delivering light at multiple wavelengths (405$nm$, 446$nm$, 477$nm$, 520$nm$, 546$nm$, 638$nm$, 740$nm$).

Specifically, the dataset was acquired using 446$nm$ and 477$nm$ laser lines. In the 477$nm$ line optical configuration, a multiband dichroic and excitation filter set (MXR00543 CELESTA-DA/FI/TR/Cy5/Cy7-A, CELESTA) was used, along with a FITC emission filter (MXR00541 FITC, CELESTA). Instead, the 446$nm$ optical configuration was equipped with dual-band filters for dichroic, excitation, and emission (MXR00544 - CELESTA CFP/YFP, CELESTA).

In addition, the microscope has an advanced environment setup to support live imaging experiments. The Oko-Cage incubation system (Okolab) equipped with environment controllers can maintain optimal conditions for live-cell imaging, including a temperature setpoint of 37 degrees Celsius, passive humidity control, and $CO_2$ levels of 5%.

Nikon NIS Elements software (version 5.42.06, Nikon) was used to manage the imaging system. Spinning disk confocal mode was selected for this dataset of two-channel z-stacks. In detail, channels 477$nm$ and 446$nm$ were imaged sequentially with identical exposure settings (200$ms$). Splitting performance was tested by changing two factors: the signal-to-noise(SNR) level (high, mid, and low) and the relative laser power percentage of each channel, as shown in the table ST10.

Multiple Z-stacks of 6 planes with a step size of 1 µm were acquired to populate the dataset for each condition.

#### F.3 The HT-H24 dataset

The HT-H24 dataset was imaged by the Harschnitz group at Human Technopole and contains immunofluorescent staining of SOX2 (555) and MAP2 (488) in DIV25 dorsal forebrain organoids generated from WTC-11 (UCSFi001-A) induced pluripotent stem cells (iPSCs) (UCSFi001-A). Human iPSC-derived forebrain organoids were generated following a triple-inhibition patterning protocol, without LIF supplementation, as published in a previous work [29]. At days in vitro (DIV) 25, forebrain organoids were fixed in 4% paraformaldehyde at 4 degrees Celsius overnight, transferred in PBS with 30% sucrose until fully immersed, embedded in OCT compound (Scigen), frozen rapidly, and stored at -20 degrees Celsius. 15 micrometer-thick sections were sliced on a cryostat (Leica). For immunofluorescence staining, organoid slices were rinsed in PBS, incubated with Retrieval Solution (Dako) for 45 minutes at 70 degrees Celsius, permeabilized with 0.5% Triton X-100 in PBS1x for 20 minutes, and blocked with 0.1% Triton X-100, 10% Donkey Serum in PBS1x for 1 hour at room temperature. Sections were incubated with either Rat $\alpha$ SOX2 1:200 (Invitrogen, 14-9811-82), and Chicken $\alpha$ MAP2 1:5000 (Invitrogen, PA1-10005), or Rat $\alpha$ CTIP2 1:500 (Abcam, ab18465) overnight, and counterstained with 1:1000 Donkey $\alpha$ Chicken Alexa Fluor 488 (Jackson ImmunoResearch, 703-545-155), or 1:1000 Donkey $\alpha$ Rat Alexa Fluor$^{TM}$ Plus 555 (Invitrogen, A48270). Nuclei were stained with DAPI (Thermo Fisher, 62248). Immunostained sections were scanned using Ti2 CREST spinning disk (Nikon), with objective magnification of 40x.

##### F.4 The HT-T24 dataset

This dataset is a 3 channels confocal microscopy imaging of fixed E37 ferret brain sections acquired by the Taverna group at Human Technopole. It is a 3 channel 2D dataset. The first two channels contain the SOX2 (transcription factor used to label stem cell nuclei) and Golgi marker Grasp65, and the last channel contains the superimposed image containing the above-mentioned two structures. For more details, please look into the metadata of the dataset files. **Organism** is Ferret (Mustela furo) and the **Sample** is Cryosection of E37 ferret brain stained with SOX2 and GRASP65 antibodies.

***Experimental Animals***

All experimental procedures were conducted in agreement with the German Animal Welfare Legislation after approval by the Landesdirektion Sachsen (license for ferret TVV2/2015). Animals used for this study were kept in standardized hygienic conditions at the Biomedical Services Facility (BMS) of the MPI-CBG with free access to food and water. All experiments were performed in the dorsolateral telencephalon of ferret embryos, at a medial position along the rostro-caudal axis at a stage corresponding to mid-neurogenesis.

***Protocol for immunofluorescence staining***

After incubation at 70oC for 30 min in Dako Target Retrieval Solution, Citrate pH 6, cryosections were permeabilized with 0.3% Triton X-100 in PBS for 30 minutes at room temperature. Blocking was performed in a blocking solution (0.2% gelatin, 300 mM NaCl, 0.3% Triton X-100 in PBS) for 30 min. Primary antibodies were incubated in the blocking solution for O/N at 4 degree Celsius. The following antibodies were used: SOX2 (goat polyclonal, AF2018, 1:200, R&D Systems) and GRASP65 (rabbit polyclonal, PA3-910, 1:200, Invitrogen). Subsequently, the sections were washed three times in PBS and incubated for 1 hour at room temperature with the following secondary antibodies: donkey anti-Goat IgG(H+L) Highly Cross-Adsorbed Secondary Antibody, Alexa Fluor Plus 555 (A32816, 1:500, Invitrogen) and donkey anti-Rabbit IgG(H+L) Highly Cross-Adsorbed Secondary Antibody, Alexa Fluor Plus 647 (A32795, 1:500, Invitrogen). After three washes in PBS, stained sections were mounted with Mowiol.

***Acquisitions of HT-T24 dataset***

All images were acquired using a spinning disk confocal system, consisting of a CrestOptics V3 light scan-head (configured with 50 um-pinholes) mounted on a Nikon Ti2-E inverted microscope equipped with a motorized stage and 4 Photometrics Prime 95B 25 mm cameras (pixel size 11 um). The samples were acquired in confocal mode with a Plan Apochromat Lambda S 100x/1.35 silicon immersion objective using Celesta Lumencor solid-state lasers as the light source. Fluorescence was collected using the following elements: Channel 1 (C1) - excitation wavelength 638 nm laser lines at 20%, excitation filter and dichroic mirror MXR00543-CELESTA-DAPI/FITC/TRITC/Cy5/Cy7-Full Multiband Penta, Cy5 emission filter; Channel 2 (C2) – excitation wavelength 547 nm laser lines at 20%, excitation filter and dichroic mirror MXR00543-CELESTA-DAPI/FITC/TRITC/Cy5/Cy7-Full Multiband Penta, TRITC emission filter; superimposed channel (Input) - excitation wavelength 547 nm and 638 nm laser lines both at 20%, excitation filter and dichroic mirror MXR00543-CELESTA-DAPI/FITC/TRITC/Cy5/Cy7-Full Multiband Penta, no emission filter. All images were acquired with the same camera parameters: binning 1, 16 bit and 400 ms of exposure time. For every field of view, a Z-stack with a $1\mu m$-step size was acquired. Once the parameters of acquisition had been defined, they were kept constant. The software used for all acquisitions was NIS Elements AR 5.42.02 (Nikon).

#### F.5 The HT-LIF24 dataset

This dataset was acquired at the Light Imaging Facility at Human Technopole, Milan, Italy.

***Sample preparation of HT-LIF24 dataset***

HeLa cells were maintained in DMEM (EuroClone) containing 10% fetal bovine serum (Thermo Fisher Scientific), supplemented with 2mM L-glutamine (EuroClone) and penicillin/streptomycin both $100\mu g/ml$ (EuroClone), at 37 degree Celsius in a humidified atmosphere and 5% $CO_2$. Cells were plated onto glass `#1.5` coverslips for immunofluorescence microscopy and then fixed with 4% PFA for 10 min. They were permeabilized with 0.1% Triton X-100 and 0.2% BSA (Bovine Serum Albumin) in PBS for 10 min. Blocking was performed in a blocking solution (2% BSA in PBS) for 30 min. Primary antibodies were incubated in the blocking solution for 1 hr. The following antibodies were used- anti-$\alpha$-tubulin mouse IgG monoclonal (T5168, Sigma-Aldrich; `1:100`); anti-laminin B1 rabbit IgG polyclonal (ab16048, Abcam; `1:200`); anti-centromere protein human IgG polyclonal (15-234, Antibodies Incorporated; `1:400`). Subsequently, after washing in PBS, cells were incubated for 40 min with the following secondary antibodies in blocking solution- Alexa Fluor 488 donkey anti-mouse IgG (A-21202, Thermo Fisher Scientific; `1:400`); Cy3 donkey anti-rabbit IgG (711-165-152, Jackson Immunoresearch; `1:400`); Cy5 goat anti-human IgG (109-175-088, Jackson Immunoresearch; `1:50`). Finally, cells were also counterstained with DAPI (4',6-Diamidino-2-phenylindole dihydrochloride, D9542, Sigma-Aldrich; `1:40000`) for 15 min before the mounting step. All the steps were performed at room temperature. The samples were then mounted in Mowiol-DABCO and acquired with a spinning disk system as described below. At least 200 non-overlapping and randomly distributed fields of view were acquired and analyzed.

***Acquisitions of HT-LIF24 dataset***

All images were acquired using a spinning disk confocal system, consisting of a CrestOptics V3 Light scan-head (configured with $50\mu$m-pinholes) mounted on a Nikon Ti2-E inverted microscope equipped with a motorized stage and a Photometrics Prime 95B 25 mm camera (pixel size 11 $\mu$m). The samples were acquired with a Plan Apochromat Lambda S 40x/1.25 silicon immersion objective using Celesta Lumencor solid-state lasers as the light source. Fluorescence was collected using 19 different lightpath configurations. In each configuration, a penta-band excitation filter (`MXR00543-CELESTA-DAPI/FITC/TRITC/Cy5/Cy7-Full` Multi-band Penta), a penta-band dichroic filter (`MXR00543-CELESTA-DAPI/FITC/TRITC/Cy5/Cy7-Full` Multiband Penta) and a penta-band emission filter (Semrock `FF01-441/511/593/684/817-25`) were used. Ground truth images were acquired using a specific additional band-pass emission filter inserted before the penta-band emission filter (in particular the DAPI, FITC, TRITC, and Cy5 emission filters were added for GT-A, GT-B, GT-C, and GT-D configurations, respectively), while for all the other channels only the penta-band emission filter was used. The excitation wavelengths of the 19 channels were set as follows - GT-A - 405 nm (40%); GT-B - 477 nm (35%); GT-C - 547 nm (5%); GT-D - 638 nm (50%); A - 405 nm (40%); B - 477 nm (35%); C - 547 nm (5%); D - 638 nm (50%); AB - 405 nm (40%), 477 nm (35%); AC - 405 nm (40%), 547 nm (5%); AD - 405 nm (40%), 638 nm (50%); BC - 477 nm (35%), 547 nm (5%); BD - 477 nm (35%), 638 nm (50%); CD - 547 nm (5%), 638 nm (50%); ABC - 405 nm (40%), 477 nm (35%), 547 nm (5%); ABD - 405 nm (40%), 477 nm (35%), 638 nm (50%); ACD - 405 nm (40%), 547 nm (5%), 638 nm (50%); BCD - 477 nm (35%), 547 nm (5%), 638 nm (50%); ABCD - 405 nm (40%), 477 nm (35%), 547 nm (5%), 638 nm (50%). All images were acquired at binning 1. For each field of view, a series of images was captured using the following exposure times - 2 ms, 3 ms, 5 ms, 20 ms, 500 ms. The software used for all acquisitions was NIS Element AR 5.42.02 (Nikon).

Simply put, it has 4 different structures- (i) A - the whole nuclei (DAPI staining), (ii) B - microtubules: one component of the cytoskeleton, which is a filamentous system in the cytoplasm (alpha-tubulin staining), (iii) C - nuclear envelope- the membrane that separates the nucleus from the cytoplasm (Lamin B1 staining), (iv) D - the kinetochore/centromere- specific area along the chromosomes-DNA which is used to connect the chromosomes themselves to the microtubules during mitosis (CREST staining).

#### F.6 The Chicago-Sch23 dataset

This dataset is four-color structured illumination super-resolution microscopy imaging of live human BJ fibroblast cells acquired at the Scherer Lab at the University of Chicago, Chicago, USA [24]. It has 4 channels of different structures: (1) actins (CellMask Orange), (2) mitochondria (MitoTracker Green), (3) microtubules (Tubulin Tracker Deep Red), and (4) nuclei (Hoechst).

***Sample preparation***

Human BJ fibroblast cells were cultured in high-glucose DMEM (Life Technologies, 10569) supplemented with 10% fetal bovine serum (Life Technologies, 26140) and penicillin-streptomycin. Cells were maintained in a humidified incubator at 37 degrees Celsius with 5% carbon dioxide. Prior to imaging, live cells were washed with PBS (Life Technologies, 15140) and trypsinized using 2.5 mL of 0.05% trypsin (Life Technologies, 25300) at 70–80% confluence. The cells were then transferred to 35 mm glass-bottom dishes (MatTek, P35G-1.5-14-C) for microscopy.

The staining solution was prepared by diluting Tubulin Tracker Deep Red (Thermo Fisher Scientific, $T34077$) to $1x$, CellMask Orange Actin Tracking Stain (Thermo Fisher Scientific, A57247) to $2x$, MitoTracker Green FM (Thermo Fisher Scientific, $M7514$) to 100 nM, and Hoechst 34580 (Thermo Fisher Scientific, $H21486$) to $5\mu g/mL$ in growth media. For imaging, cells were incubated with $1mL$ of the staining solution for 30 minutes at 37 degree Celsius, rinsed five times with FluoroBrite DMEM (Thermo Fisher Scientific, $A1896701$), and subsequently imaged and analyzed in FluoroBrite DMEM.

***Multi-color structured illumination microscopy (SIM) super-resolution imaging***

A custom-built structured illumination microscope (SIM) was used for multi-color imaging1. Excitation wavelengths of 642nm (Spectra-Physics, Excelsior One 642), 532nm (Spectra-Physics, Millennia V), 488nm (Spectra-Physics, Excelsior One 488), and 405nm (Spectra-Physics, Excelsior One 405) were employed to excite Tubulin Tracker, CellMask Orange, MitoTracker, and Hoechst, respectively. The four laser beams were combined using three dichroic mirrors (Semrock, Di03-R405-t1; Semrock, Di03-R488-t1; Thorlabs, DMSP550R) and subsequently expanded by 4x. The combined beams were first diffracted into monochromatic beams by a blazed grating (Thorlabs, GR13-0605) and then recombined at the plane of a digital micromirror device (DMD; Texas Instruments, DLP9000X VIS WQXGA). The DMD-generated patterns were projected onto the sample plane through an objective lens (Nikon, SR Plan Apo, 60x, 1.27 WI). A multi-band dichroic mirror (Semrock, Di01-R405/488/532/635-25x36) and an emission filter (Semrock, FF01-446/510/581/703-25) were used to separate excitation and emission light. Additionally, a $2\times$ beam expander was employed to further magnify the emission signal, resulting in a total magnification of $120\times$. Fluorescence images were captured using an sCMOS detector (Photometrics, Kinetix).

For SIM super-resolution imaging, striped binary patterns with a second-order spatial frequency of $2.86\mu m^{-1}$ at the sample plane were displayed on the DMD. Patterns at three different angles with six phase shifts each were sequentially projected, with an exposure time of 100 ms per pattern. Super-resolution SIM reconstruction was performed using FairSIM 2, an ImageJ-based open-source software. Each raw image stack of $2048 \times 2048 \times 18$ was reconstructed into $4096 \times 4096$ super-resolution images, where each pixel corresponds to 27 nm.

#### F.7 The HHMI-D25 dataset

***Animal Experiments:***

Heterozygous PhAMexcised female mice carrying the Mito-Dendra2 transgene were generated through in-house breeding: first, PhAMexcised heterozygous males and females (strain #018397, Jackson Laboratories) were crossed to derive homozygous males, which were then bred with wild-type C57BL/6J females. Mice were housed in sound-attenuated, temperature- and humidity-controlled rooms under a 12-hour light/dark cycle, with food and water provided ad libitum. All procedures were conducted in accordance with NIH guidelines and were approved by the Institutional Animal Care and Use Committee (IACUC) at the Janelia Research Campus (Protocol #22-0229.04). Livers were collected via cardiac perfusion: first with $1\times$PBS to remove blood, followed by 30 mL of 4% paraformaldehyde (PFA) at a flow rate of 2.5 mL/min to minimize endothelial damage. Tissues were post-fixed in 4% PFA for 24 hours, rinsed three times in $1\times$ PBS, and stored in 1% PFA until further processing.

***Immunostaining and Image Acquisition:***

For imaging, samples were embedded in 4.6% low-melting-point agarose and sectioned into 120 $\mu$m slices using a Leica VT 1200S vibratome. Sections were blocked in 10% fetal bovine serum with 0.5% Triton X-100 for 1 hour and incubated with primary antibodies at 4°C for 48 hours. Mouse anti-PMP70 (MilliporeSigma, SAB4200181, 1:75) and rabbit anti-LAMP1 (Abcam, AB208943, 1:50) were used to label peroxisomes and lysosomes, respectively. Secondary antibodies — Alexa Fluor 647 goat anti-mouse (Thermo Fisher, A21235, 1:500) and Alexa Fluor 750 goat anti-rabbit (Thermo Fisher, A21039, 1:500) — were applied overnight at

4°C. Additional markers included Alexa Fluor Plus 555 Phalloidin (Thermo Fisher, A30106, 1:100) and HCS LipidTox Red (Thermo Fisher, H34476, 1:100) to label actin and lipid droplets, respectively. Nuclei were stained with DAPI (Thermo Fisher, D3571, 1:1000 dilution of 1 mg/mL stock) during a 25-minute PBS wash the following day. Sections were cleared using EasyIndex (LifeCanvas Technologies, EI-500-1.52) by incubating samples first in 50% EasyIndex for 1 hour, followed by 100% EasyIndex for 3–5 hours, and mounted using Secure-Seal spacers (Thermo Fisher, 0523073). Imaging was performed on a Leica Stellaris 8 confocal microscope using a 63×/1.40 NA oil immersion objective. Images were acquired at 2048 × 2048 pixels using bidirectional scanning, 2× optical zoom, and a pinhole size of 0.5 Airy units (AU). A total depth of 10$\mu$m was captured across 50 z-sections. Fluorophores were imaged using two acquisition lines: the first included mitochondria, actin, and lysosomes, while the second included nuclei, lipid droplets, and peroxisomes.

### G Further Details on all Experiments (Learning Tasks)

In this section, we mention specific details about individual tasks mentioned in the Table 1. Unless explicitly specified, we use the same hyperparameters for all tasks. Please refer to the code for details on the hyperparameters.

#### G.1 Two-channel semantic unmixing tasks

##### *Task I*

It is created from the HT-H24 dataset. It works on 3D z-stacks. The acquisition mode used in this dataset was *Training Mode I*. We show one qualitative example in Supplementary Figure S18.

##### *Task II*

It is created from HT-P23A dataset. It works on 3D z-stacks. The acquisition mode used in this dataset was *Training Mode I*. We show one qualitative example in Supplementary Figure S19.

##### *Task III*

It is created from the HT-P23B dataset. It works on 3D z-stacks. The acquisition mode used in this dataset was *Training Mode I*. We show one qualitative example in Supplementary Figure S20.

##### *Task IV-XII*

These tasks are created from the Pavia-P24 dataset. The acquisition mode used in them was *Training Mode III-a*. These tasks work on 2D frames. There are nine different acquisitions within this dataset that differ in (a) the laser power distribution among the two target channels and (b) the overall SNR. SNR has three levels, namely low, mid, and high, with low having the lowest SNR and high having the highest SNR level. The SNR level fixes the total combined laser power used for both channels. Within a single SNR level, we distribute the power among the two channels in three ways, namely, `50/50`, `66/33`, and `84/16` denoting the percentage laser power allocation to the respective channel. So, `84/16` means 84% of the total laser power was allocated to channel 1 and 16% was allocated to channel 2. We show one qualitative example per task in Supplementary Figures S27, S29, S28, S30, S31, S32, S33, S34, S35.

##### *Task XIII*

This task was created from the HT-T24 dataset. It works on 2D frames. The acquisition mode used in them was *Training Mode III-a*. We show one qualitative example in Supplementary Figure S36.

##### *Task XIV*

This task was created from the HT-LIF dataset. It used (Nucleus) and Microtubules as the two channels. It works on 2D frames. The acquisition mode used in them was *Training Mode III-a*. This task worked with the sub-dataset that had the exposure duration of 5$ms$. We show one qualitative example in Supplementary Figure S37.

***Task XV-XX***

These tasks were created from the Chicago-Sch23 dataset. The Chicago-Sch23 dataset has four structures, and these tasks capture all possible 2-channel semantic unmixing tasks from these four structures. These are Structured Illumination Microscopy (SIM) images, and due to the computational post-processing that happens in SIM, the resulting images have very different noise characteristics. Therefore, we disabled the use of noise models for all tasks generated from the Chicago-Sch23 dataset. More specifically, in Equation 2, we set $w = 0$. We show one qualitative example per task in Supplementary Figures S21, S22, S23, S24, S25, S26.

#### G.2 Three-channel semantic unmixing tasks

***Task XXI***

This task is a 3-channel semantic unmixing task generated from the CBG-Z18 dataset. This is a 3D dataset and so, we employ the 3D version of Micro$\mathbb{S}$plit. We show one qualitative example in Supplementary Figure S16.

***Task XXII***

This task is a 3-channel semantic unmixing task generated from the CBG-N18 dataset. This is a 3D dataset and so, we employ the 3D version of Micro$\mathbb{S}$plit. We show one qualitative example in Supplementary Figure S17.

***Task XXIII***

This task is a 3-channel 3D semantic unmixing task generated from HHMI-D25 dataset. Specifically, mitochondria, lysosomes and nuclei channels from HHMI-D25$_{8bit}$ sub-dataset is used to create this task. This is a 3D dataset and so, we employ 3D version of Micro$\mathbb{S}$plit. We show one qualitative example in Supplementary Figure S57.

***Tasks XXIV-XXVIII***

These tasks are generated from the HT-LIF24 dataset which has four structures. The three structures picked for these tasks are Nucleus, Microtubules and Kinetocore. While these tasks aim at splitting apart the above-mentioned three structures, they differ in the exposure duration of the training and evaluation dataset. The exposure duration used is mentioned in *Task Details* column of Table 1.

We show one qualitative example per task in Supplementary Figures S13, S14, S15, S11, S12.

***Task XXXI, XXXII***

These two tasks are 3-channel 3D semantic unmixing tasks generated from the denoised version of HHMI-D25$_{8bit}$ dataset, with denoising done by Noise2Void [4]. Mitochondria, lysosomes and nuclei channels from HHMI-D25$_{8bit}$ sub-dataset are used. This is a 3D dataset and so, we employ 3D version of Micro$\mathbb{S}$plit. While the task XXXI works directly on the above-mentioned data whereas task XXXII adds Gaussian and Poisson noise on top of the individual channels. Please refer to Supplementary sub-section B.1.1 on how the noise was added. On a technical node, since the task XXXI directly works on the denoised data, there is no rationale to use noise models on such data. Hence the denoi$\mathbb{S}$plit loss component is completely disabled ($w = 0$ in Equation 1) and the Task XXXI is effectively trained with $\mu$Split configuration. We show one qualitative example for each task in Supplementary Figure S61.

***Task XXXIII-XXXV***

These three tasks are 3-channel 3D semantic unmixing tasks generated from HHMI-D25$_{16bit}$ dataset. Mitochondria, lysosomes and nuclei channels from HHMI-D25$_{16bit}$ sub-dataset are used. This is a 3D dataset and so, we employ 3D version of Micro$\mathbb{S}$plit. While the task XXXIII works directly on the above-mentioned data whereas tasks XXXIV and XXXV adds Gaussian and Poisson noise on top of the individual channels. Please refer to Supplementary sub-section B.1.1 on how the noise was added. We show one qualitative example for each task in Supplementary Figure S62.

***Task XXXVI***

This task is a 3-channel 3D semantic unmixing task generated from HHMI-D25$_{16bit,0.25}$ dataset. Mitochondria, lysosomes and nuclei channels from HHMI-D25$_{16bit,0.25}$ sub-dataset are used. This is a 3D dataset and

| Training Mode | Correlation between structures preserved? | Input data is… | Note |
|---|---|---|---|
| *Training Mode I* | Yes | …mixed computationally. | Existing imaging data can be used (no special acquisitions required). |
| *Training Mode II* | No | …mixed computationally. | Existing imaging data can be used (no special acquisitions required). Results might be of lesser quality if reliable structural correlations between channels to be unmixed do exist. |
| *Training Mode III* | Yes | …imaged directly. | Mixed input must directly be imaged along with the individual target channels. Can lead to best performance when imaging is done well, but requires additional microscopy work. |

**Table ST11 Overview of Training Modes:** Properties and key advantages/ disadvantages of the training modes we propose. We say that the correlation between structures is preserved when their relative positioning in the computationally mixed input maintains their biologically occurring relative positioning in the sample.

so, we employ 3D version of Micro𝕊plit. We show one qualitative example for the task in Supplementary Figure S59.

#### G.3 Four-channel semantic unmixing tasks

##### *Task XXIX*

This task is generated from the HT-LIF24 dataset and uses all four structures to create this task. The exposure duration used for this task is $5ms$. We show one qualitative example per task in Figure 1f.

##### *Task XXX*

This task is generated from the Chicago-Sch23 dataset and uses all four structures to create this task. Due to the same reason as described for Tasks `XV-XX`, we have disabled the Noise model for this task as well. We show one qualitative example in Supplementary Figure S10.

| Task Idx | Dataset | Task Details | 2D/3D | PSNR | | MicroMS-SSIM | |
|---|---|---|---|---|---|---|---|
| | | | | C1 | C2 | C1 | C2 |
| I | HT-H24 | - | 3D | 0.09 | 0.55 | 0.973 | 0.956 |
| II | HT-P23A | - | 3D | 0.53 | 0.45 | 0.016 | 0.006 |
| III | HT-P23B | - | 3D | 0.29 | 0.39 | 0.005 | 0.019 |
| IV | Pavia-P24 | high, 50/50 | 2D | 0.17 | 1.1 | 0.010 | 0.009 |
| V | Pavia-P24 | high, 66/33 | 2D | 1.83 | 0.19 | 0.045 | 0.002 |
| VI | Pavia-P24 | high, 84/16 | 2D | 0.35 | 1.44 | 0.016 | 0.049 |
| VII | Pavia-P24 | mid, 50/50 | 2D | 0.26 | 0.55 | 0.005 | 0.013 |
| VIII | Pavia-P24 | mid, 66/33 | 2D | 0.25 | 0.09 | 0.009 | 0.003 |
| IX | Pavia-P24 | mid,84/16 | 2D | 0.02 | 0.42 | 0.005 | 0.006 |
| X | Pavia-P24 | low, 50/50 | 2D | 0.41 | 0.42 | 0.010 | 0.012 |
| XI | Pavia-P24 | low, 66/33 | 2D | 0.02 | 0.28 | 0.006 | 0.006 |
| XII | Pavia-P24 | low, 84/16 | 2D | 0.72 | 0.76 | 0.025 | 0.018 |
| XIII | HT-T24 | - | 2D | 0.66 | 0.52 | 0.005 | 0.004 |
| XIV | HT-LIF24 | - | 2D | 0.66 | 1.21 | 0.003 | 0.004 |
| XV | Chicago-Sch23 | C0 vs C1 | 2D | 1.80 | 0.92 | 0.003 | 0.001 |
| XVI | Chicago-Sch23 | C0 vs C2 | 2D | 1.68 | 1.32 | 0.005 | 0.002 |
| XVII | Chicago-Sch23 | C0 vs C3 | 2D | 1.20 | 0.54 | 0.000 | 0.000 |
| XVIII | Chicago-Sch23 | C1 vs C2 | 2D | 0.89 | 1.46 | 0.001 | 0.001 |
| XIX | Chicago-Sch23 | C1 vs C3 | 2D | 0.99 | 0.81 | 0.000 | 0.001 |
| XX | Chicago-Sch23 | C2 vs C3 | 2D | 1.01 | 0.82 | 0.000 | 0.000 |

| Task Idx | Dataset | Task Details | 2D/3D | PSNR | | | MicroMS-SSIM | | |
|---|---|---|---|---|---|---|---|---|---|
| | | | | C1 | C2 | C3 | C1 | C2 | C3 |
| XXI | CBG-Z18 | - | 3D | 0.12 | 0.18 | 0.12 | 0.002 | 0.001 | 0.001 |
| XXII | CBG-N18 | - | 3D | 0.18 | 0.21 | 0.06 | 0.000 | 0.000 | 0.000 |
| XXIII | HHMI-D25$_{8bit}$ | - | 3D | 0.018 | 0.035 | 0.027 | 0.000 | 0.002 | 0.001 |
| XXIV | HT-LIF24 | 2ms | 2D | 1.19 | 0.90 | 0.66 | 0.007 | 0.003 | 0.004 |
| XXV | HT-LIF24 | 3ms | 2D | 1.30 | 0.96 | 0.75 | 0.008 | 0.003 | 0.004 |
| XXVI | HT-LIF24 | 5ms | 2D | 1.11 | 0.89 | 0.53 | 0.004 | 0.002 | 0.004 |
| XXVII | HT-LIF24 | 20ms | 2D | 0.85 | 0.62 | 0.73 | 0.002 | 0.001 | 0.002 |
| XXVIII | HT-LIF24 | 500ms | 2D | 0.61 | 0.35 | 0.66 | 0.001 | 0.001 | 0.001 |
| XXXI | HHMI-D25$_{8bit,denoised}$ | - | 3D | 0.073 | 0.11 | 0.151 | 0.000 | 0.001 | 0.001 |
| XXXII | HHMI-D25$_{8bit,denoised}$ | $\sigma = 20, \lambda = 30$ | 3D | 0.076 | 0.094 | 0.153 | 0.000 | 0.003 | 0.002 |
| XXXIII | HHMI-D25$_{16bit}$ | - | 3D | 0.03 | 0.126 | 0.016 | 0.002 | 0.003 | 0.001 |
| XXXIV | HHMI-D25$_{16bit}$ | $\sigma = 2K, \lambda = 5K$ | 3D | 0.031 | 0.13 | 0.014 | 0.002 | 0.003 | 0.001 |
| XXXV | HHMI-D25$_{16bit}$ | $\sigma = 4K, \lambda = 10K$ | 3D | 0.033 | 0.133 | 0.016 | 0.002 | 0.004 | 0.001 |
| XXXVI | HHMI-D25$_{16bit,0.25}$ | - | 3D | 0.082 | 0.103 | 0.021 | 0.000 | 0.001 | 0.000 |

| Task Idx | Dataset | 2D/3D | PSNR | | | | MicroMS-SSIM | | | |
|---|---|---|---|---|---|---|---|---|---|---|
| | | | C1 | C2 | C3 | C4 | C1 | C2 | C3 | C4 |
| XXIX | HT-LIF24 | 2D | 0.78 | 1.05 | 0.86 | 0.53 | 0.004 | 0.004 | 0.002 | 0.006 |
| XXX | Chicago-Sch23 | 2D | 1.57 | 0.87 | 1.24 | 0.65 | 0.006 | 0.002 | 0.003 | 0.008 |

**Table ST12** Standard error values for all entries of Table 1.

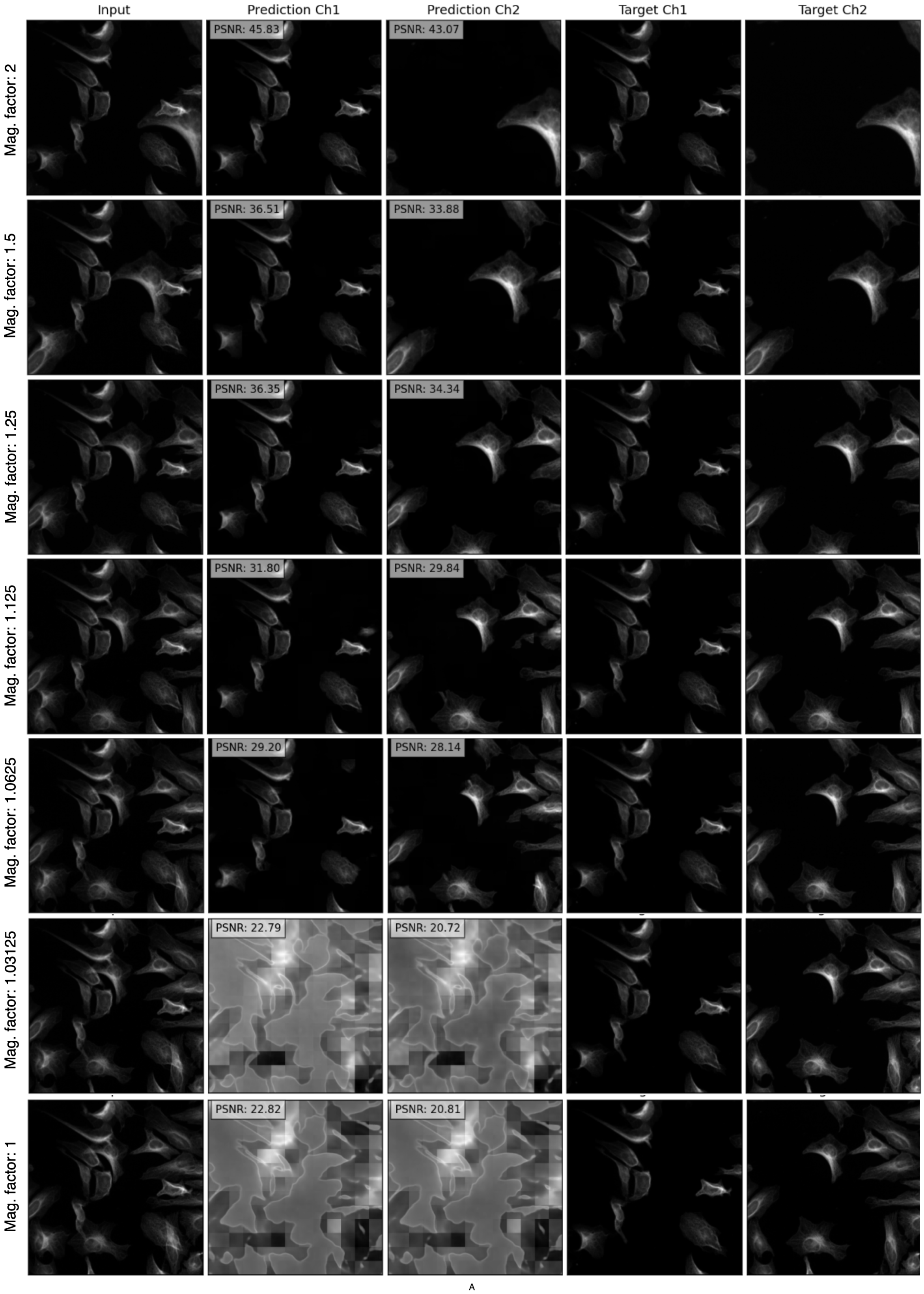


**Fig. S2 Semantic unmixing of similar structures with Micro𝕊plit.** We show how the same structure in two target channels can still be unmixed, even if the only structural difference is a controllable scaling factor. See Supplementary Section B.3 for a detailed description of the conducted experiments.

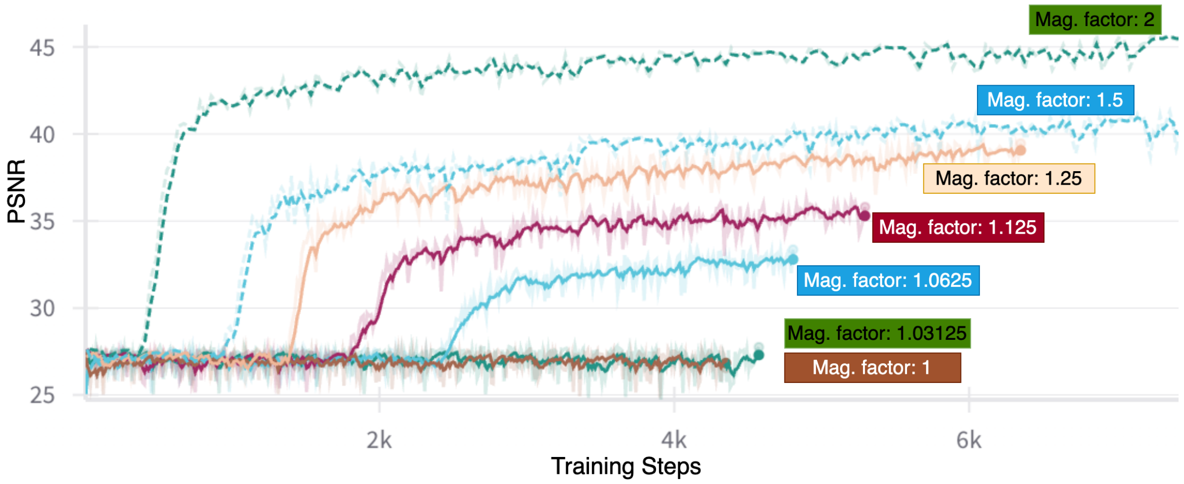


**Fig. S3 Semantic unmixing of similar structures with Micro𝕊plit.** Here we plot *Validation PSNR vs. training steps* for all experiments shown in Figure S2. Observe that, as the magnification factor gets closer to 1, it takes longer for the network to initiate the learning, and the quality of the splitting converges to an overall lower level. See Supplementary Section B.3 for a detailed description of the conducted experiments.

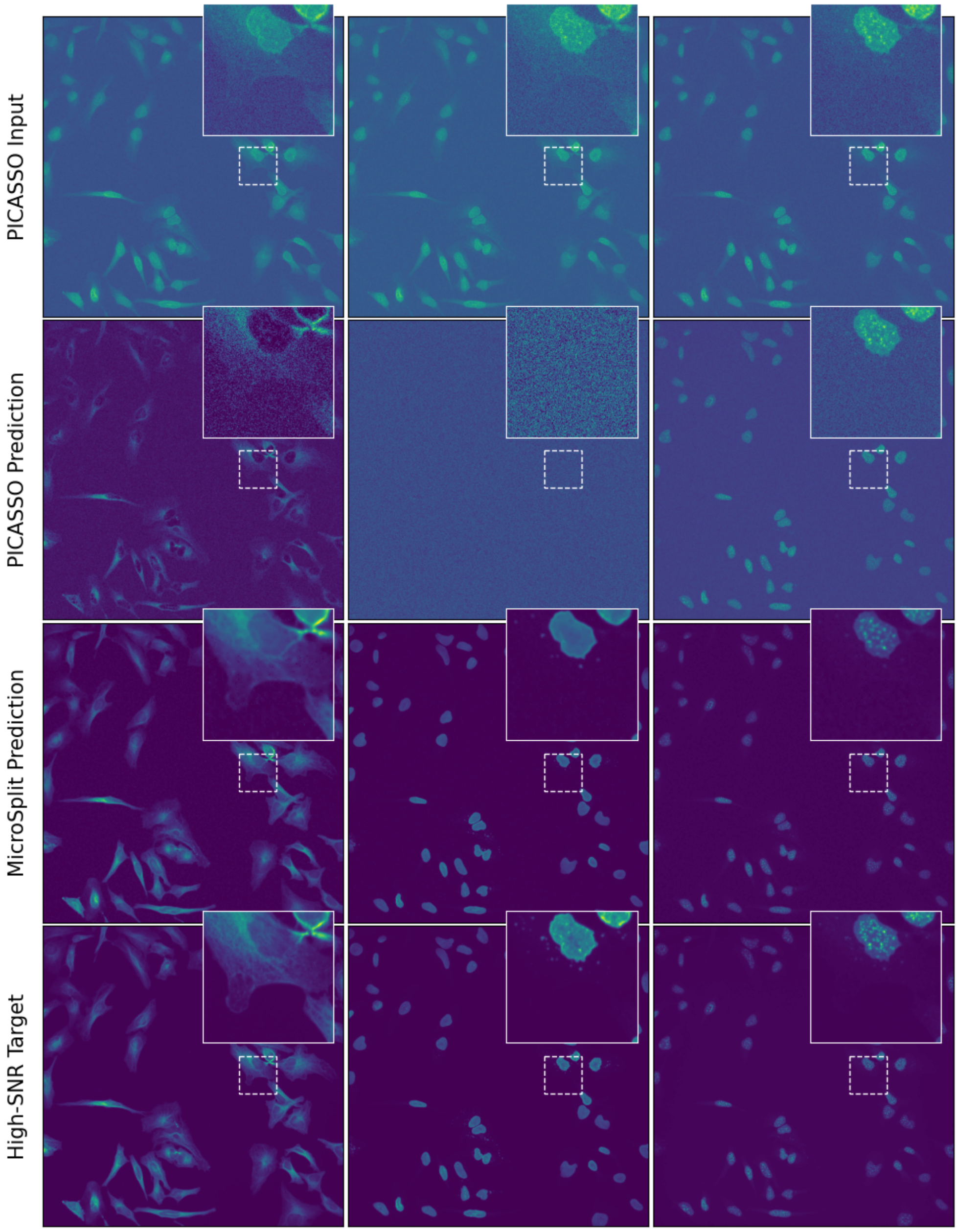


**Fig. S4 Comparison with PICASSO** on a 3-channel unmixing task. Please note that Micro𝕊plit distinguishes itself from PICASSO [20] by only requiring a single superimposed image as input, rather than multiple images with different spectral overlaps. Here, we used the high-SNR data from the HT-LIF24 dataset, prepared inputs that are suitable for PICASSO, and added additional (synthetic) noise to demonstrate how different both systems deal with noisy data. In row 1, we show example images from the the three input channels used for PICASSO. In rows 2 and 3, we show prediction by PICASSO and Micro𝕊plit, respectively. In the last row, we show the high-SNR ground truth for visual evaluation. We observe that, in the presence of noise, Picasso starts to be challenged to unmix the data, mostly if it starts having similar features (see columns 2 and 3, where PICASSO removes data around the nucleus region seen in column 1.

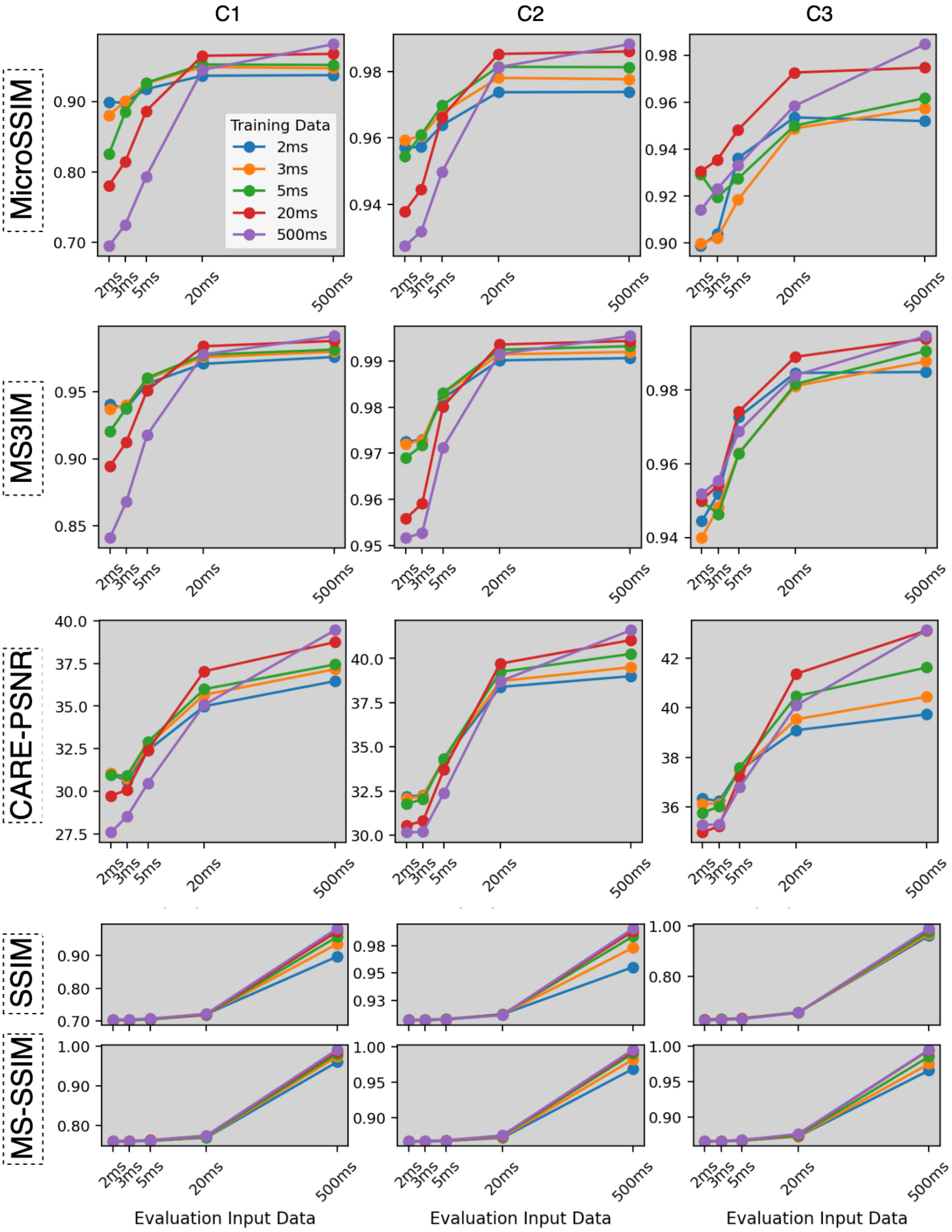


**Fig. S5 Quantitative evaluations of the effect of signal-to-noise ratios (SNR) on in-distribution and out-of-distribution unmixing results (using the HT-LIF24 dataset).** We trained MicroSplit models on micrographs acquired using a range of different exposure times. Note that the underlying sample ROIs remain identical. We then used all trained models (for 2ms, 3ms, 5ms, 20ms and 500ms exposure time data, indicated in the legend) and evaluated their semantic unmixing performance on all exposure times (x-axis), respectively. The rows show unmixed results quantified using MicroSSIM, MicroMS-SSIM, CARE-PSNR, SSIM and MS-SSIM metrics. All plots utilize the legend presented in the plot in the first row, first column. Note that the plots in this figure also demonstrate that the MS-SSIM and SSIM metrics do not work well on microscopy data (while MicroSSIM and MicroMS-SSIM show better sensitivity [8]).

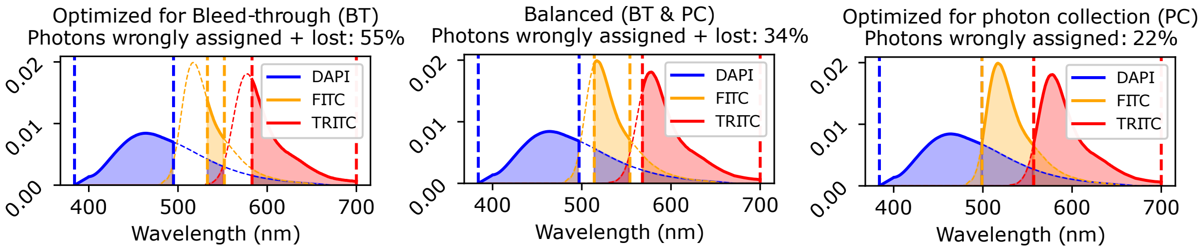


| | GT Noisy vs GT High-SNR | | | | | | Prediction vs GT High-SNR | | | | | |
|---|---|---|---|---|---|---|---|---|---|---|---|---|
| Acq. | PSNR | | | MicroMS-SSIM | | | PSNR | | | MicroMS-SSIM | | |
| Duration | C1 | C2 | C3 | C1 | C2 | C3 | C1 | C2 | C3 | C1 | C2 | C3 |
| 2ms | 23.3 | 25.1 | 26.1 | 0.839 | 0.772 | 0.869 | 31.0 | 32.2 | 36.3 | 0.940 | 0.973 | 0.944 |
| 3ms | 23.6 | 26.0 | 27.2 | 0.842 | 0.780 | 0.871 | 30.8 | 32.2 | 36.1 | 0.940 | 0.973 | 0.948 |
| 5ms | 24.6 | 28.3 | 29.9 | 0.857 | 0.817 | 0.875 | 32.9 | 34.3 | 37.6 | 0.960 | 0.983 | 0.963 |
| 20ms | 30.0 | 35.6 | 38.16 | 0.920 | 0.942 | 0.914 | 37.0 | 39.7 | 41.4 | 0.984 | 0.994 | 0.989 |

**Fig. S6 Photon efficient imaging with Micro𝕊plit: a concrete example.** We quantify two mechanisms through which Micro𝕊plit reduces the required photon budget: (*i*) *Reduced photon filtering* ***(top):*** Emission spectra of the three fluorophores DAPI, FITC, and TRITC are shown together with example emission filter bands used in conventional multi-color imaging. From left to right, we illustrate three settings that increasingly trade off bleed-through (BT) against photon efficiency: a configuration that strongly suppresses BT, a balanced configuration that collects more photons at the cost of some BT, and a highly permissive configuration that captures nearly all emitted photons but suffers notable spectral overlap. Micro𝕊plit, by contrast, allows imaging multiple structures within a single broad emission band and subsequently reassigning the collected intensities to their respective output channels leading to the indicated photon-efficiency increases. (*ii*)*Denoising* ***(bottom):*** Denoising enables repurposing of the available photon budget by acquiring lower-SNR micrographs, which Micro𝕊plit can restore to high-SNR predictions. We compare raw data acquired at 2ms, 3ms, 5ms, and 20ms exposure times with high-SNR (500ms) reference images of the same regions in the HT-LIF24 dataset (three channels: Nucleus, Microtubules, Kinetochore). The first six data columns in the table quantify the similarity of low-exposure raw data to the 500ms reference, while the rightmost six columns show the corresponding similarities for Micro𝕊plit predictions (identical data as in Table1). In all cases, the Micro𝕊plit predictions exhibit higher quality than the corresponding raw inputs. Strikingly, predictions from 2ms exposures already surpass the quality of 5ms raw data for all channels, and for Channel 1 even exceed the 20ms raw data. In this example, this corresponds to at least a three-fold reduction in required photon budget per acquisition, and likely closer to an order of magnitude when averaged across the three channels.

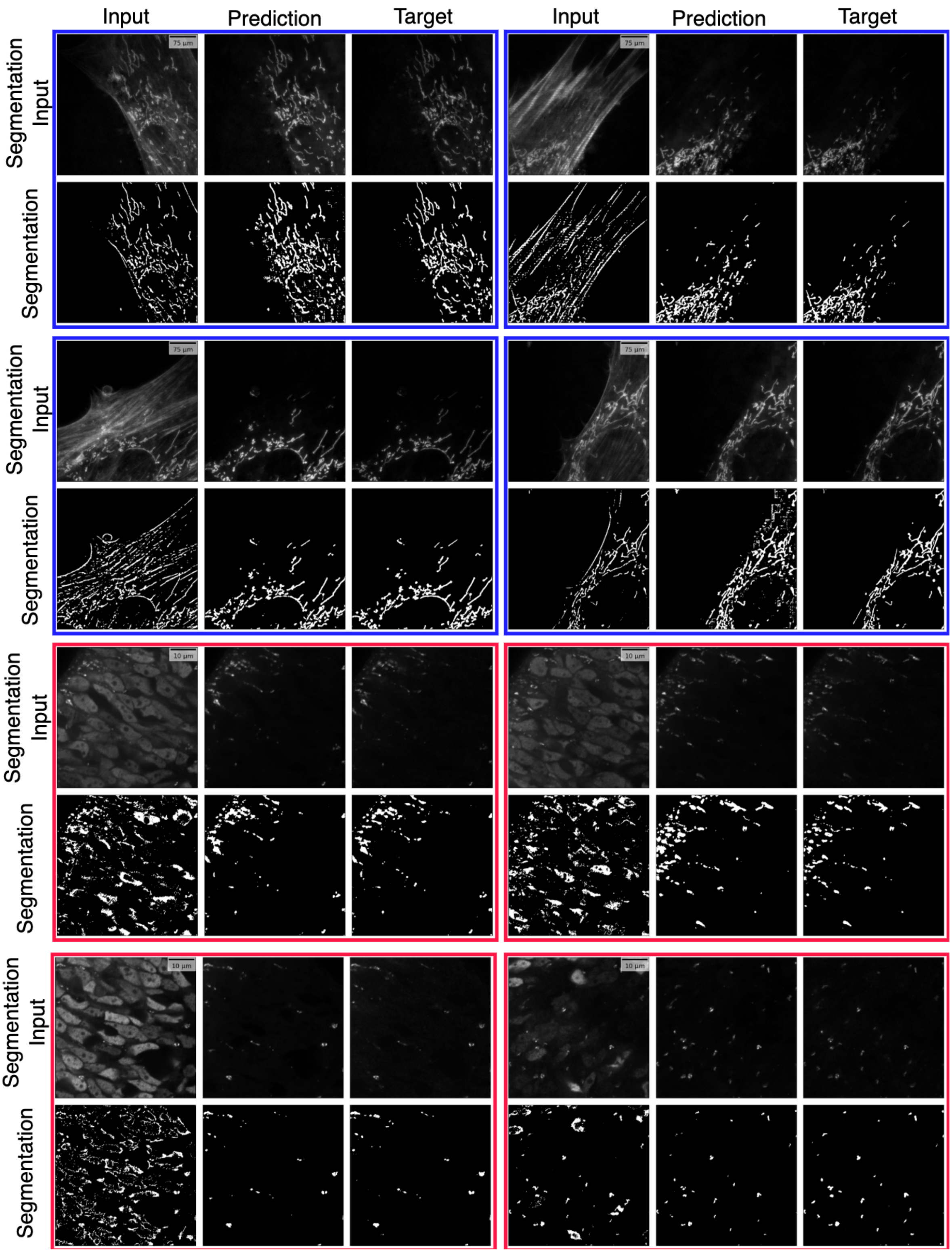


**Fig. S7 Segmentation results using common combined labels**. In Fig. 3, for training the segmentation model, analysts annotated the target images and trained the segmentation model on target images with those annotations. Similar steps were taken for prediction images. In this experiment, we combined the annotations of both target and prediction images and trained three models, one for target images, one for prediction images, and one for the input images. The purpose for training the segmentation model on the input images was to ascertain whether one can segment one structure from the superimposed input itself. The motivation to use common annotations for training all segmentation models using common annotations was to eliminate the situation where annotation for either target or prediction images were imperfect and hence the segmentation results became inferior to what they should otherwise be.

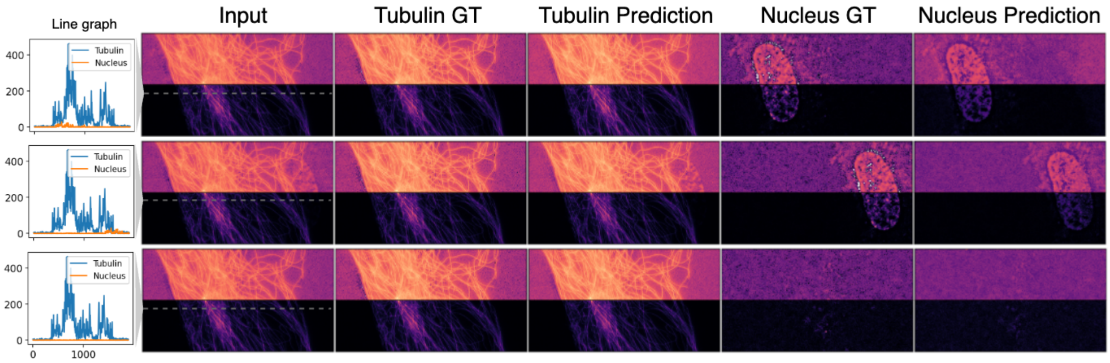


**Fig. S8 Unmixing unequally intense channels.** Here, we investigate what happens when one of the channels to be unmixed is much dimmer than the other (we called the divergence of structure intensities 'skew' in the main text). We used Tubulin and Nucleus channel from the 4 channel Chicago-Sch23 dataset. Line graphs of Tubulin and Nuclei shows that nucleus channel is considerably dimmer, such that the signal amplitude of this channel is only around the noise amplitude of the Tubulin channel. For every patch, we visualize half of the patch in *LogNorm* so that even the faintest signal becomes visible. We intend to answer the question whether the prediction of nucleus channel is mostly due to learned spatial correlation with the Tubulin channel or if the trained model is indeed capable to unmix such highly skewed intensities. To answer this question, we first trained the Micro𝕊plit network on the original data and then conducted three test-time experiments (showing one per row in the figure). For these experiments, we picked a location in test data which contained both nucleus and tubulin. Micro𝕊plit was training using *Training Mode I*. (*a*) We evaluate on the superimposed image patch created from the pre-selected location. (*b*) We tinkered with the inter-structure spatial correlations by shifting the nucleus channel relative to tubulin channel consistently by 1000 pixels. (*c*) We removed nuclei from the nucleus channel entirely by replacing them with background patches from the same channel. The results clearly indicate that Micro𝕊plit was able to unmix the dim nucleus signals, even if the spatial correlation with the tubulin channel was tinkered with (observe the last two columns, *i.e.* ground truth *vs.* prediction of the nucleus channel).

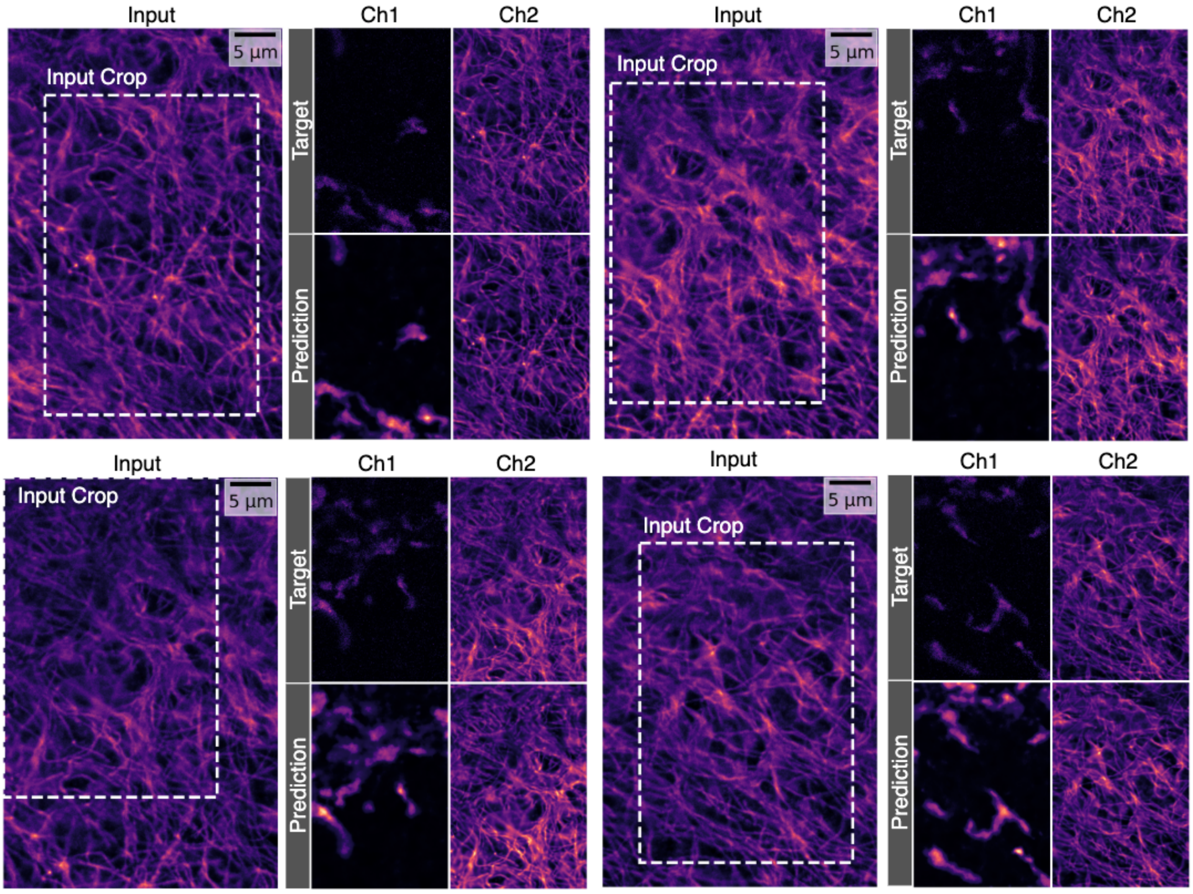


**Fig. S9 Task II, HT-P23A** This task has a high skew with Mitochondria channel (Ch1) being weaker than Microtubule channel (Ch2), so much so that it is difficult to locate the Mitochondria in the superimposed input. Micro𝕊plit MMSE predictions are subject to a certain 'blurryness' which indicates the model's inability to give a consistent prediction for the Mitochondria channel.

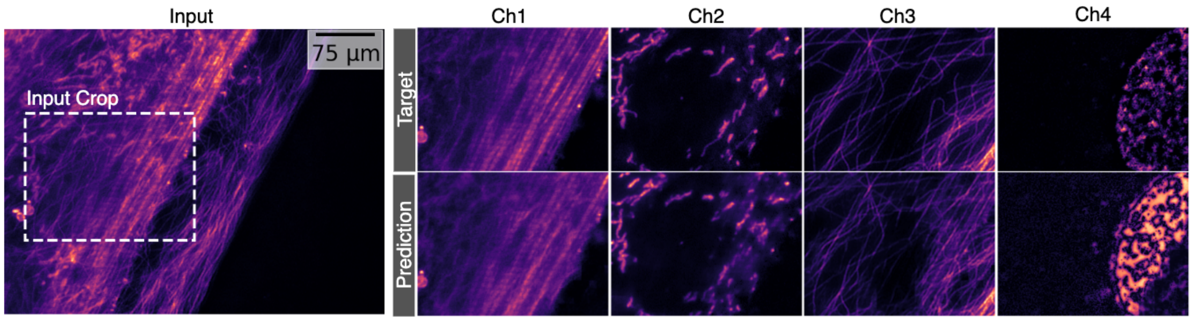


**Fig. S10** Qualitative Evaluation for Task XXX from Chicago-Sch23 dataset. Note that we show the target and the prediction corresponding to the input crop which is denoted in *Input* panel by a white dotted rectangle.

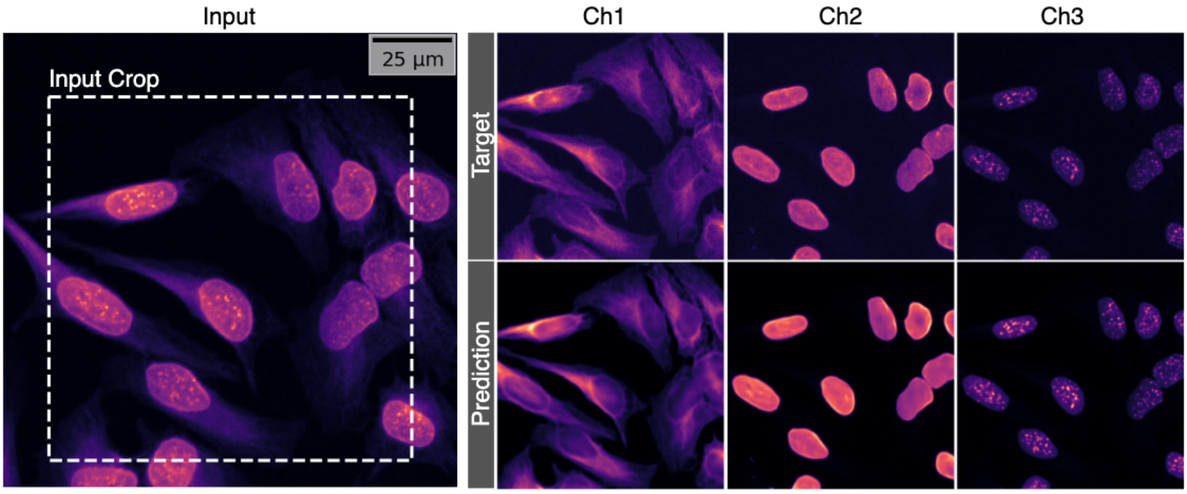


**Fig. S11** Qualitative Evaluation for Task XXVII from HT-LIF24 dataset. Note that we show the target and the prediction corresponding to the input crop which is denoted in *Input* panel by a white dotted rectangle.

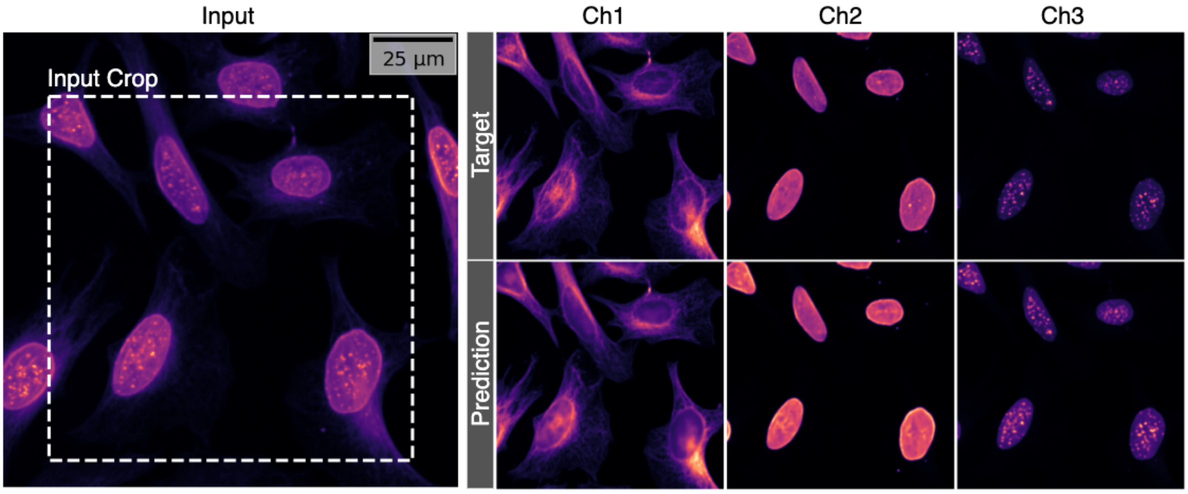


**Fig. S12** Qualitative Evaluation for Task XXVIII from HT-LIF24 dataset. Note that we show the target and the prediction corresponding to the input crop which is denoted in *Input* panel by a white dotted rectangle.

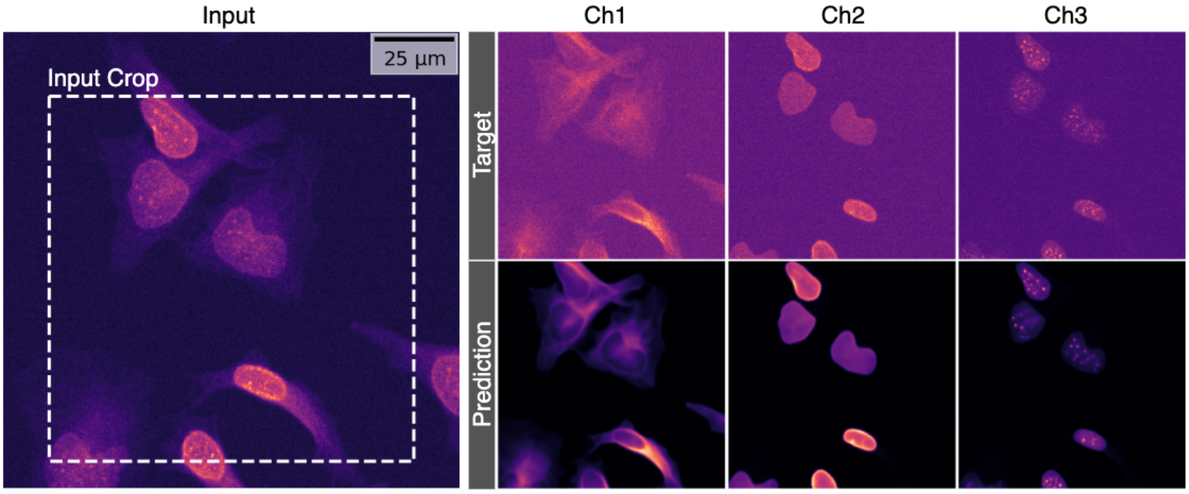


**Fig. S13** Qualitative Evaluation for Task XXIV from HT-LIF24 dataset. Note that we show the target and the prediction corresponding to the input crop which is denoted in *Input* panel by a white dotted rectangle.

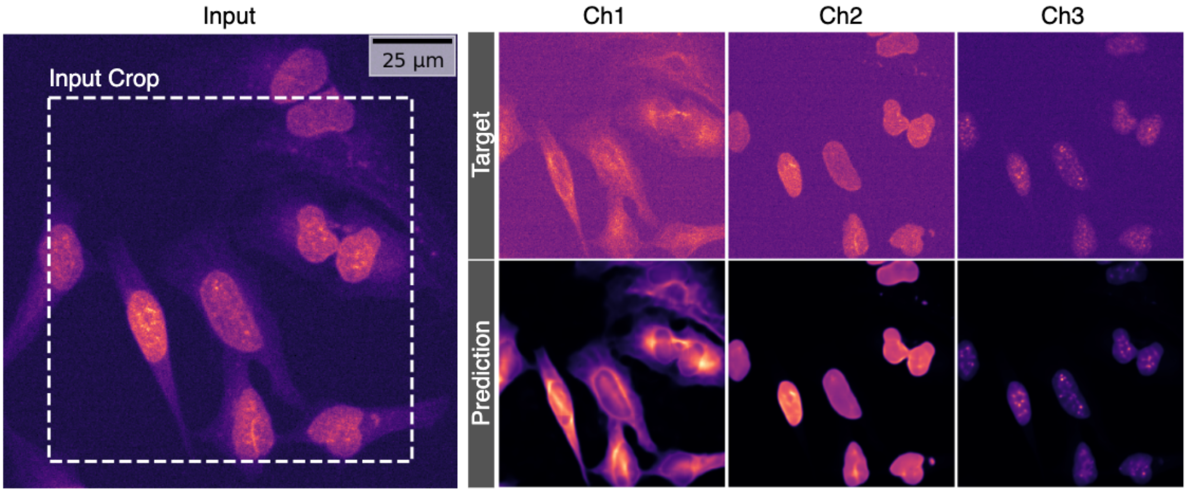


**Fig. S14** Qualitative Evaluation for Task XXV from HT-LIF24 dataset. Note that we show the target and the prediction corresponding to the input crop which is denoted in *Input* panel by a white dotted rectangle.

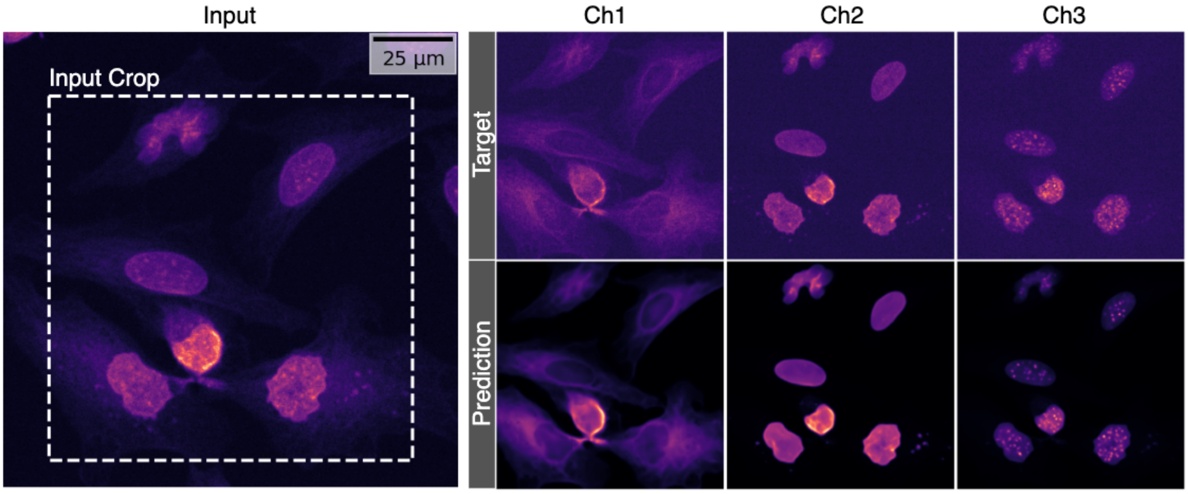


**Fig. S15** Qualitative Evaluation for Task XXVI from HT-LIF24 dataset. Note that we show the target and the prediction corresponding to the input crop which is denoted in *Input* panel by a white dotted rectangle.

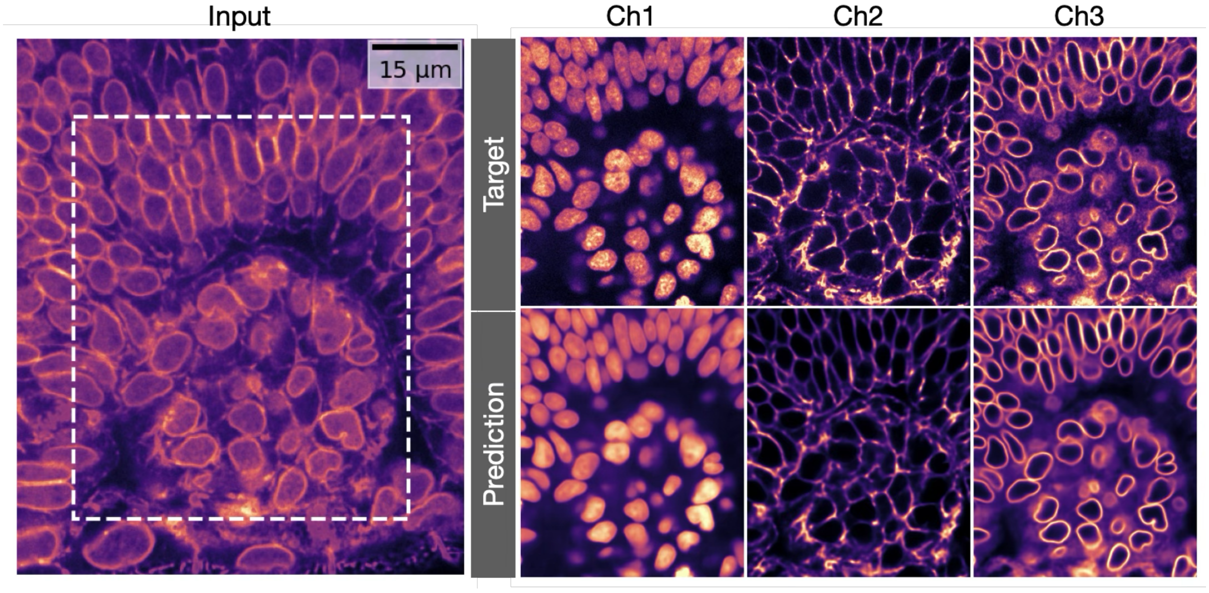


**Fig. S16** Qualitative Evaluation for Task XXI from CBG-Z18 dataset. Note that we show the target and the prediction corresponding to the input crop which is denoted in *Input* panel by a white dotted rectangle.

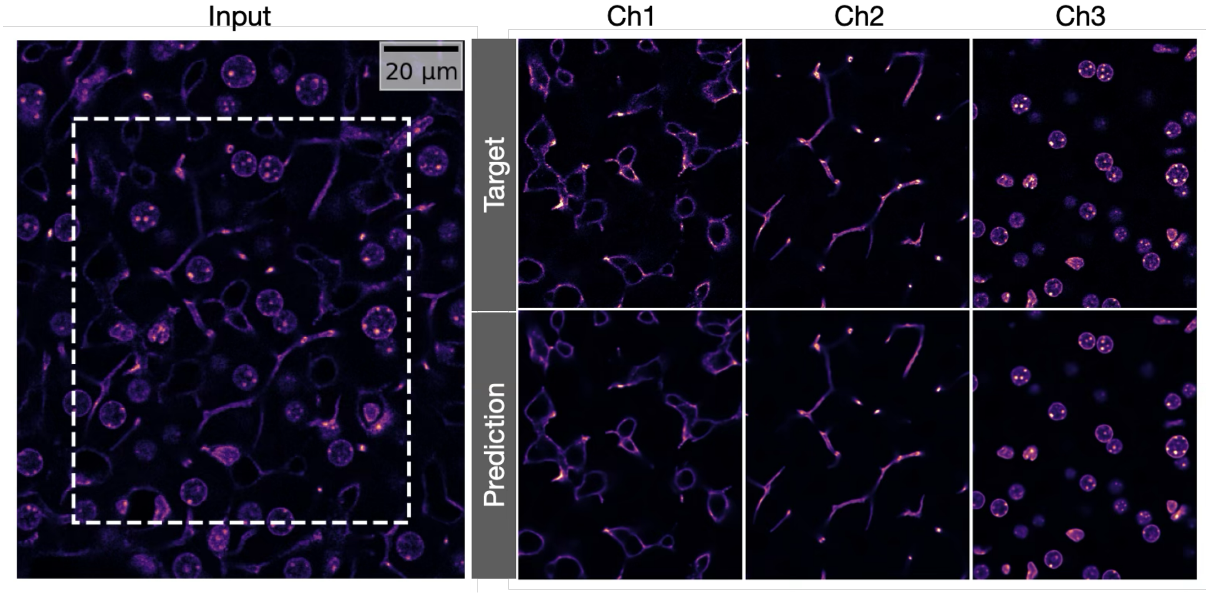


**Fig. S17** Qualitative Evaluation for Task XXII from CBG-N18 dataset. Note that we show the target and the prediction corresponding to the input crop which is denoted in *Input* panel by a white dotted rectangle.

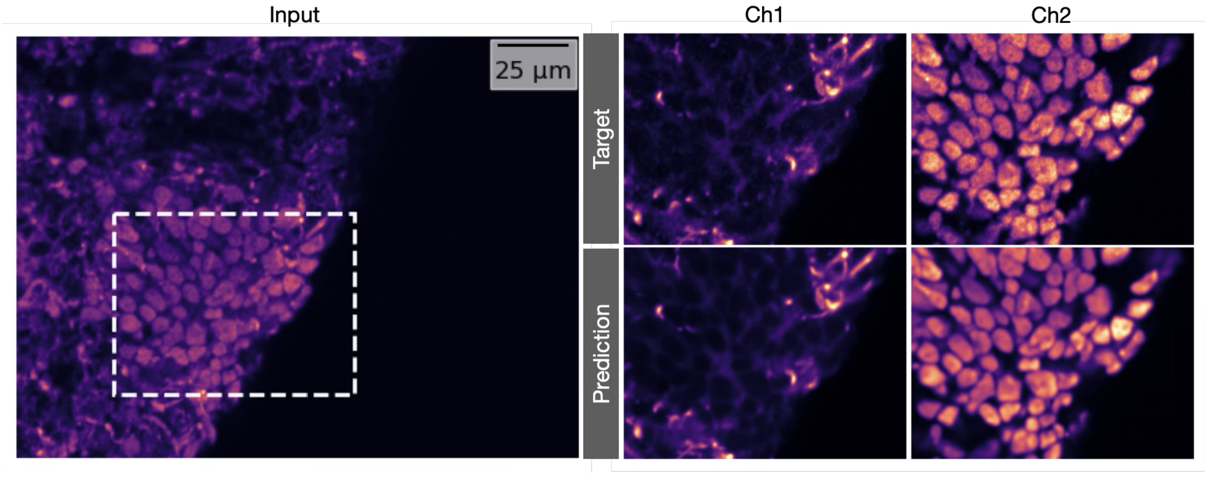


**Fig. S18** Qualitative Evaluation for Task I from HT-H24 dataset. Note that we show the target and the prediction corresponding to the input crop which is denoted in *Input* panel by a white dotted rectangle.

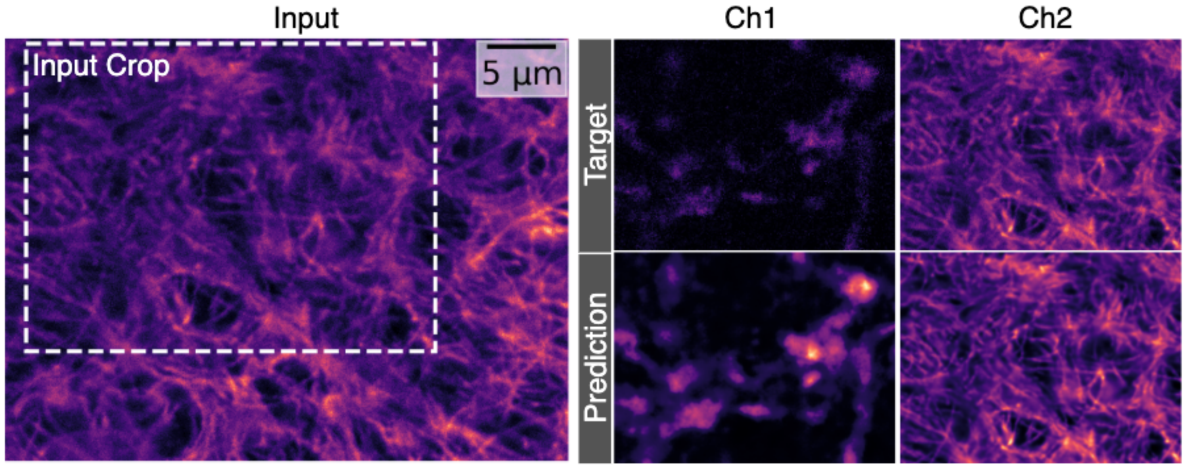


**Fig. S19** Qualitative Evaluation for Task II from HT-P23A dataset. Note that we show the target and the prediction corresponding to the input crop which is denoted in *Input* panel by a white dotted rectangle.

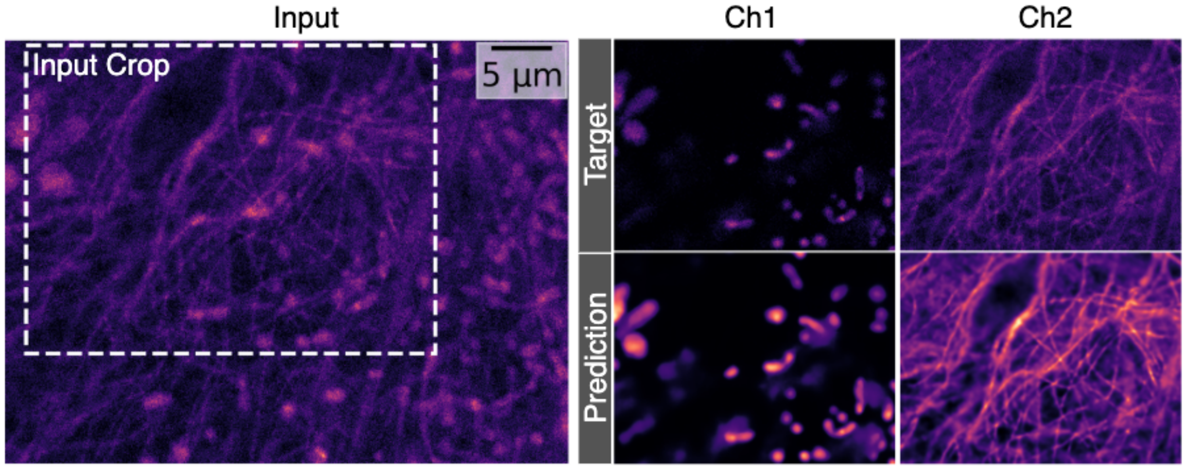


**Fig. S20** Qualitative Evaluation for Task III from HT-P23B dataset. Note that we show the target and the prediction corresponding to the input crop which is denoted in *Input* panel by a white dotted rectangle.

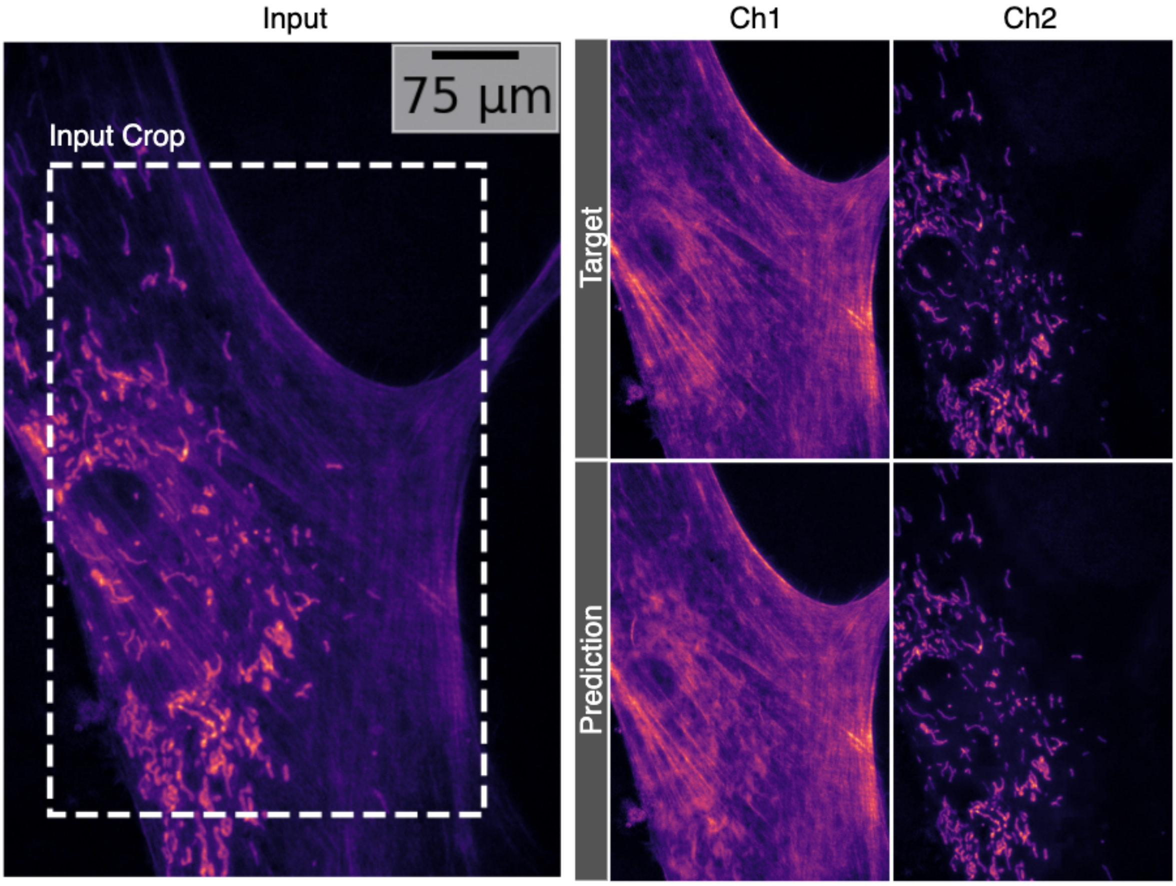


**Fig. S21** Qualitative Evaluation for Task XV from Chicago-Sch23 dataset. Note that we show the target and the prediction corresponding to the input crop which is denoted in *Input* panel by a white dotted rectangle.

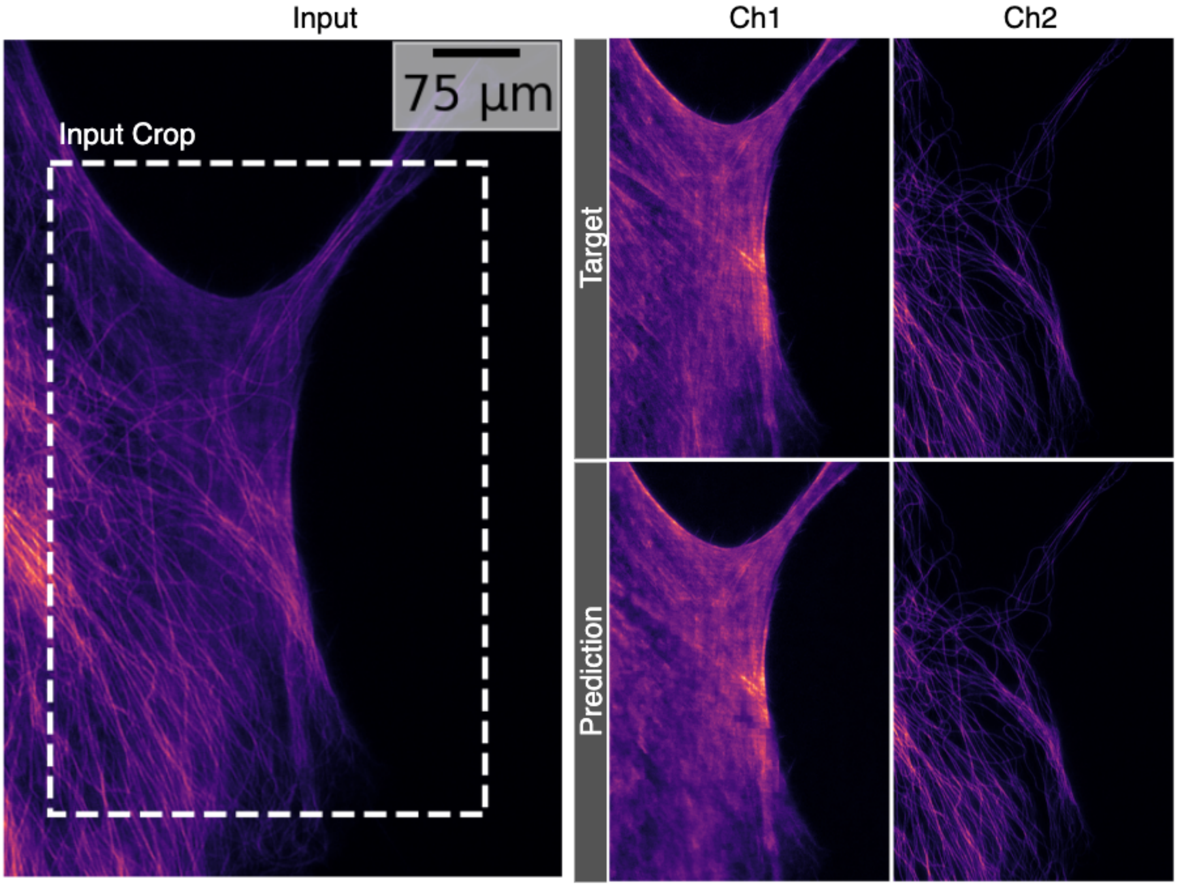


**Fig. S22** Qualitative Evaluation for Task XVI from Chicago-Sch23 dataset. Note that we show the target and the prediction corresponding to the input crop which is denoted in *Input* panel by a white dotted rectangle.

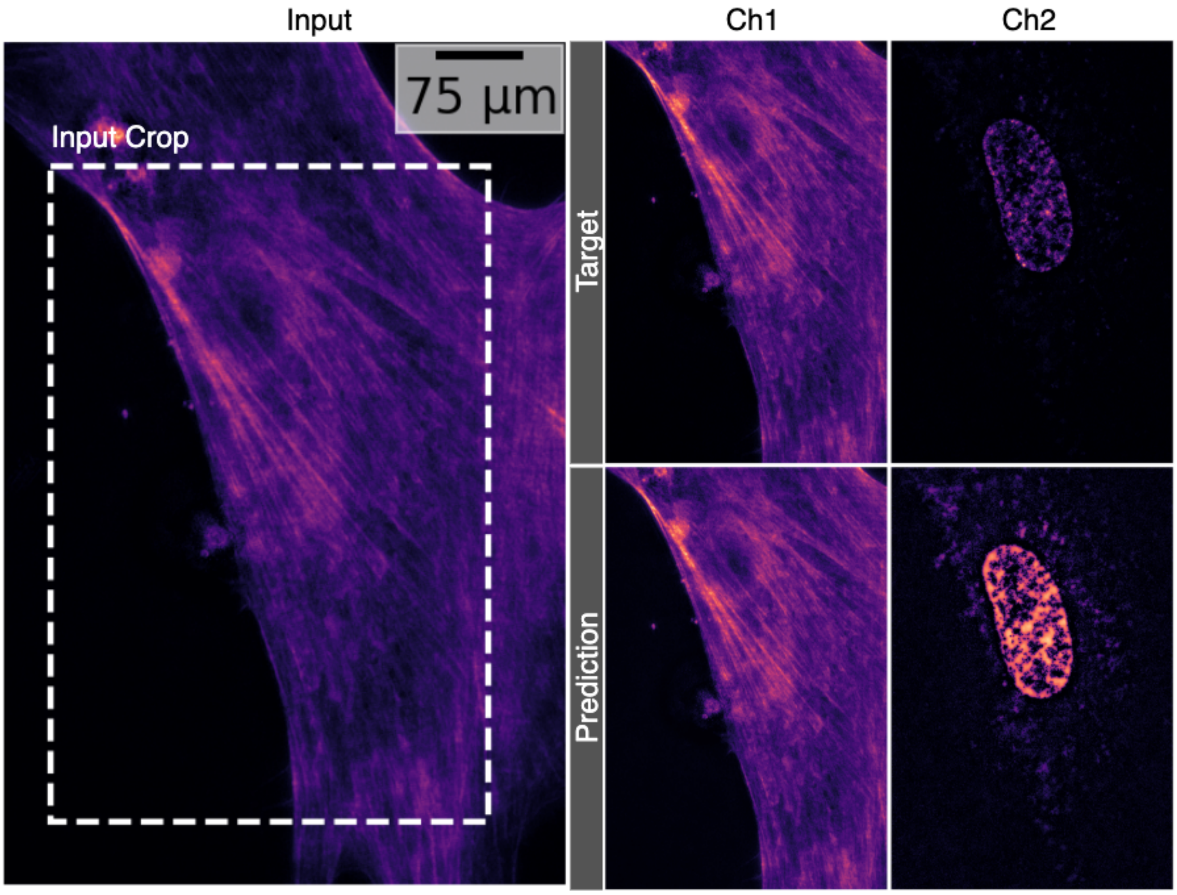


**Fig. S23** Qualitative Evaluation for Task XVII from Chicago-Sch23 dataset. Note that we show the target and the prediction corresponding to the input crop which is denoted in *Input* panel by a white dotted rectangle.

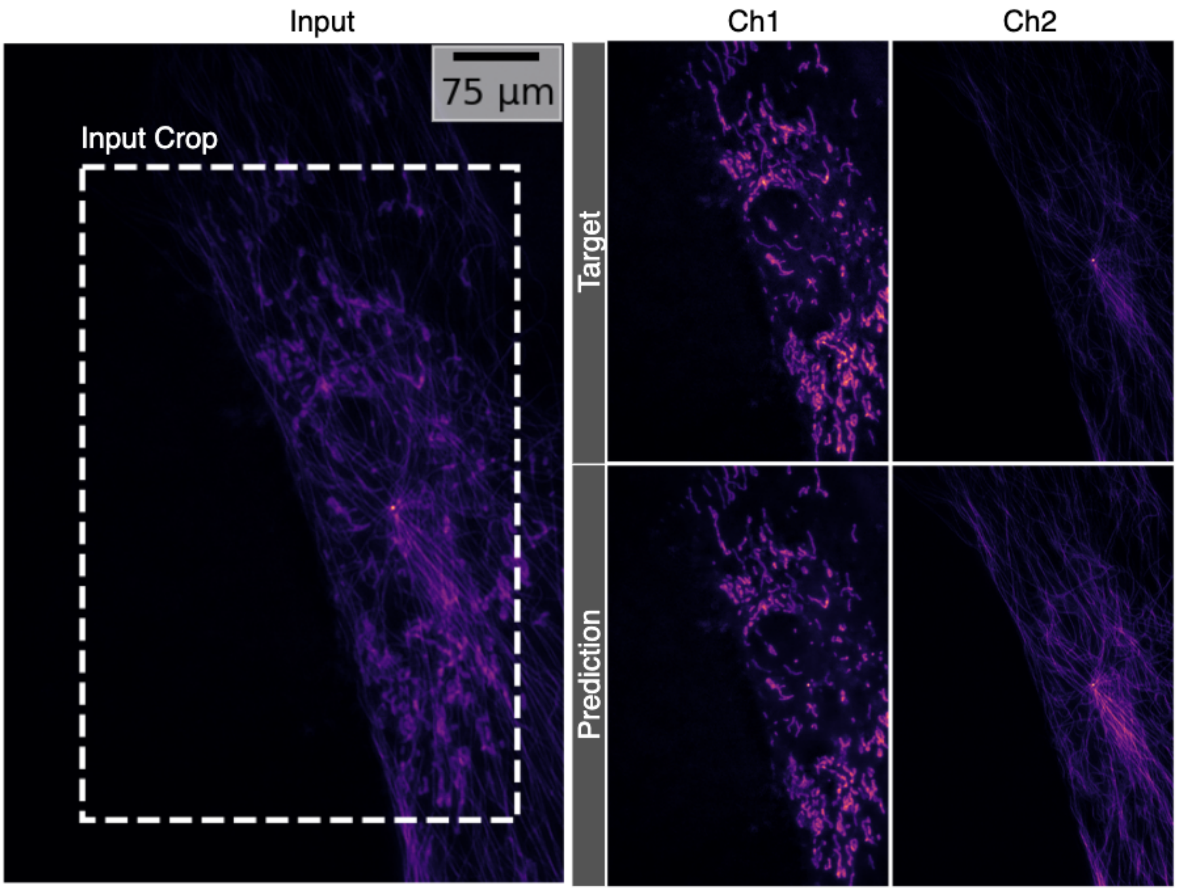


**Fig. S24** Qualitative Evaluation for Task XVIII from Chicago-Sch23 dataset. Note that we show the target and the prediction corresponding to the input crop which is denoted in *Input* panel by a white dotted rectangle.

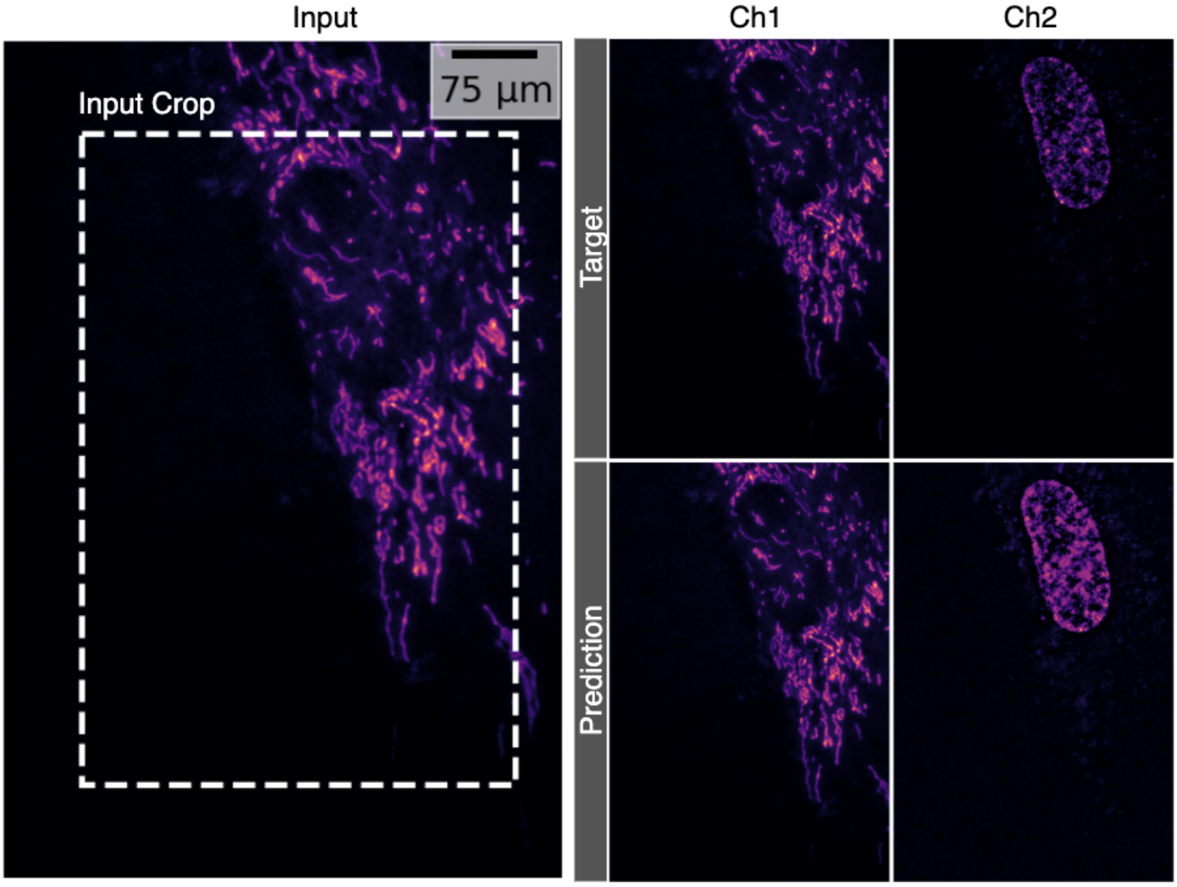


**Fig. S25** Qualitative Evaluation for Task XIX from Chicago-Sch23 dataset. Note that we show the target and the prediction corresponding to the input crop which is denoted in *Input* panel by a white dotted rectangle.

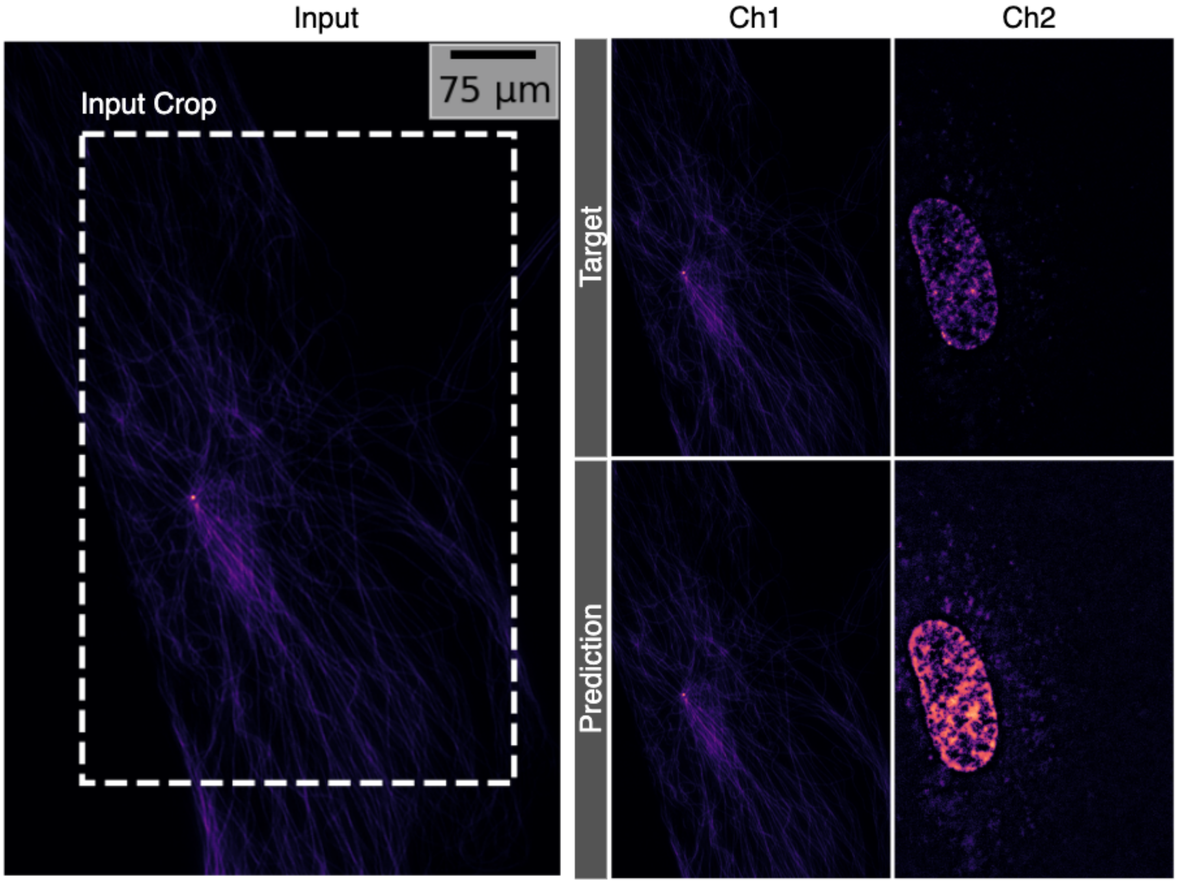


**Fig. S26** Qualitative Evaluation for Task XX from Chicago-Sch23 dataset. Note that we show the target and the prediction corresponding to the input crop which is denoted in *Input* panel by a white dotted rectangle.

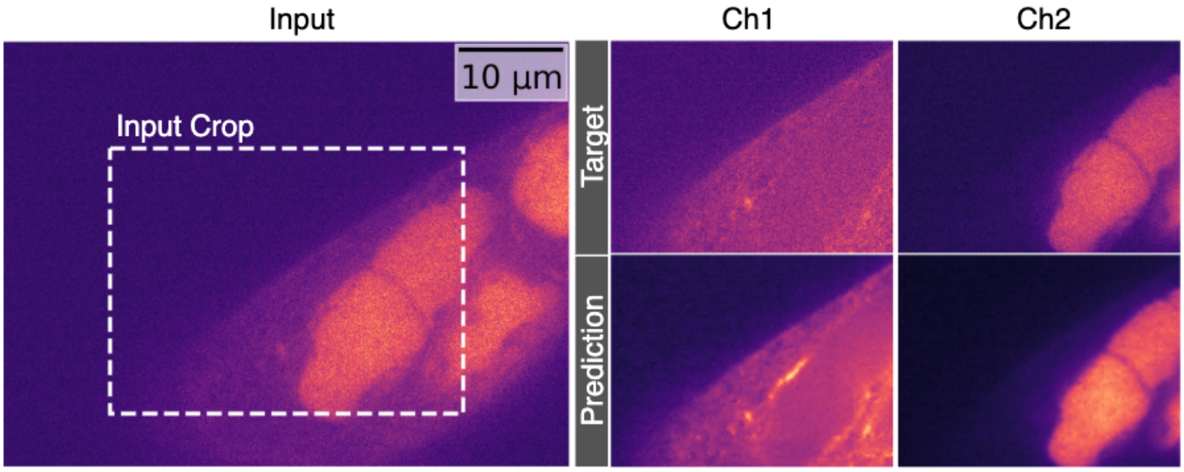


**Fig. S27** Qualitative Evaluation for Task IV from Pavia-P24 dataset. Note that we show the target and the prediction corresponding to the input crop which is denoted in *Input* panel by a white dotted rectangle.

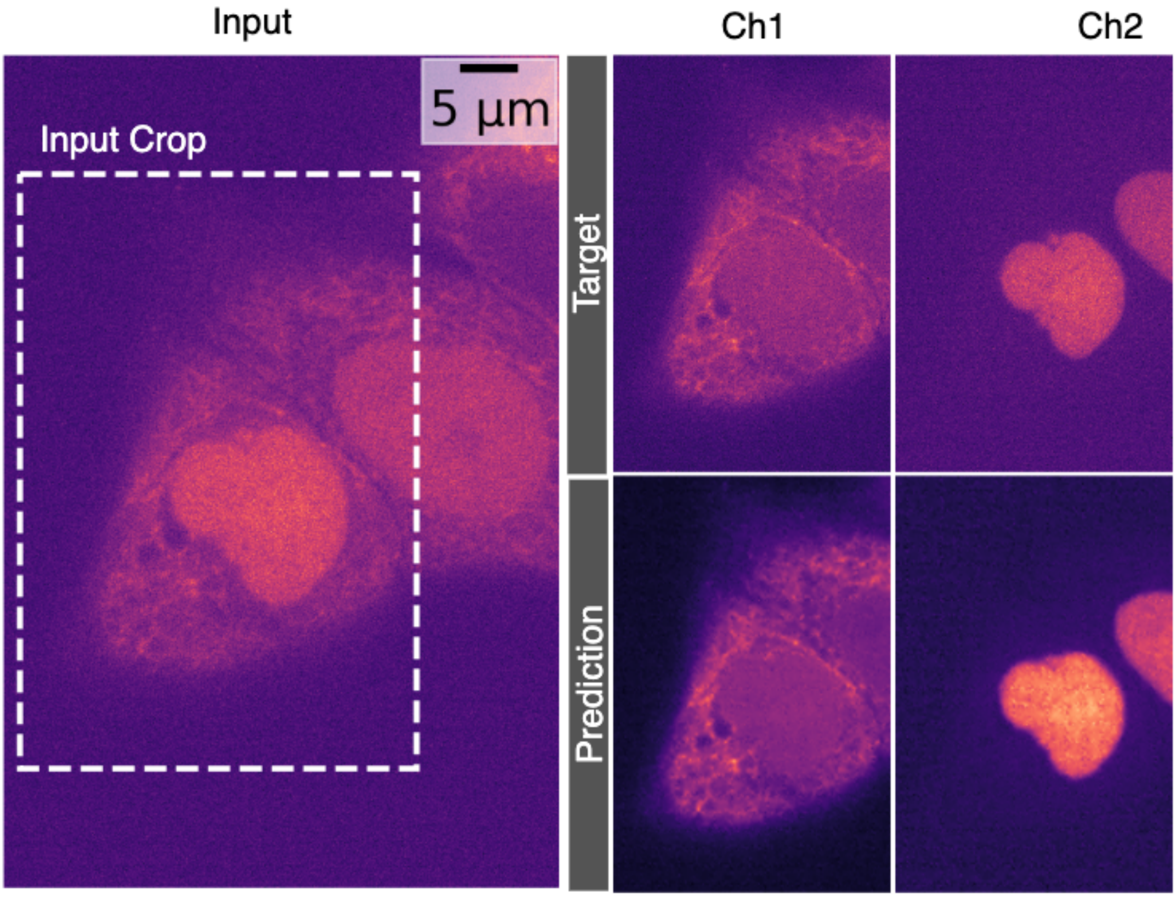


**Fig. S28** Qualitative Evaluation for Task VI from Pavia-P24 dataset. Note that we show the target and the prediction corresponding to the input crop which is denoted in *Input* panel by a white dotted rectangle.

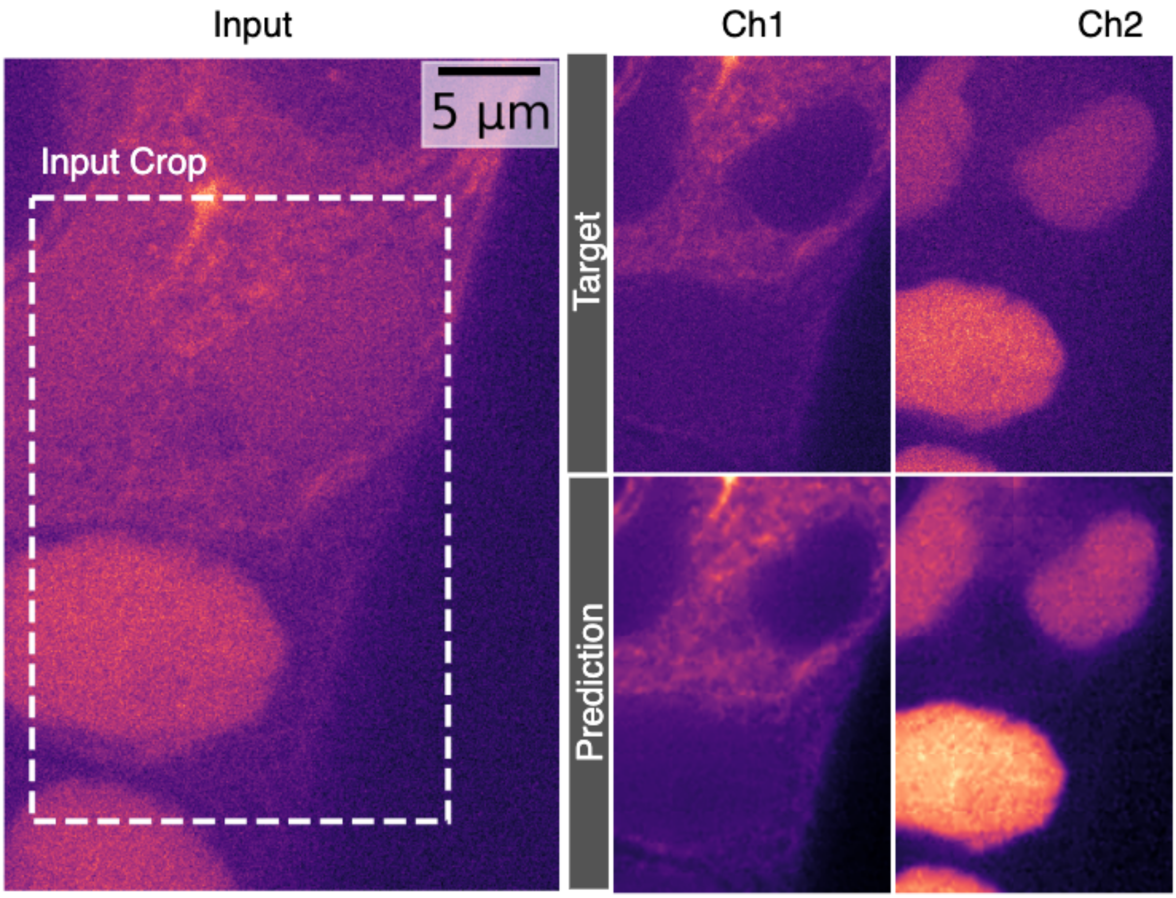


**Fig. S29** Qualitative Evaluation for Task V from Pavia-P24 dataset. Note that we show the target and the prediction corresponding to the input crop which is denoted in *Input* panel by a white dotted rectangle.

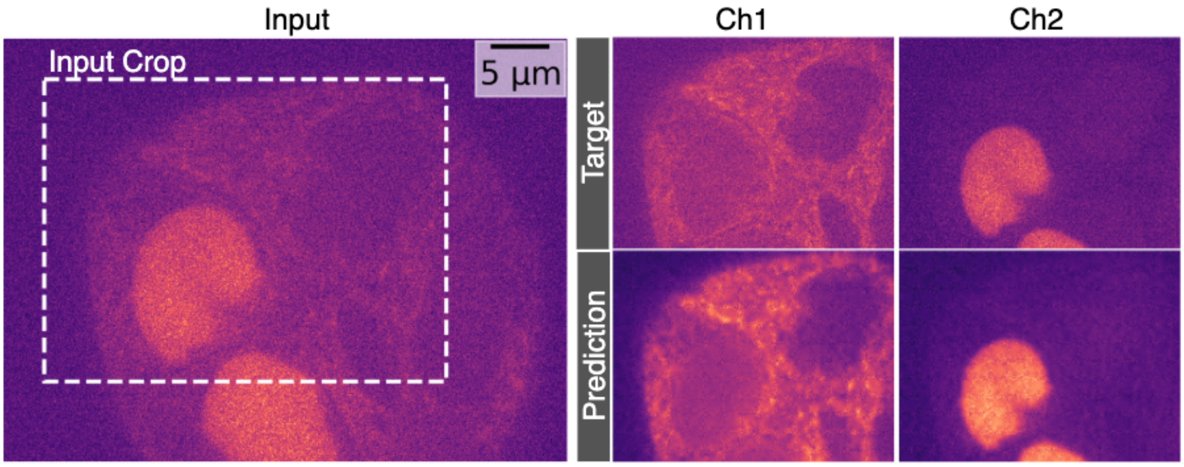


**Fig. S30** Qualitative Evaluation for Task VII from Pavia-P24 dataset. Note that we show the target and the prediction corresponding to the input crop which is denoted in *Input* panel by a white dotted rectangle.

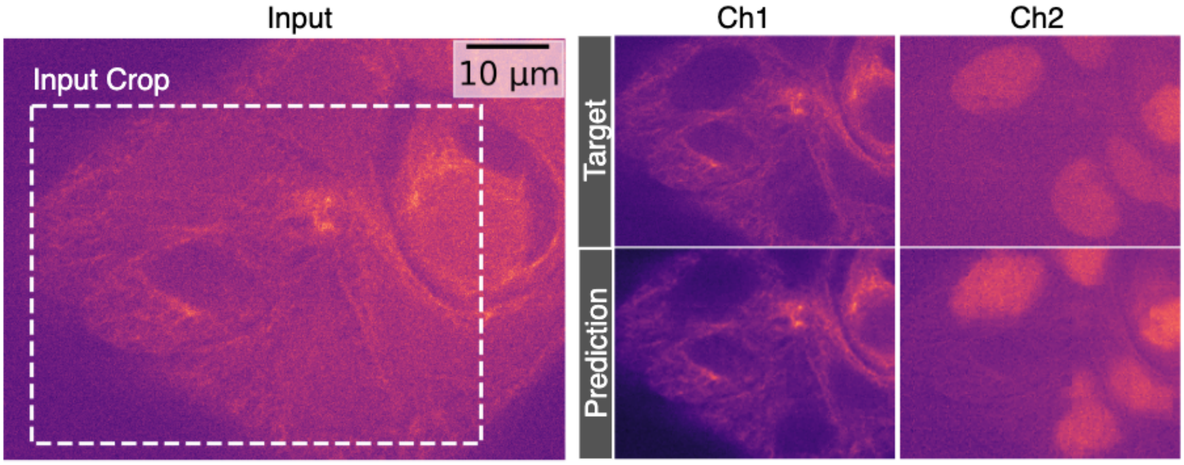


**Fig. S31** Qualitative Evaluation for Task VIII from Pavia-P24 dataset. Note that we show the target and the prediction corresponding to the input crop which is denoted in *Input* panel by a white dotted rectangle.

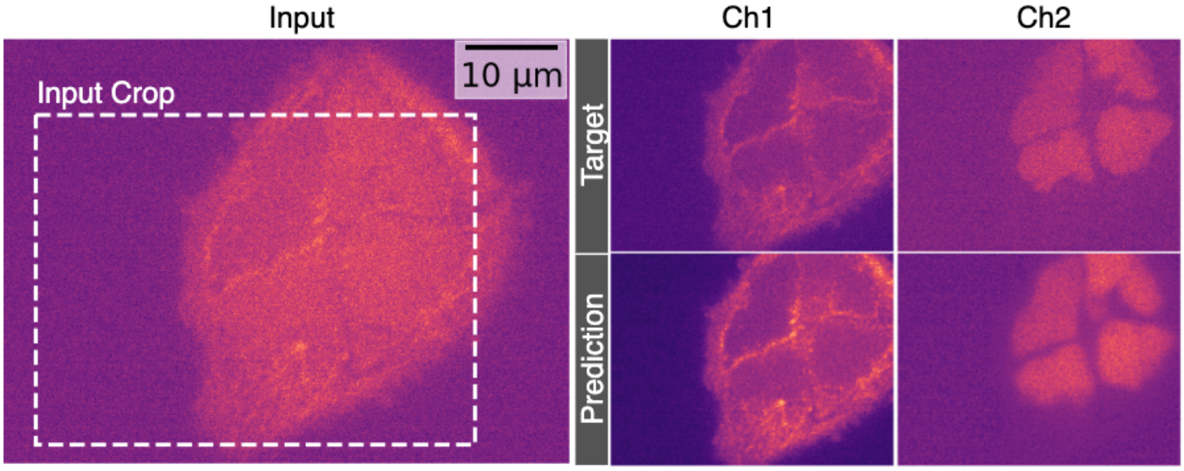


**Fig. S32** Qualitative Evaluation for Task IX from Pavia-P24 dataset. Note that we show the target and the prediction corresponding to the input crop which is denoted in *Input* panel by a white dotted rectangle.

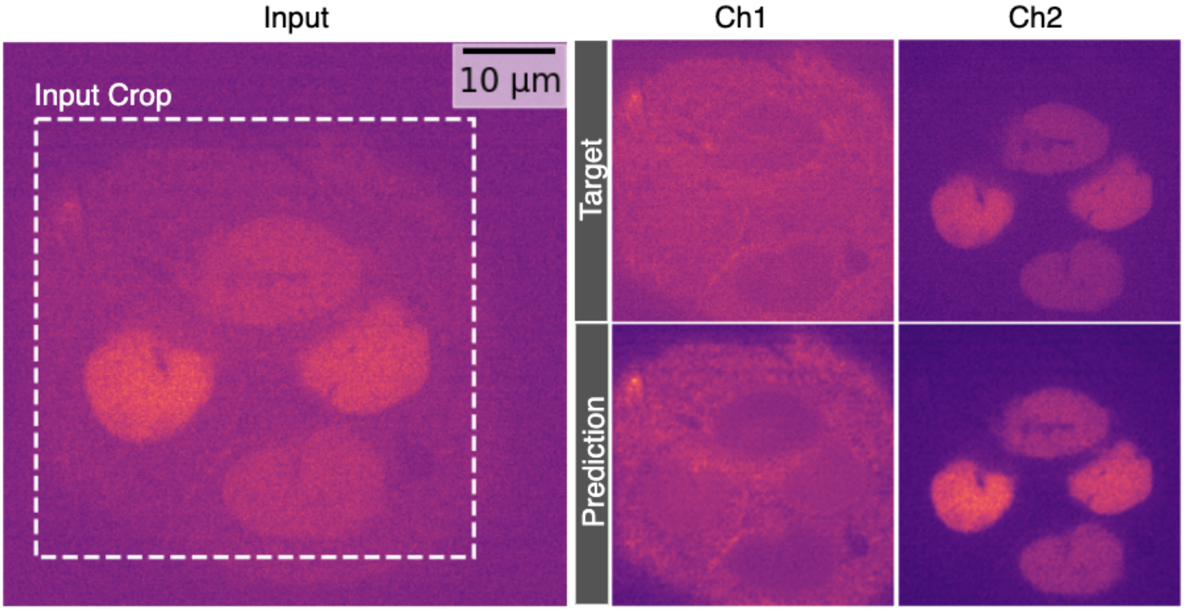


**Fig. S33** Qualitative Evaluation for Task X from Pavia-P24 dataset. Note that we show the target and the prediction corresponding to the input crop which is denoted in *Input* panel by a white dotted rectangle.

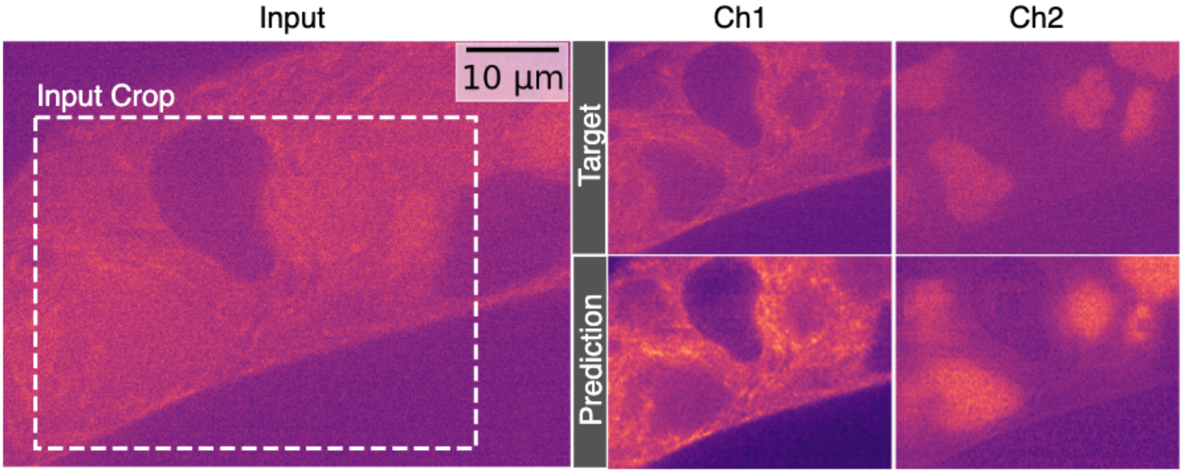


**Fig. S34** Qualitative Evaluation for Task XI from Pavia-P24 dataset. Note that we show the target and the prediction corresponding to the input crop which is denoted in *Input* panel by a white dotted rectangle.

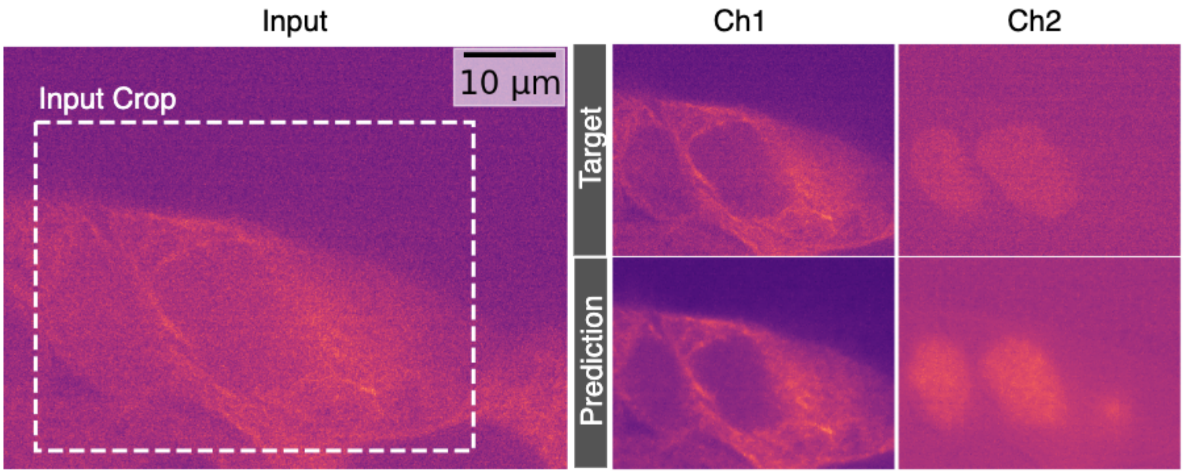


**Fig. S35** Qualitative Evaluation for Task XII from Pavia-P24 dataset. Note that we show the target and the prediction corresponding to the input crop which is denoted in *Input* panel by a white dotted rectangle.

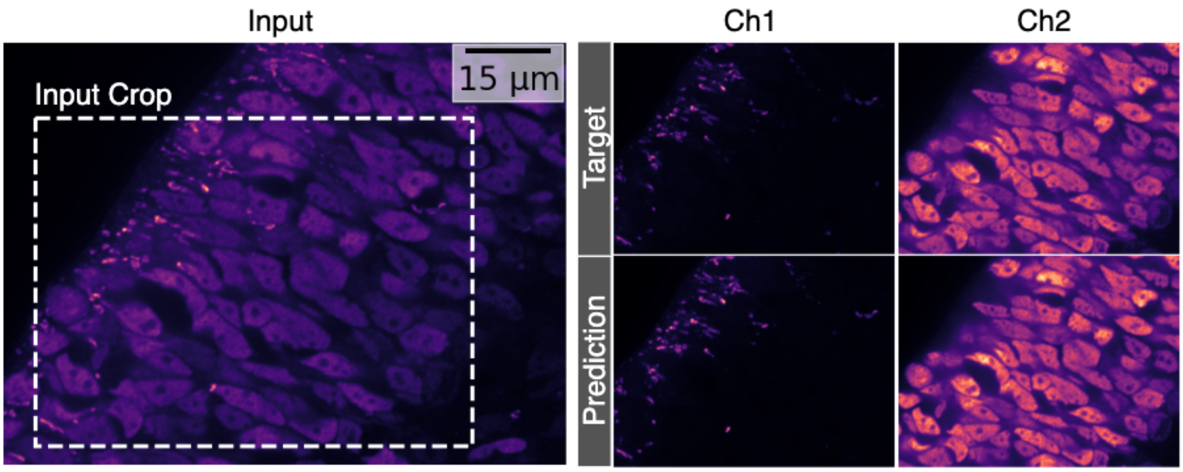


**Fig. S36** Qualitative Evaluation for Task XIII from HT-T24 dataset. Note that we show the target and the prediction corresponding to the input crop which is denoted in *Input* panel by a white dotted rectangle.

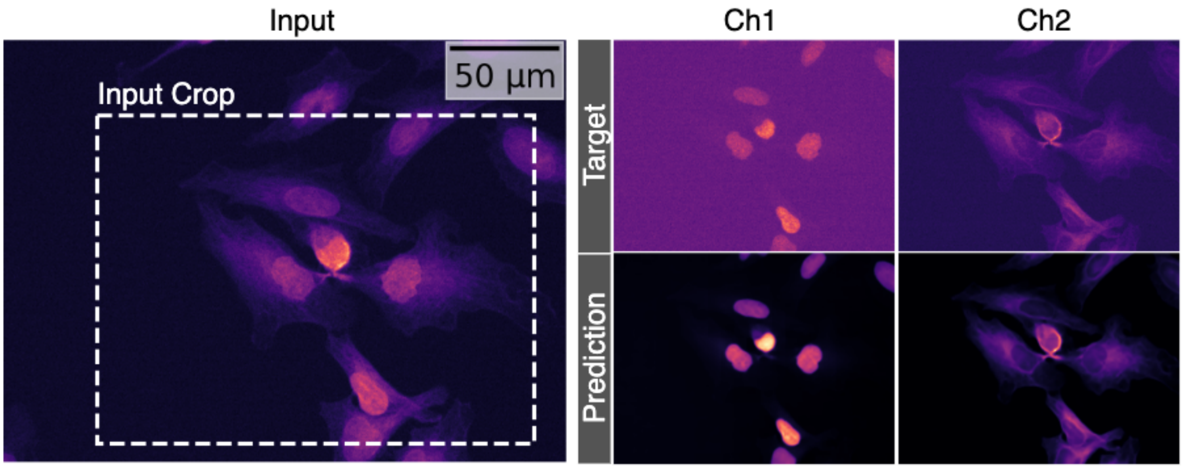


**Fig. S37** Qualitative Evaluation for Task XIV from HT-LIF24 dataset. Note that we show the target and the prediction corresponding to the input crop which is denoted in *Input* panel by a white dotted rectangle.

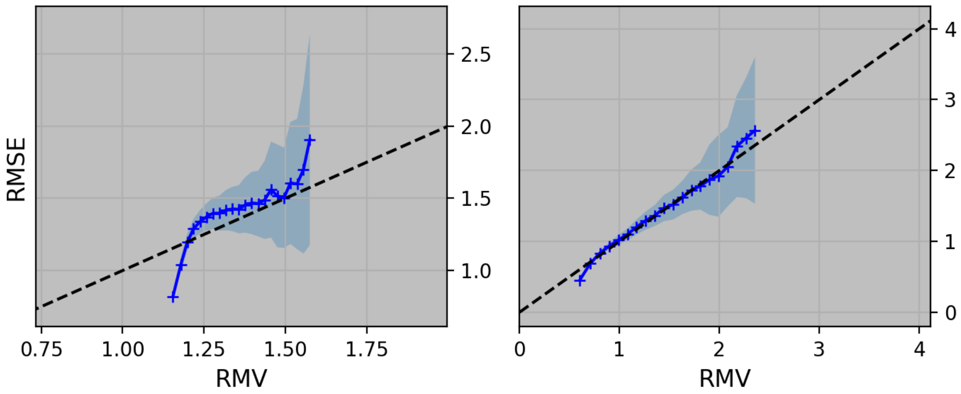


**Fig. S38** Calibration plot for Task XIV from Dataset HT-LIF24

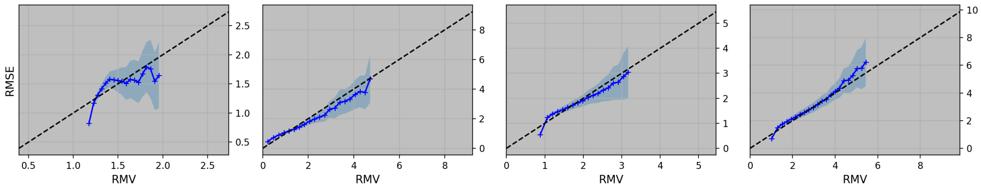


**Fig. S39** Calibration plot for Task XXVIII from Dataset HT-LIF24

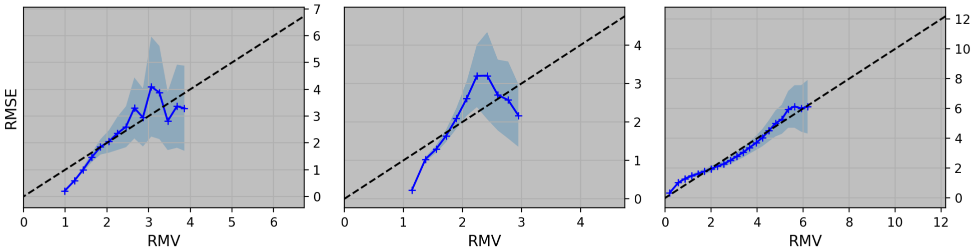


**Fig. S40** Calibration plot for Task XXVI from Dataset HT-LIF24

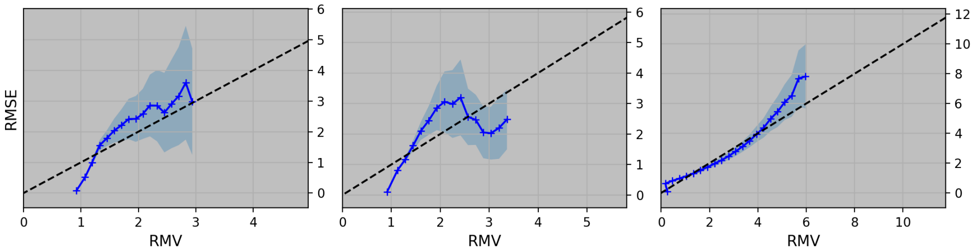


**Fig. S41** Calibration plot for Task XXVII from Dataset HT-LIF24

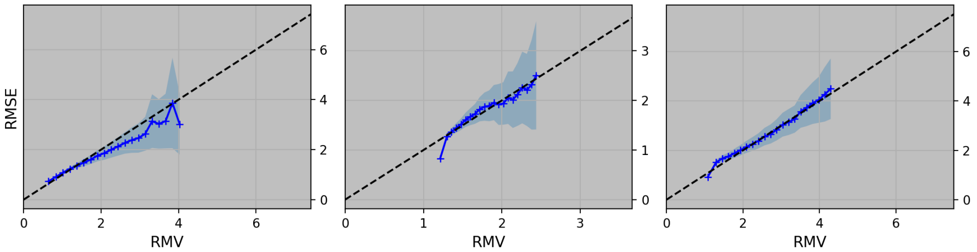


**Fig. S42** Calibration plot for Task XXIII from Dataset HT-LIF24 2ms

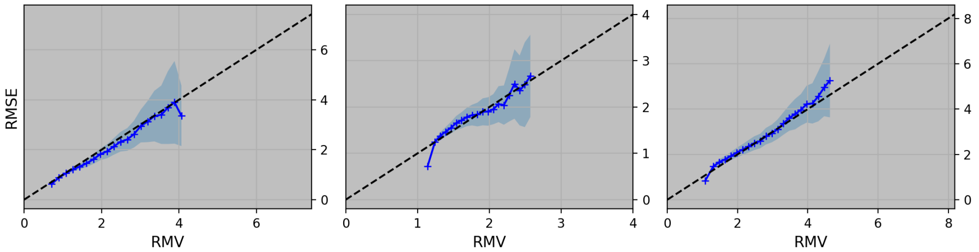


**Fig. S43** Calibration plot for Task XXIV from Dataset HT-LIF24

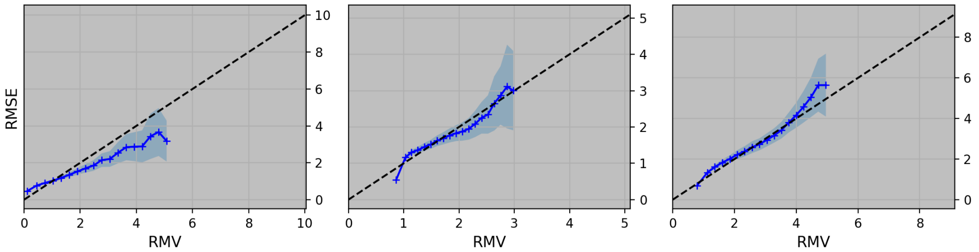


**Fig. S44** Calibration plot for Task XXV from Dataset HT-LIF24

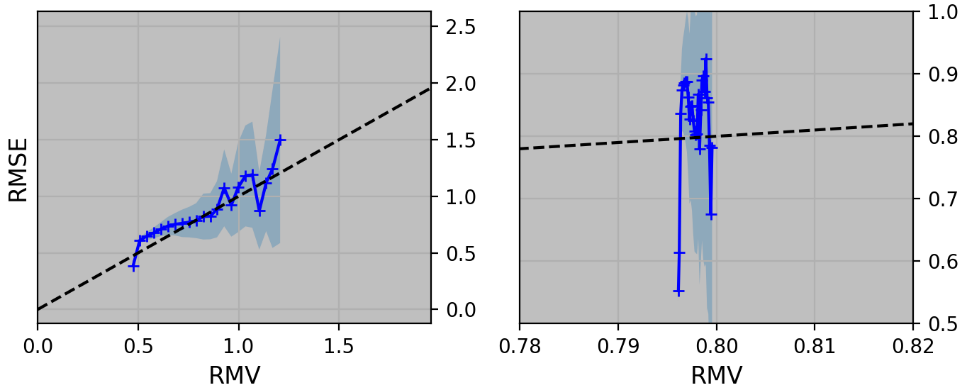


**Fig. S45** Calibration plot for Task V from Dataset Pavia-P24, mediumskew, high

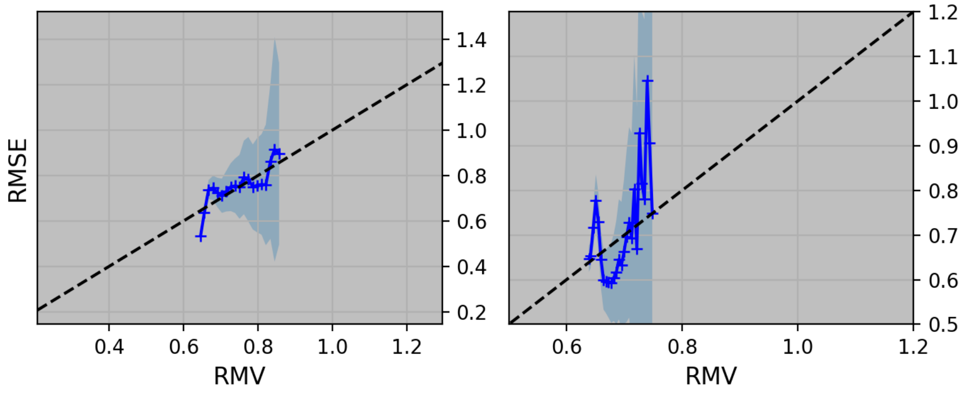


**Fig. S46** Calibration plot for Task VII from Dataset Pavia-P24, balanced, medium

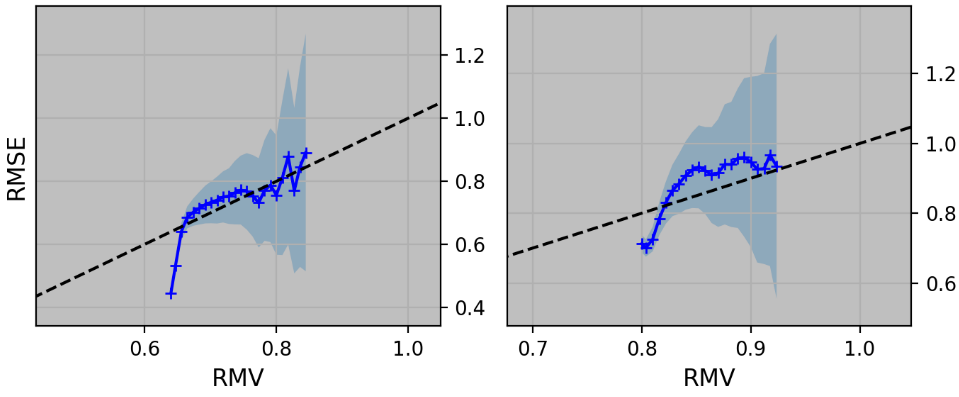


**Fig. S47** **Calibration plot for Task VIII from Dataset Pavia-P24, mediumskew, medium**

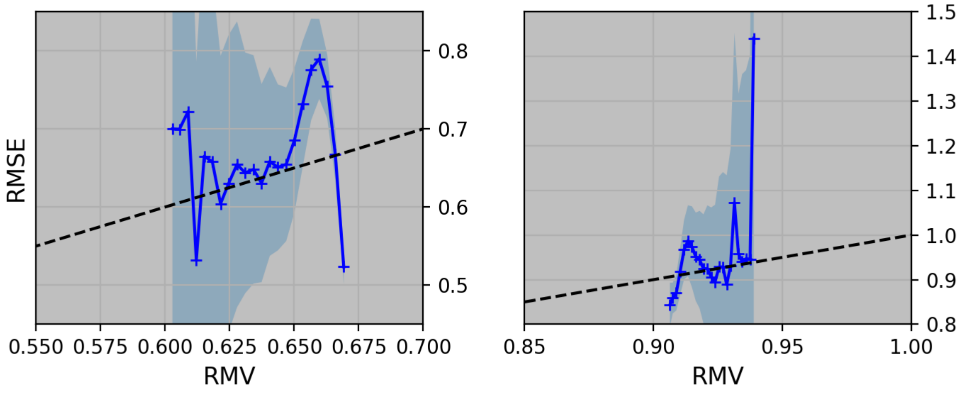


**Fig. S48** **Calibration plot for Task IX from Dataset Pavia-P24, highskew medium**

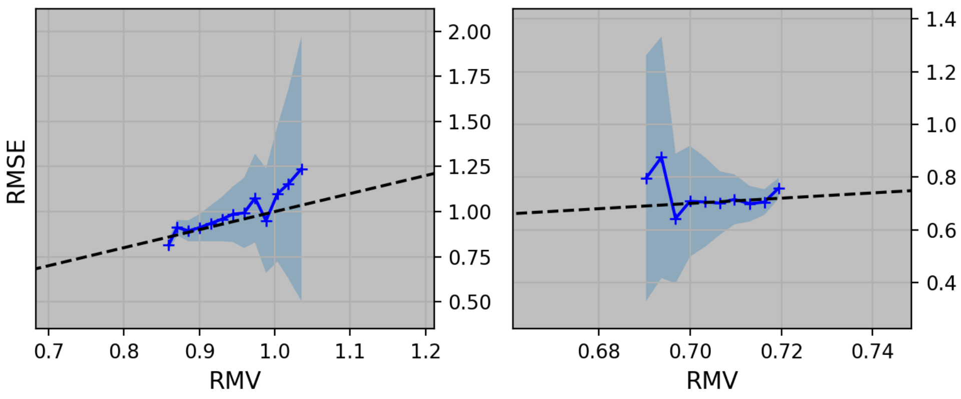


**Fig. S49** **Calibration plot for Task X from Dataset Pavia-P24, balanced low**

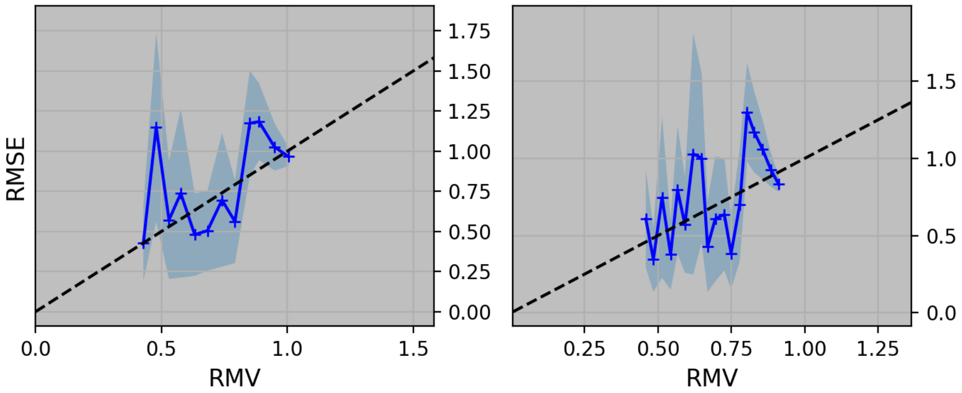


**Fig. S50** **Calibration plot for Task XI from Dataset Pavia-P24, mediumskew, low**

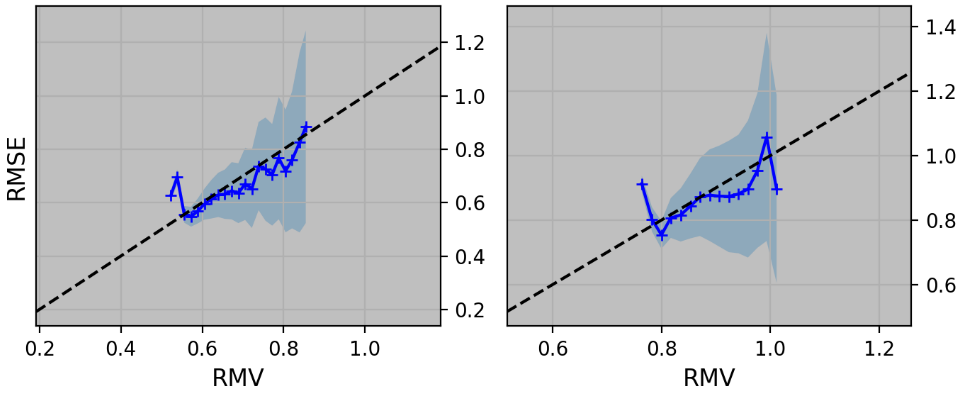


**Fig. S51** **Calibration plot for Task XII from Dataset Pavia-P24, highskew, low**

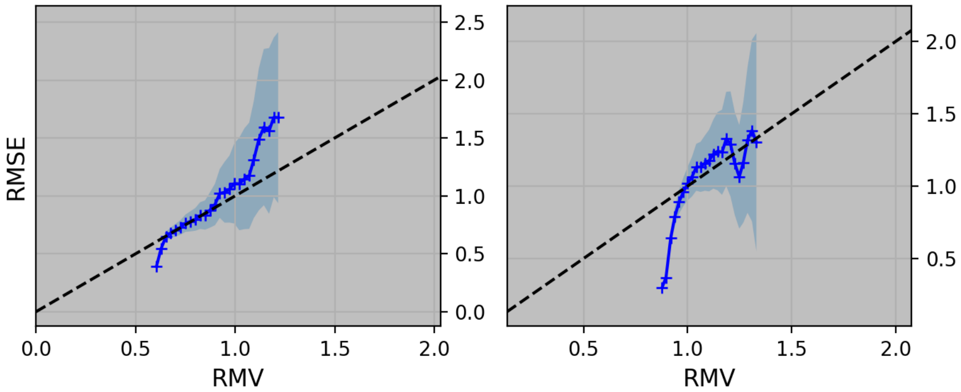


**Fig. S52** **Calibration plot for Task XX from Dataset Pavia-P24, balanced, high**

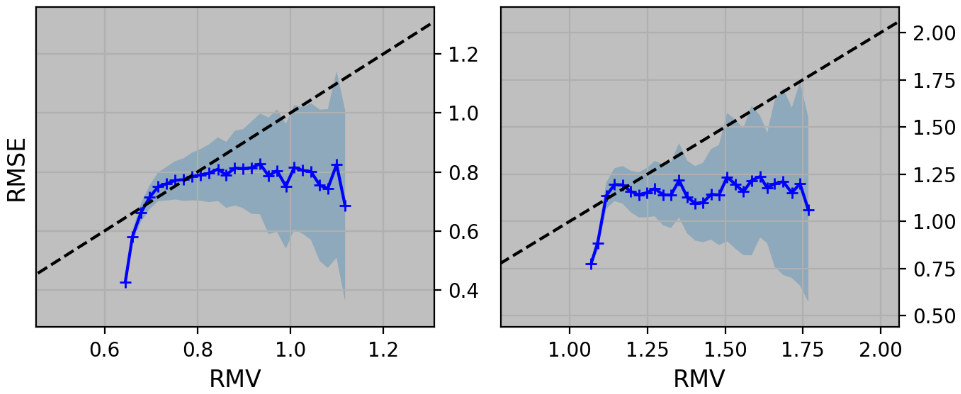


**Fig. S53** **Calibration plot for Task IV from Dataset Pavia-P24, highskew, high**

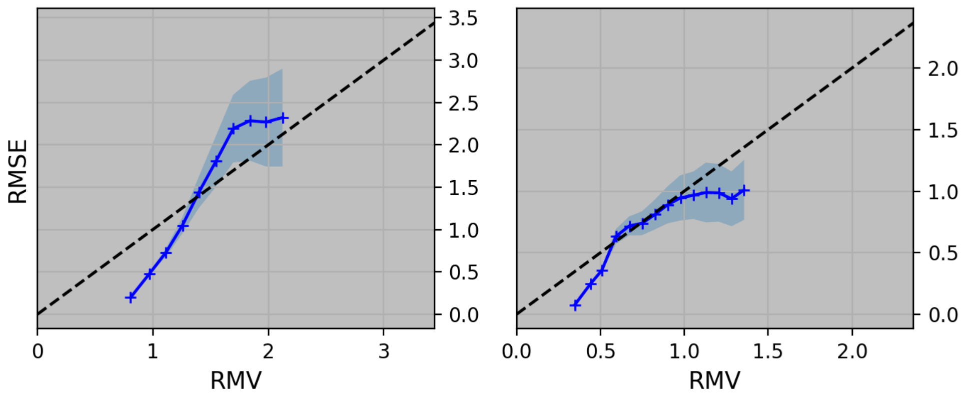


**Fig. S54 Calibration plot for Task I from Dataset HT-H24**

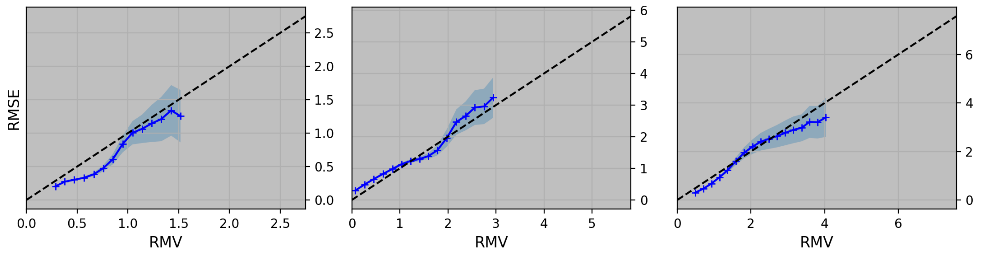


**Fig. S55 Calibration plot for Task XXI from Dataset CBZ-Z18**

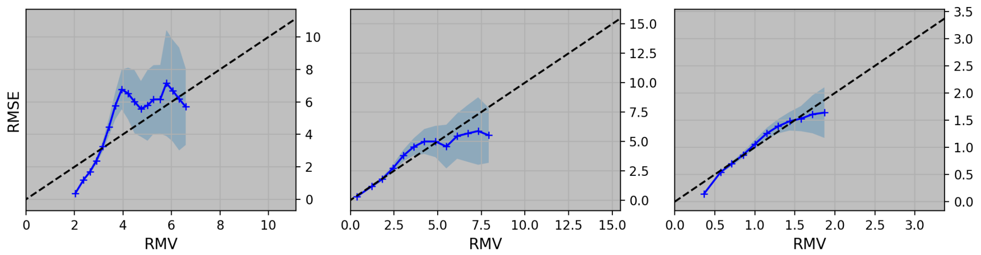


**Fig. S56 Calibration plot for Task XXII from Dataset CBZ-N18**

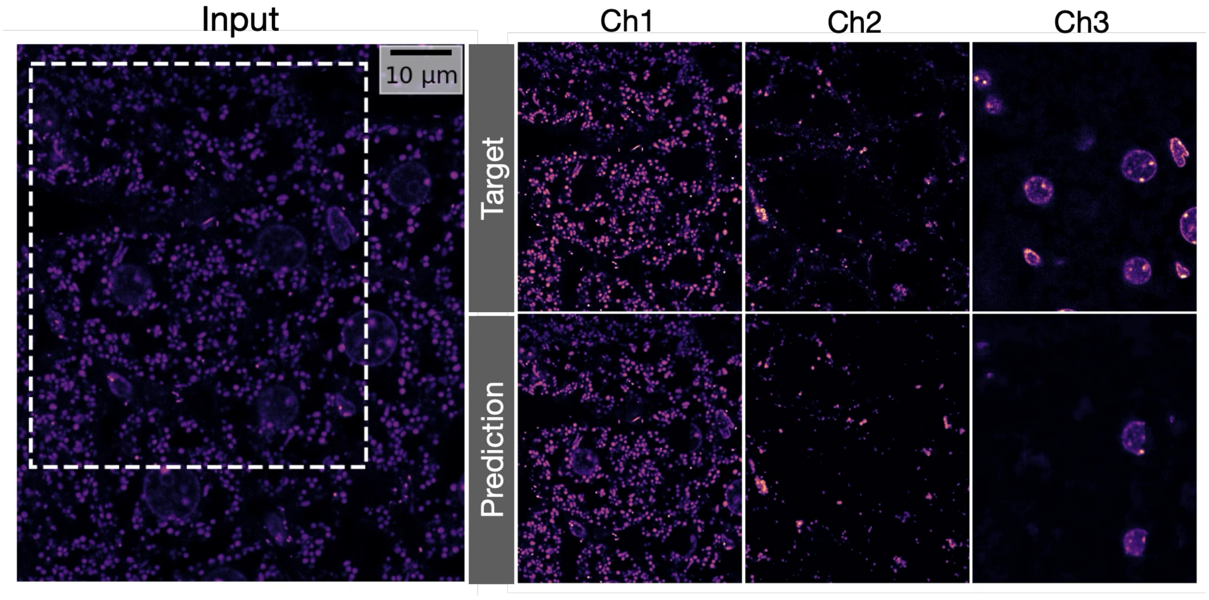


**Fig. S57** Qualitative Evaluation for Task XXIII from HHMI-D25$_{8bit}$ dataset. Note that we show the target and the prediction corresponding to the input crop which is denoted in *Input* panel by a white dotted rectangle. Also note that the predictions for channel 3 are of rather poor quality and that you can find a description of how this problem was solved in the Supplementary Section B.

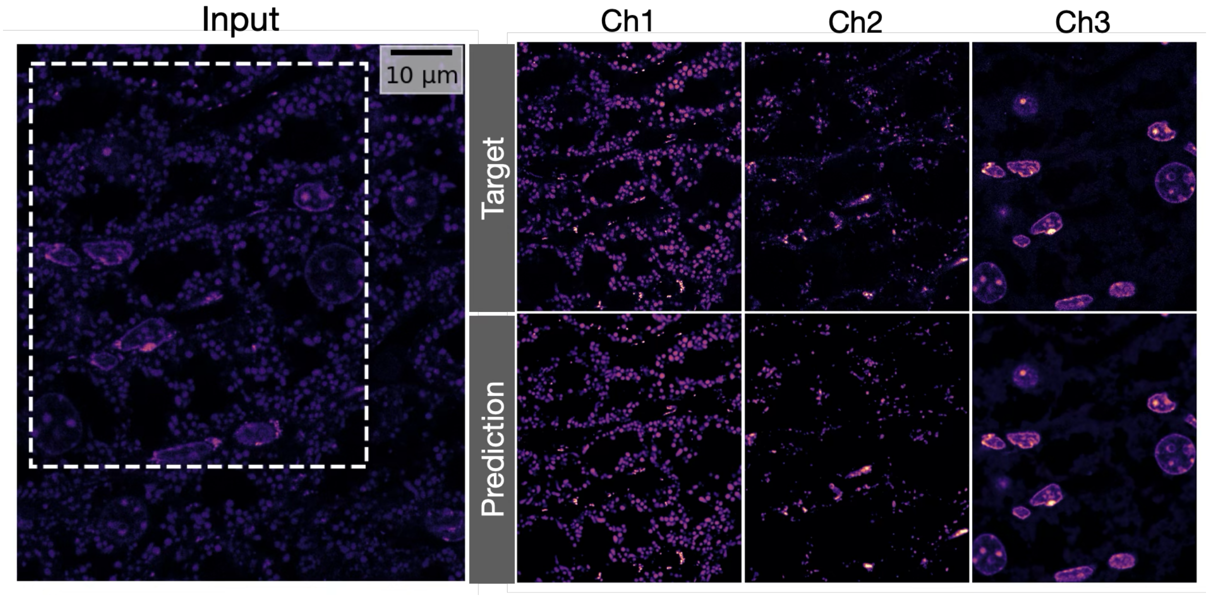


**Fig. S58** Qualitative Evaluation for Task XXXIII from HHMI-D25$_{16bit}$ dataset. Note that we show the target and the prediction corresponding to the input crop which is denoted in *Input* panel by a white dotted rectangle.

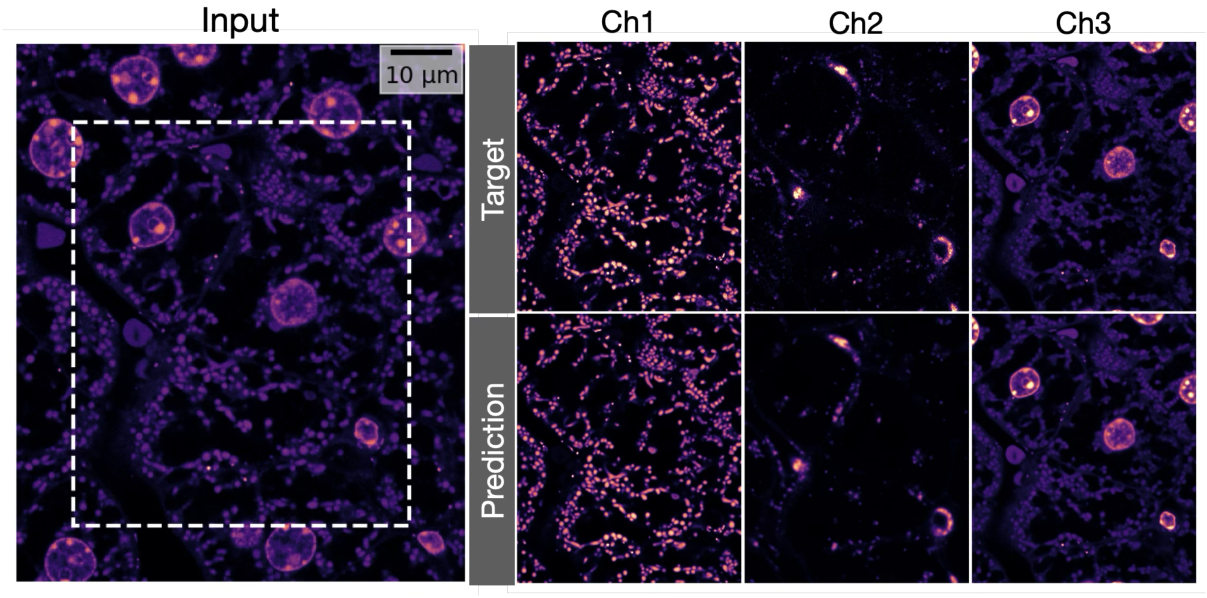


**Fig. S59** Qualitative Evaluation for Task XXXVI from HHMI-D25$_{16bit,0.25}$ dataset. Note that we show the target and the prediction corresponding to the input crop which is denoted in *Input* panel by a white dotted rectangle.

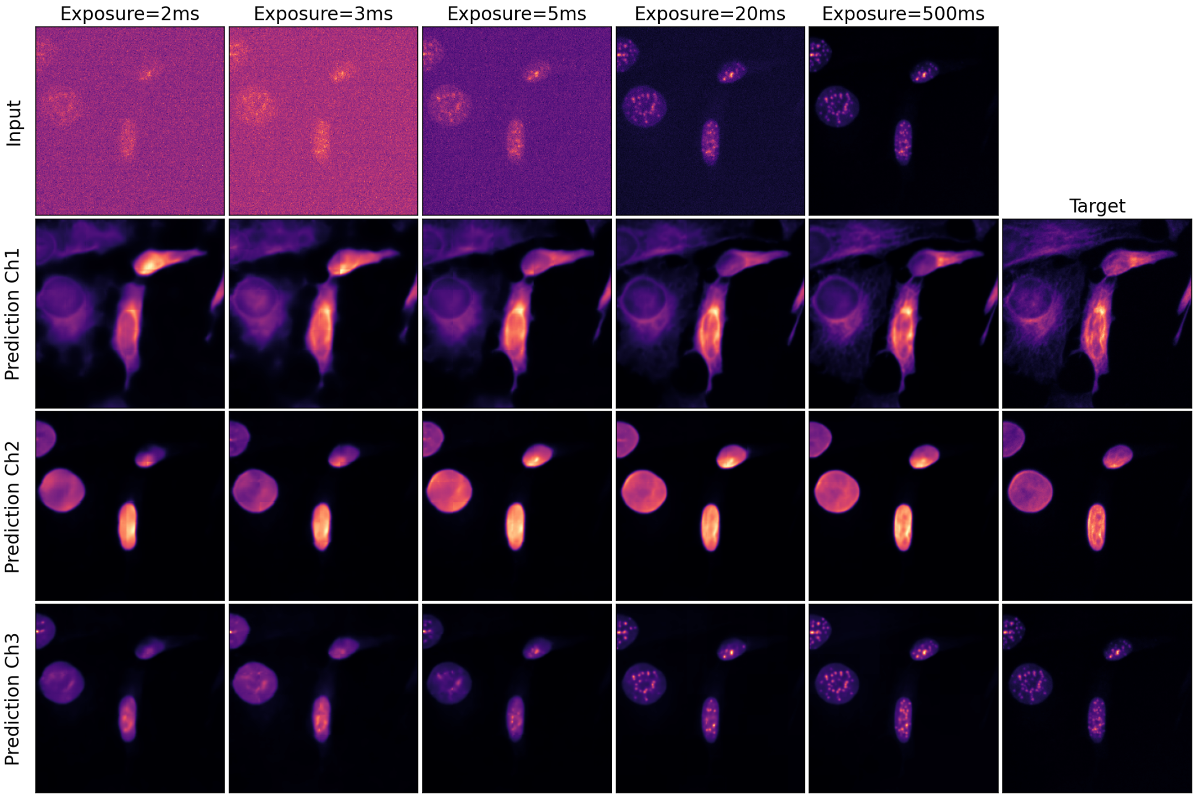


**Fig. S60 Effect of SNR on Model Performance (HT-LIF24 Dataset):** We evaluate how SNR influences model performance using the HT-LIF24 dataset. Different models are trained on data subsets acquired with varying exposure durations—leading to different SNRs—and their predictions are compared over a common region of interest. High-frequency details in the predictions (especially the third channel) are visibly reduced when the input SNR is lower.

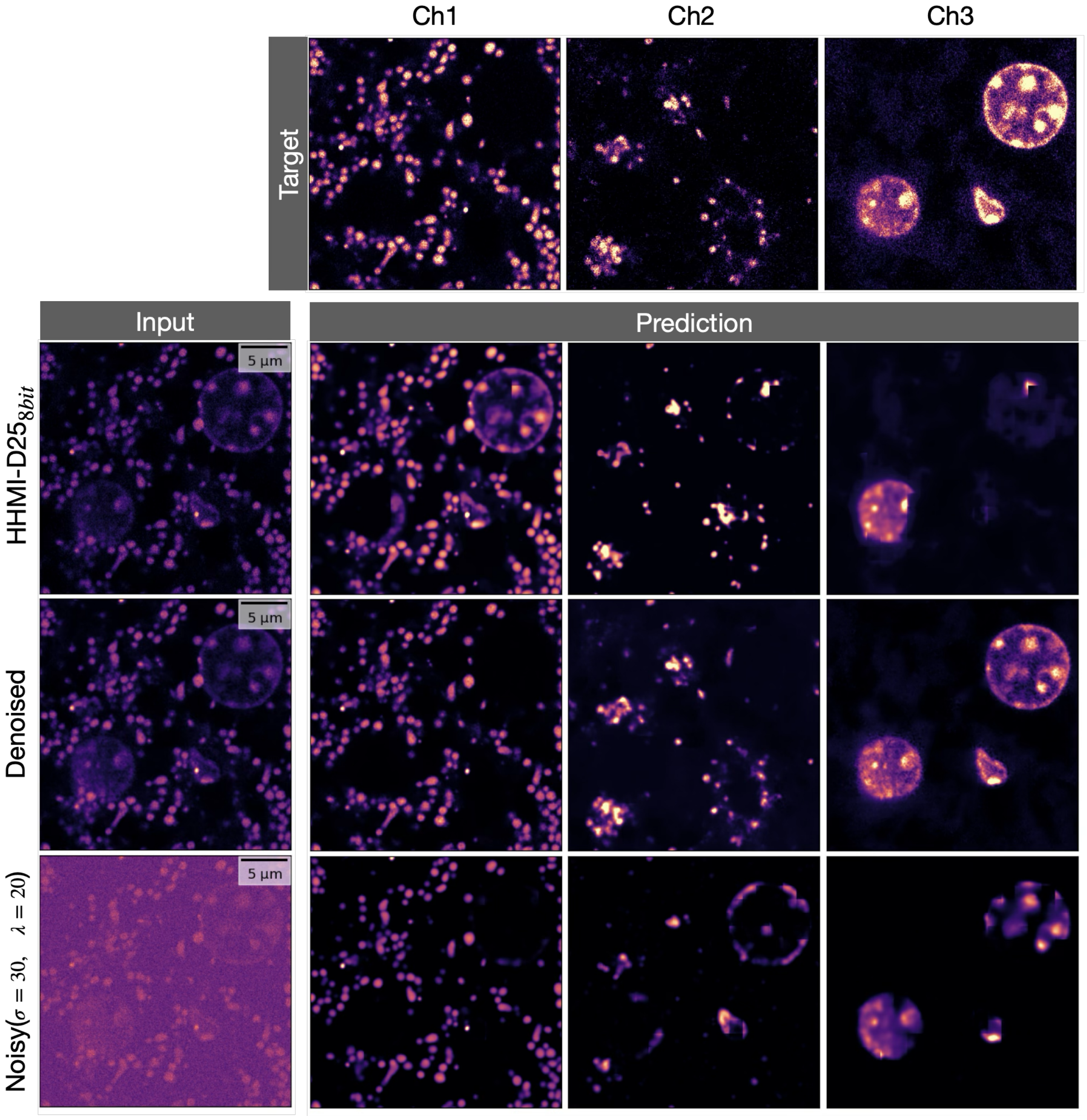


**Fig. S61 Effect of SNR on Model Performance (HHMI-D25$_{8bit}$ Dataset):** We assess the impact of SNR on a subset of the HHMI-D25 dataset. Comparing the predictions (row 2, results of Task XXIII) with the ground truth (row 1), we observe that the prediction quality, particularly for the third channel, is not good, with entire parts of the structures being put into the other channels. We then train Micro𝕊plit using a Noise2Void [4] denoised version of the same data, leading to much improved semantic unmixing performance (row 3, Task XXXI). Finally, we re-introduced synthetic Gaussian and Poisson noise to the denoised HHMI-D25 data used in Task XXXI and retrained Micro𝕊plit on this lower-SNR data (row 4, Task XXXII). As it was likely to be expected, this does again drop the semantic unmixing performance.

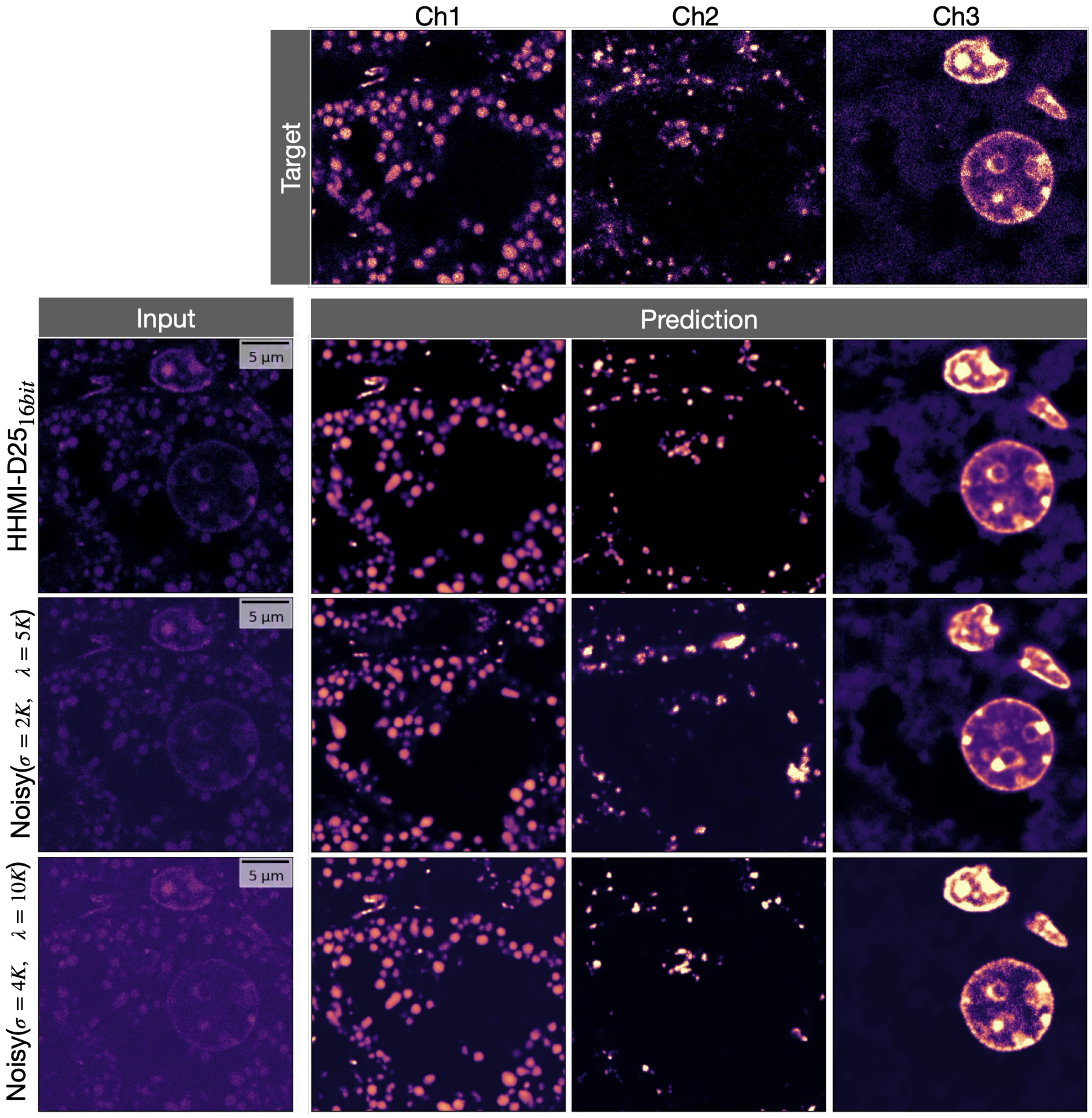


**Fig. S62 Effect of SNR on Model Performance (HHMI-D25$_{16bit}$ Dataset):** We assess the impact of SNR on the HHMI-D25$_{16bit}$ dataset. For this part of the HHMI-D25 data, the predictions (row 2, Task XXXIII) are visually more close to the target images compared to results on HHMI-D25$_{8bit}$ dataset (Figure S61, row 2, Task XXIII). We then introduce two levels of additional Gaussian and Poisson noise to the HHMI-D25$_{16bit}$ data to reduce SNR and retrain Micro𝕊plit on those noisier versions of the data (row 3 and 4, Tasks XXXIV and XXXV, respectively). As it was likely to be expected, the reduced SNR leads to a noticeable decline in the semantic unmixing performance.
